# Insulin-mediated endothelin signaling is antiviral during West Nile virus infection

**DOI:** 10.1101/2023.01.17.524426

**Authors:** Chasity E. Trammell, Evelyn H. Rowe, Brianne J. Jones, Aditya B. Char, Stephen Fawcett, Laura R.H. Ahlers, Alan G. Goodman

## Abstract

West Nile virus (WNV) is the most prevalent mosquito-borne virus in the United States with approximately 2,000 cases each year. There are currently no approved human vaccines and a lack of prophylactic and therapeutic treatments. Understanding host responses to infection may reveal potential intervention targets to reduce virus replication and disease progression. The use of *Drosophila melanogaster* as a model organism to understand innate immunity and host antiviral responses is well established. Previous studies revealed that insulin-mediated signaling regulates WNV infection in invertebrates by regulating canonical antiviral pathways. Because insulin signaling is well-conserved across insect and mammalian species, we sought to determine if results using *D. melanogaster* can be extrapolated for the analysis of orthologous pathways in humans. Here, we identify insulin-mediated endothelin signaling using the *D. melanogaster* model and evaluate an orthologous pathway in human cells during WNV infection. We demonstrate that endothelin signaling reduces WNV replication through the activation of canonical antiviral signaling. Taken together, our findings show that endothelin-mediated antiviral immunity is broadly conserved across species and reduces replication of viruses that can cause severe human disease.

**IMPORTANCE:** Arboviruses, particularly those transmitted by mosquitoes, pose a significant threat to humans and are an increasing concern because of climate change, human activity, and expanding vector-competent populations. West Nile virus is of significant concern as the most frequent mosquito-borne disease transmitted annually within the continental United States. Here, we identify a previously uncharacterized signaling pathway that impacts West Nile virus infection, namely endothelin signaling. Additionally, we demonstrate that we can successfully translate results obtained from *D. melanogaster* into the more relevant human system. Our results add to the growing field of insulin-mediated antiviral immunity and identifies potential biomarkers or intervention targets to better address West Nile virus infection and severe disease.

## INTRODUCTION

West Nile virus (WNV) is a member of the family *Flaviviridae* and is transmitted predominately between *Culex quinquefasciatus* and birds with humans as incidental “dead-end” hosts (1). WNV was introduced to the Western Hemisphere in New York in 1999 and has since become endemic in the United States (2–4). Like other arthropod-borne viruses, WNV poses a significant health threat due to the expansion of mosquito range and activity (5–7) without effective means to address these concerns at a transmission or clinical level. While our ability to intervene arboviral exposure at the vector-transmission level has progressed significantly in the past decade through genetic (8), microbial (9), or small molecule (10) targeting of mosquito responses, addressing WNV clinical cases has lagged. There are currently no vaccines or specific treatments available for treating WNV with the best approaches being disease management and pain relief (11).

*Drosophila melanogaster* is an established model organism that has been used for studying host responses. This is due to its readily accessible and annotated genome that permits broad or targeted study of specific signaling pathways or interactions. *D. melanogaster* has been successfully used to study innate immune responses to flavivirus infection including WNV (12, 13) and Zika virus (ZIKV) (14). Previous investigation identified insulin-mediated induction of JAK/STAT as a critical component of host survival and immunity to WNV in *D. melanogaster* that was conserved in *Culex quinquefasciatus* (13). Because of the broad conservation that the insulin signaling pathway is across species, especially from *D. melanogaster* to human systems (15, 16), we rationalize that insulin-mediated antiviral immunity may exist in the human innate immune system as well.

Previous studies have shown that viral infection may target components of insulin signaling that can result in insulin resistance and dysfunction (17–21), but there is limited investigation about how this host-virus interaction can be a potential intervention target. Because of the substantial number of downstream signaling pathways insulin signaling impacts, we sought to identify previously unidentified signaling pathways that canonical insulin signaling regulates and may have important roles in the host response to viral infection. In addition, because of the significant conservation that insulin signaling possesses across species and the genetic power of the *D. melanogaster* model, we propose that we can extrapolate identified pathways from *D. melanogaster* and their orthologous pathways in the human system.

In this study, we performed RNA sequencing (RNAseq) in *D. melanogaster* during WNV infection to identify novel antiviral response elements that are activated in the presence of insulin. We find that insulin induces both canonical antiviral response elements, as well as genes that are were previously unidentified components of host immunity. Specifically, we identified the endothelin signaling pathway and evaluated its importance for host survival and reducing WNV infection. Endothelin signaling is primarily associated in vasoconstriction and cardiovascular function (22) but has been suggested as a biomarker for various infectious disease pathogenesis (23–25), immune dysregulation (26, 27), and insulin sensitivity (28, 29). We then used this information to evaluate endothelin signaling in human cells. We similarly found that endothelin signaling was important for regulating viral replication and regulating insulin-mediated responses to infection against both attenuated and virulent WNV strains. These results suggest that insulin regulates endothelin signaling such that the loss of endothelin results in deficient

host antiviral immunity. These pathways are conserved across species and may be a potential avenue for future therapeutic research.

## RESULTS

### *Transcriptomic profiling of* D. melanogaster *S2 cells identifies antiviral pathways linked to insulin-signaling*

We first sought to generate a complete transcript profile of *D. melanogaster* S2 cells following 24 h treatment with 1.7 μM bovine insulin and 8 h infection with WNV-Kun (MOI 1 PFU/cell). Gene expression in treated and/or infected cells was measured relative to that in controls receiving neither bovine insulin or virus (Fig. 1A). These experimental conditions were selected based on previous data showing that bovine insulin treatment induces sufficient insulin and JAK/STAT signaling in S2 cells (13). The average number of sequence reads mapped to the *D. melanogaster* genome is approximately 93.22% (Table S1, Sheet 1).

**Figure 1:**
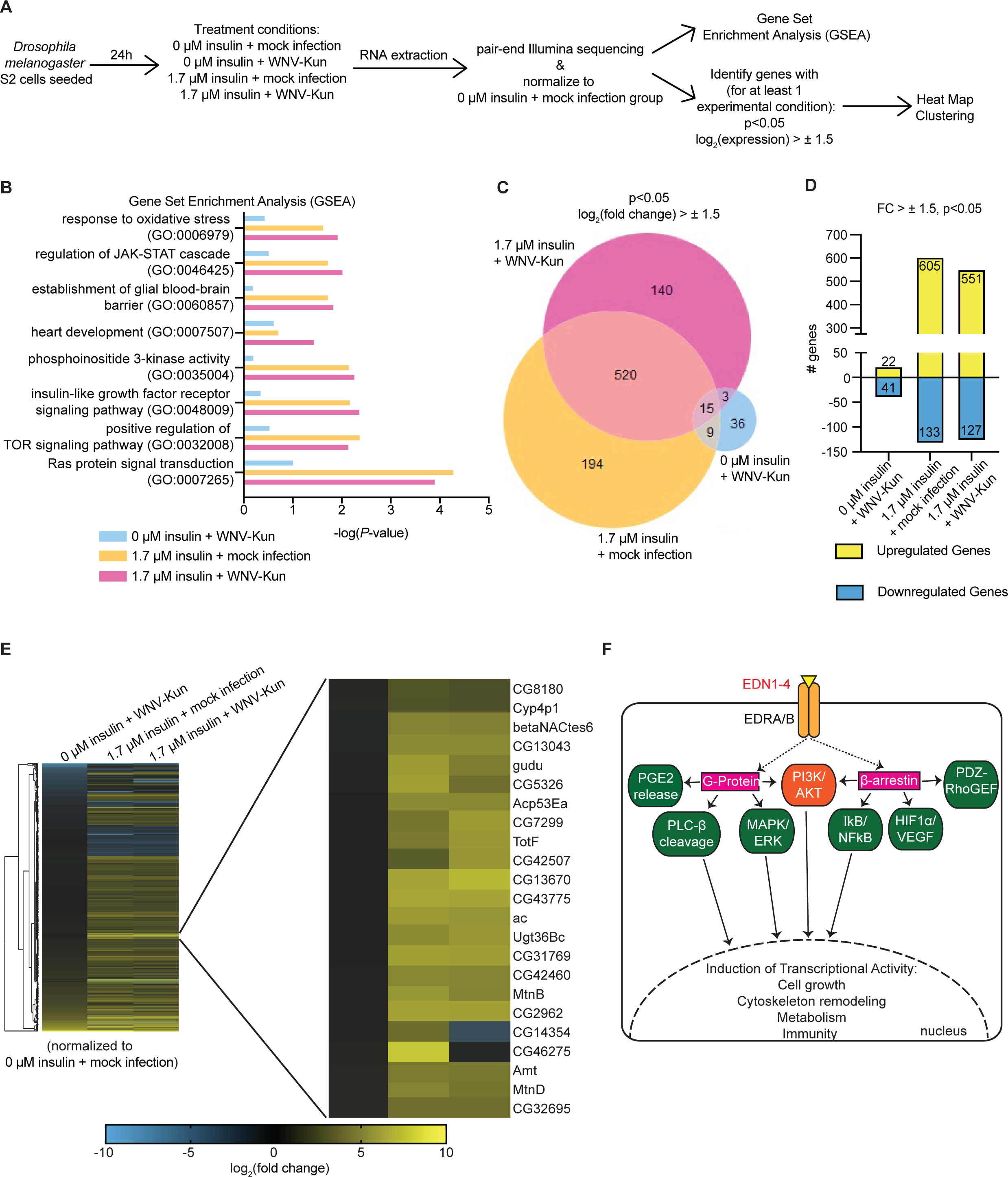
Insulin treatment during WNV-Kun infection in *D. melanogaster* S2 cells induces canonical and previously unidentified signaling pathways. (A) Schematic illustrating experimental design for RNA extraction and RNA sequencing (RNAseq) analysis of *D. melanogaster* S2 cells with or without insulin treatment or WNV-Kun infection. (B) Gene Set Enrichment Analysis (GSEA) using transcript levels for each experimental condition normalized to 0 μM insulin + mock infection from the RNAseq analysis. Gene ontology (GO) categories were selected based on GSEA p value (p < 0.05) for at least one experimental condition. (C) Venn Diagram of all transcripts enriched or suppressed for each experimental condition normalized to 0 μM insulin + mock infection. Transcripts were selected based on their log_2_(fold change) (FC) > ± 1.5 and p < 0.05 for at least one experimental condition. (D) The number of genes transcriptionally enriched (yellow) or suppressed (blue) for each experimental condition normalized to 0 μM insulin + mock infection. Hierarchal clustering and heat map expression of genes transcriptionally enriched or suppressed as identified in (C-D). Genes shown in enlarged cluster identify a subset of genes that showed the most up-regulation compared to no insulin treatment. GO analysis identifies this get set associated with endothelin signaling. (F) Schematic of canonical endothelin signaling in mammals and its intracellular and transcriptional activity.

Gene set enrichment analysis (GSEA) was performed to identify and compare enriched gene sets in 0 μM insulin + WNV-Kun, 1.7 μM insulin + mock infection, and 1.7 μM insulin + WNV-Kun (Fig. 1B). Analysis was completed to identify previously unidentified gene sets for further analysis as well to compare to previous, targeted qRT-PCR analysis showing enrichment of insulin and JAK/STAT signaling (13). Gene sets were filtered for p < 0.05 in at least one experimental condition and were selected based on their association with immunity and WNV disease (Table S2). We identified eight gene sets that are significantly enriched in the presence of insulin including immune response elements (response to oxidative stress, regulation of JAK-STAT cascade), canonical insulin signaling (phosphoinositide 3-kinase activity, insulin-like growth factor receptor signaling pathway, positive regulation of TOR signaling pathway, Ras protein signal transduction), and physiological development (establishment of glial blood-brain barrier, heart development). (Fig. 1B).

Further analysis into the specific genes that were transcriptionally induced or suppressed was carried out to better understand the impact that infection or insulin treatment have on the *D. melanogaster* transcriptome. Genes were filtered for p < 0.05 and a log_2_(fold change) > ± 1.5 for at least one experimental condition. There was a ∼10-fold increase in the number of differentially expressed genes in cells that received insulin treatment and those that received no insulin (Fig. 1 C-D). 535 genes were commonly regulated in the presence of insulin regardless of WNV-Kun infection status (Fig. 1C). Together, this suggests that insulin treatment enriches or suppresses transcriptional activity with a high overlap in genes affected between mock infection and WNV-Kun infection. Cells that were not treated with insulin but were infected with WNV-Kun only exhibited 22 upregulated genes and 41 downregulated genes. Cells that received only insulin treatment reported 605 upregulated genes and 133 downregulated genes. Cells that received insulin treatment and WNV-Kun infection exhibited 551 upregulated genes and 127 downregulated genes (Fig. 1D). These results suggest that insulin-treatment regulates a large set of genes during early stages of infection that can potentially impact later virus-specific responses.

Genes that were transcriptionally altered in Fig. 1C-D were used to generate a hierarchical clustering heatmap (Fig. 1E). As the goal of this study was to investigate effectors involved in insulin-mediated antiviral immunity, we were specifically interested in identifying and evaluating gene clusters that were enriched in the presence of insulin treatment (Fig. 1E, expanded node). Genes identified within the selected cluster were then imported into PANTHER Classification System to identify gene ontology (GO) categories that were overrepresented (30, 31) (Table S1, Sheet 2). Using this gene set, only the endothelin signaling pathway was identified. Endothelin signaling is primarily associated with cardiovascular function and smooth muscle constriction (22, 32). Through this functional role, endothelin signaling also interacts and impacts associated components linked to insulin signaling including the PI3K/AKT/FOXO axis (28, 29, 33–37) and MAPK/ERK axis (38, 39) (Fig. 1F). The endothelin signaling pathway is not a canonical immune pathway; however, it has been linked to *Mycobacterium tuberculosis* (23) and Hepatitis B/C virus (HBV) (HCV) infection (24, 40) which leads us to consider that endothelin may also be involved during WNV infection and should be further analyzed. Further analysis of *D. melanogaster* genes associated with endothelin signaling outside the heatmap shows that insulin treatment + mock infection or insulin treatment + WNV-Kun infection cells had significant transcriptional activity compared to only WNV-Kun infected cells (Fig. S1). These endothelin-related genes were selected based on their PANTHER GO classification and designation (30, 31). Because of the lack of knowledge pertaining to endothelin signaling in the insect or in the context of WNV, we further investigated this pathway to determine if it may be a mediator of insulin-mediated antiviral immunity against WNV.

### D. melanogaster CG43775 *contributes to insulin-mediated antiviral immunity*

To validate and expand upon our RNAseq results, we more closely examined the magnitudes of fold changes presented in Fig. 1E. *CG43775* was one of the most up-regulated genes in the insulin treatment conditions found within the endothelin signaling-identified cluster in Fig. 1E. (Table S1, Sheet 3). Further analysis of this gene also identified a potential human ortholog, peptidase inhibitor 16 (41), which is associated with insulin (42, 43) and cardiovascular-related function (44, 45) similar to endothelin signaling (46). Based on this knowledge, we hypothesize that *CG43775* is an uncharacterized gene associated with insulin signaling and contributes to host immunity. We examined *CG43775* induction under the same conditions using qRT-PCR. We observed significant induction of *CG43775* in S2 cells with 1.7 μM insulin + WNV-Kun relative to other experimental conditions (Fig. 2A). We next experimented with adult flies that contained a transposable element insertion in *CG43775* to disrupt its expression (*CG43775^MB08418^)* compared to genetic control flies (*w^1118^*) (47, 48). Survival of female flies that received either 5,000 PFU/fly or a mock infection was measured over 30 days. A hazard ratio was generated as a metric of host mortality which compares the mortality rate of the virus-infected mutant flies to that of the virus-infected control flies (Fig. 2B). We observed significant mortality in *CG43775* mutant flies with a mortality rate approximately 7-times greater compared to the control flies. To expand the role that *CG43775* has on host survival to viral infection, we measured viral titer in mutant and control flies at 1-, 5-, and 10-days post-infection (d p.i.) by standard plaque assay (Fig. 2C). We observed significantly higher virus replication in mutant flies by 10 d. These data suggest that *CG43775* is important for host survival to WNV-Kun infection due to its ability to reduce virus replication.

**Figure 2:**
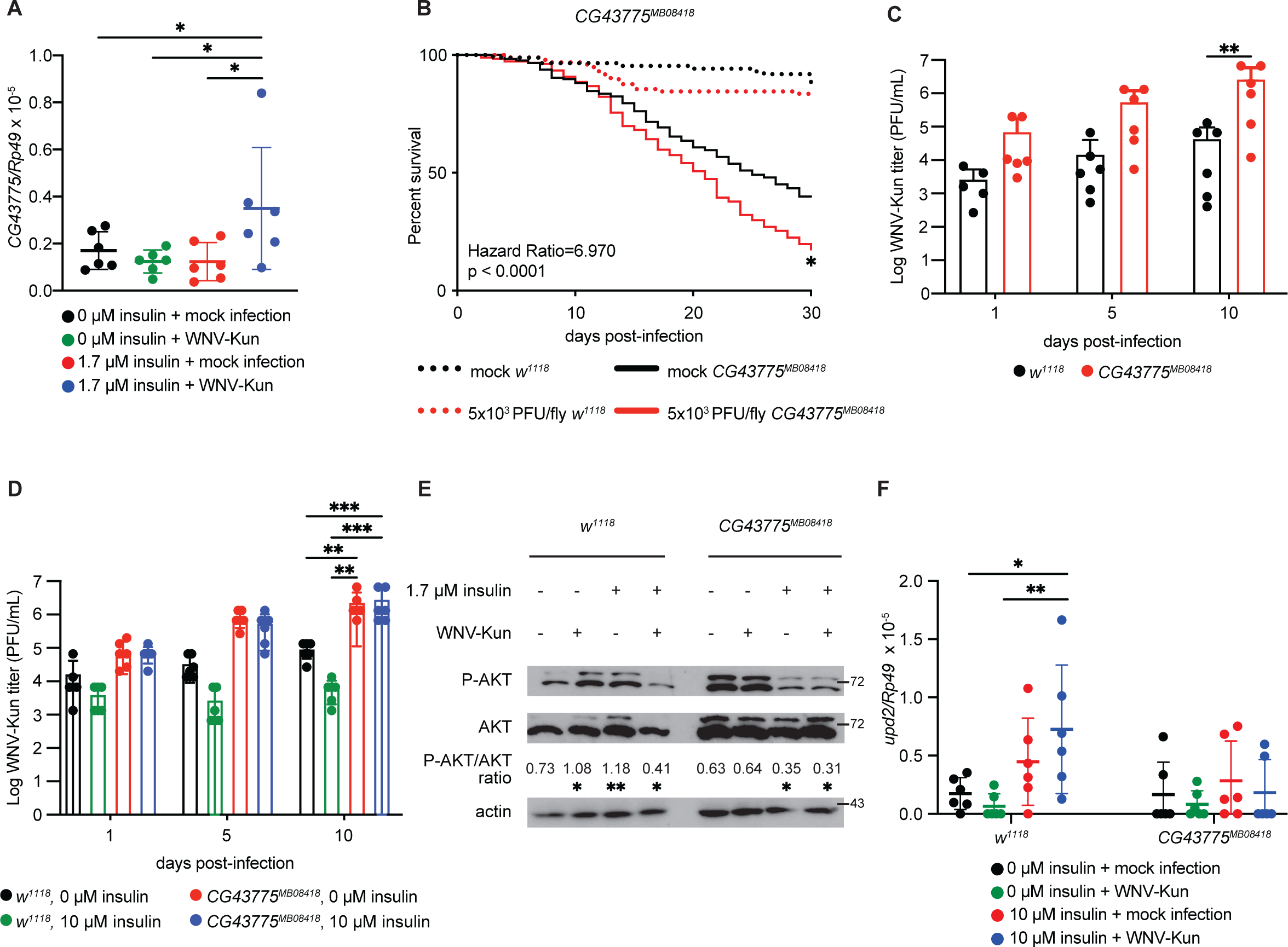
*CG43775* mutant flies are more susceptible to WNV-Kun infection due to deficient insulin-mediated antiviral protection. (A) *CG43775* is induced in *D. melanogaster* S2 cells that were insulin-treated and WNV-Kun infected (*p < 0.05, One-way ANOVA). (B) Flies with mutations in *CG43775* (solid red line) have higher mortality to WNV-Kun infection compared with the *w ^1118^* genetic control (dotted red line). (C) WNV-Kun titer is higher in *CG43775^MB08418^* flies relative to *w ^1118^* genetic control by 10 d p.i. (**p < 0.01, 2-way ANOVA). (D) Insulin treatment reduces WNV-Kun titer in control *w ^1118^* flies but not in *CG43775^MB08418^* flies (**p < 0.01, ***p < 0.001, 2-way ANOVA). (E) Akt is phosphorylated and active in the presence of insulin for *w ^1118^* flies but not in *CG43775^MB08418^* flies at 5 d p.i. (*p < 0.05, **p < 0.01, One-way ANOVA, see quantification in Fig. S2) (F) *CG43775^MB08418^* flies have impaired induction of *upd2* compared to genetic control *w ^1118^* flies. For qRT-PCR results, each circle represents individual biological replications consisting of individual well (A) or pooled collection of 3 flies (F). For titer results each circle represents individual biological replications consisting of pooled collection of 5 flies. Titer and qRT-PCR results (B-D, F) are representative of triplicate independent experiments western blot results are representative of duplicate independent experiments (E).

Upon establishing that *CG43775* impacts host survival and WNV-Kun replication, we sought to examine the role of *CG43775* in insulin-mediated antiviral immunity. We fed mutant and control flies 0 or 10 μM insulin two days prior to and during infection and collected flies at 1-, 5-, and 10 d p.i to measure virus replication (Fig. 2D). Similar to the previous results, we observed that mutant flies had higher viral titers relative to the genetic control. We also observed that while insulin-treated control flies had a reduction in viral titers, there was no difference between 0 or 10 μM insulin-treated *CG43775* mutant flies. These results indicate that loss of *CG43775* expression results in a loss of insulin-mediated reduction in viral replication.

To further dissect the role that *CG43775* has on insulin-mediated antiviral immunity, we sought to evaluate the impact that *CG43775* expression has on insulin signaling and JAK/STAT activation. Previous results demonstrate that insulin treatment of S2 cells activates AKT and JAK/STAT signaling, leading to the reduction of WNV-Kun (13). At 5 d p.i, we observed increased AKT phosphorylation in insulin-treated *w^1118^* flies compared to *CG43775* mutant flies (Fig. 2E) and quantified using densitometry analysis (Fig. S2). This leads us to conclude that *CG43775* mutant flies have a dysfunctional insulin signaling response that may impact insulin-mediated induction of antiviral JAK/STAT signaling. However, in the presence of insulin treatment and WNV-Kun infection, AKT phosphorylation was similar between genotypes, which may be due to virus-induced inhibition of AKT activation (49). Furthermore, *w^1118^* flies that were treated with insulin and infected with WNV-Kun had diminished AKT phosphorylation compared to flies that received either insulin or WNV-Kun. This may be caused by a secondary physiological signaling pathway which is absent *in vitro* and results in diminished AKT phosphorylation regardless of insulin treatment but remains sufficient to protect against WNV disease (50–52). Insulin treatment in S2 cells leads to the induction of *unpaired* (*upd*) cytokines and JAK/STAT activation (13). Thus, we examined *upd2* induction in control and *CG43775* mutant flies. At 5 d p.i, we observed significant induction of *upd2* in insulin-treated control flies, but not in *CG43775* mutant flies (Fig. 2F). Collectively these data suggest that *CG43775*, a previously uncharacterized gene that was identified within the endothelin signaling gene set cluster, contributes to antiviral immunity during WNV-Kun infection through canonical insulin and JAK/STAT signaling.

### Insulin and endothelin signaling reduce WNV-Kun replication in human HepG2 cells

The endothelin signaling pathway is not well-characterized in *D. melanogaster*; however, the pathway has been heavily dissected in mammals and permits us to investigate its potential role as an antiviral mediator to WNV-Kun in human cells. We first evaluated the extent to which insulin-mediated antiviral immunity functions in this model system. Human HepG2 liver cells were treated with either 0 or 1.7 μM bovine insulin and infected with WNV-Kun (MOI 0.01 PFU/cell). Viral titer was measured at 1, 2, 3, and 5 d p.i. (Fig. 3A). As previously observed in fruit fly and mosquito cells (13), we observed a significant reduction in viral titer in cells treated with insulin. We followed up viral titer analysis by comparing insulin-treated cells to cells that received interferon (IFN)-β or -γ treatment (Fig. S3) (53). IFN treatment is known to reduce WNV replication in human cells (54–58), so this comparison was to determine the efficacy of insulin in reducing WNV-Kun replication. We observed that insulin had a similar efficacy in reducing virus replication as IFN-γ and IFN-β at 2 d p.i.

**Figure 3:**
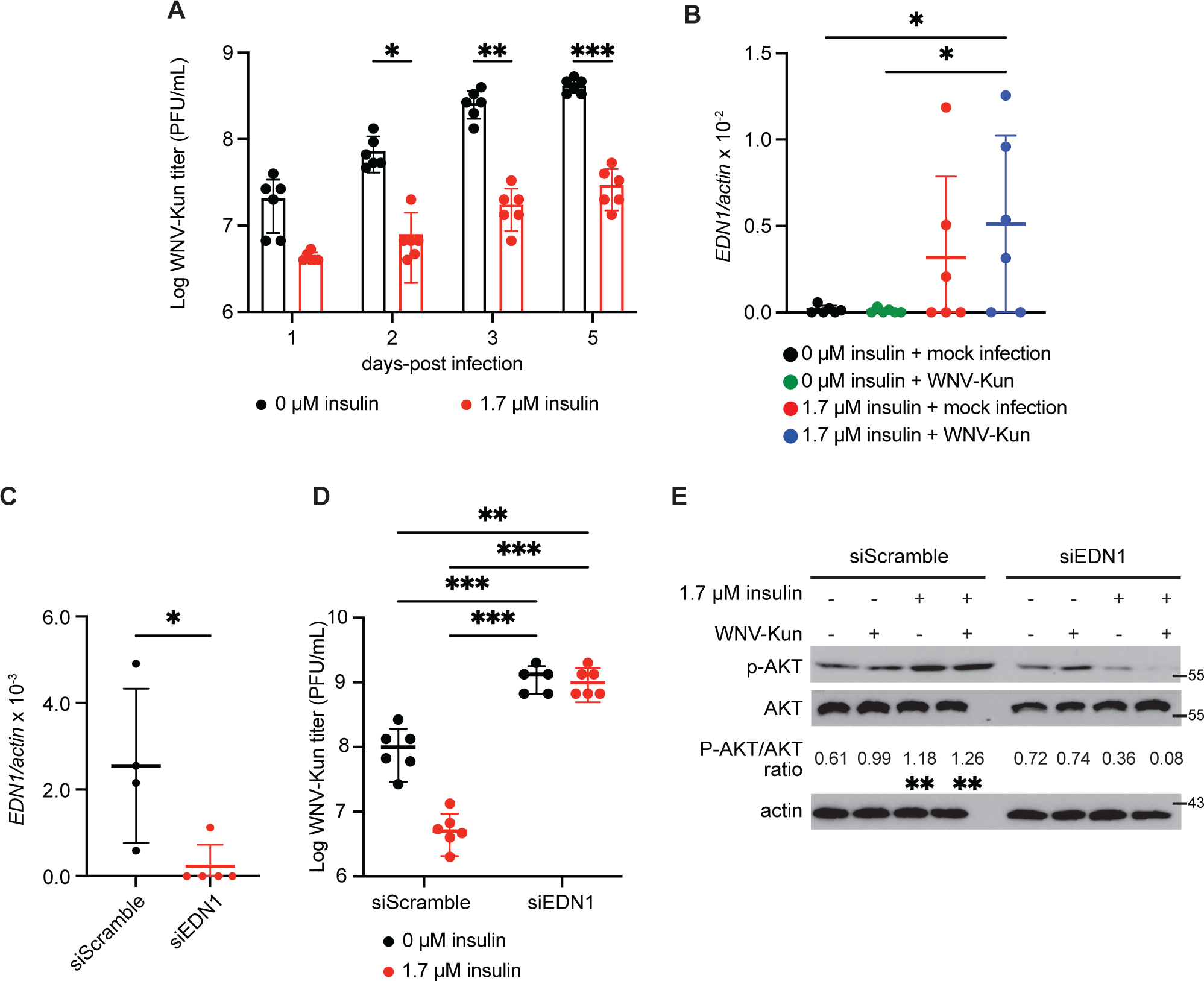
Endothelin signaling is antiviral to WNV-Kun through an insulin-dependent mechanism in human HepG2 cells. (A) Insulin-treatment of HepG2 cells reduces WNV-Kun titer (MOI 0.01 PFU/cell) (*p < 0.05, **p < 0.01, ***p < 0.001, 2-way ANOVA). (B) *EDN1* is induced in insulin-treated and WNV-Kun-infected HepG2 cells (*p < 0.05, One-way ANOVA). (C-E) *EDN1* was knocked down in HepG2 cells (C) (*p < 0.05, upaired t-test) 48h prior to insulin-treatment and WNV-Kun infection and viral titer was measured by standard plaque assay at 2 days-post infection (D) (**p < 0.01, ***p < 0.001, 2-way ANOVA). (E) Insulin-mediated Akt phosphorylation is decreased in the absence of *EDN1* (**p < 0.01, One-way ANOVA, see quantification in Fig. S4). Circles represent individual biological replications. Horizontal bars represent the mean. Error bars represent SDs. Titer and qRT-PCR results (A-D) are representative of triplicate independent experiments western blot results are representative of duplicate independent experiments (E).

To investigate endothelin signaling during insulin-mediated antiviral immunity, we measured induction of the ligand *endothelin 1* (*EDN1*) in HepG2 cells during WNV-Kun infection and insulin treatment (Fig. 3B). We observed significant induction of *EDN1* induction in the presence of insulin and WNV-Kun infection. This indicates that like our previous observations in *D. melanogaster*, endothelin signaling may be involved in insulin-mediated antiviral immunity in human cells. To further evaluate this hypothesis, we transfected HepG2 cells with either non-targeting control siRNA (siScramble) or EDN1 siRNA (siEDN1) (Fig. 3C). We observed a 91% reduction in *EDN1* expression in cells transfected with siEDN1. In cells knocked-down for EDN1 and treated with insulin, we observed that while the siScramble control cells maintain a reduction in WNV-Kun replication in the presence of insulin, we lose this insulin-mediated antiviral protection when EDN1 expression is diminished (Fig. 3D). We also observed a significant increase in overall WNV-Kun replication even in the absence of insulin treatment. Taken together, endothelin signaling may be connected with the insulin-mediated antiviral response previously observed by others in the mammalian system (59–62).

We next tested the role that *EDN1* expression has on insulin signaling by measuring phosphorylation of AKT in HepG2 cells following insulin treatment and WNV-Kun infection at 2 d p.i. These cells were also transfected with siScramble or siEDN1 (Fig. 3E). We observed that control cells had higher expression of P-AKT in the presence of insulin and infection while the loss of EDN1 had diminished P-AKT expression regardless of insulin-treatment (Fig. 3E) and quantified using densitometry analysis (Fig. S4). This further connects endothelin as a mediator of antiviral protection through an insulin-specific mechanism.

### Insulin- and endothelin-mediated signaling is antiviral to virulent WNV-NY99

Previous analysis of insulin-mediated antiviral immunity in an insect (13) and present mammalian context has used the attenuated Kunjin subtype of WNV. While useful in dissecting and evaluating host immunity to WNV in a general context, a present limitation is that this strain causes limited disease in immune-competent human hosts. This is due to a number of factors including increased sensitivity to type I interferon responses (63) and decreased efficacy in antagonizing JAK/STAT signaling due to a mutation in the NS5 protein (55). Because of this limitation regarding clinical relevance, we sought to evaluate whether insulin-mediated antiviral protection was present against more virulent flaviviruses and if so the impact that endothelin signaling possesses for regulating viral replication. Like previous experiments, we used HepG2 cells that received either 0 or 1.7 μM insulin treatment 24 h prior to and during WNV-NY99 (MOI 0.01 PFU/cell) infection and measured viral titer at 1, 2, 3, and 5 d p.i. (Fig. 4A). We observed that WNV-NY99 titer was reduced in cells that received insulin treatment. We also observed a higher virus titer in WNV-NY99 infected cells compared to WNV-Kun infected cells. Because of the established link that insulin signaling induces JAK/STAT in mammals (64, 65) and insects (13), this increase in overall viral titer is likely due to the enhanced antagonism WNV-NY99 can successfully initiate as opposed to the attenuated WNV-Kun strain (55, 63). We followed up this analysis by measuring WNV-NY99 titer in HepG2 cells that received either non-targeting or EDN1 siRNA (Fig. 4B). We observed a similar loss of insulin-mediated protection and increased viral load in siEDN1-transfected cells that was previously observed during WNV-Kun infection. This observation ultimately leads us to conclude that downstream components of insulin-mediated antiviral immunity, specifically endothelin signaling, plays a role in reducing WNV replication for both attenuated and more virulent strains that may be a potential target for future clinical or therapeutic research.

**Figure 4:**
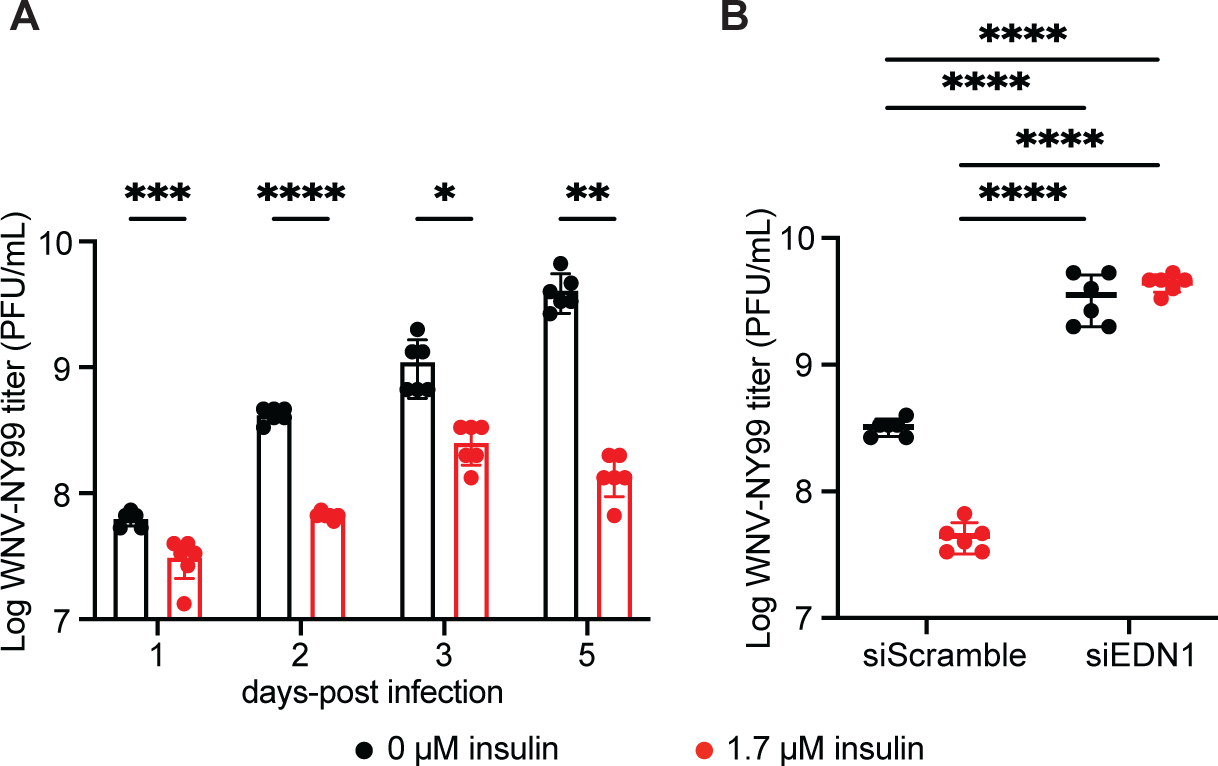
Endothelin and insulin-mediated signaling is conserved against more virulent WNV-NY99 strain in HepG2 cells. (A) Insulin-treatment reduces WNV-NY99 titer (MOI=0.01 PFU/cell) (*p < 0.05, **p < 0.01, ***p < 0.001, ****p < 0.0001, 2-way ANOVA). (B) siRNA silencing of EDN1 results in increased WNV-NY99 viral replication and loss of insulin-mediated protection compared to non-specific siScramble control at 2 days post-infection (****p < 0.0001, Two-way ANOVA). Circles represent individual biological replications. Horizontal bars represent the mean. Error bars represent SDs. Results are representative of triplicate independent experiments.

## DISCUSSION

Arbovirus infections are a growing health threat that require more effective means of intervention both environmentally (i.e., vector transmission) and clinically. While our ability to develop more effective vector control protocols has improved, the ability to understand and clinically address human infections and severe disease remains underdeveloped. As WNV, along with other mosquito-borne diseases, continue to expand in both global distribution and incidence (5, 7, 66, 67), the need for effective preventatives and treatments is more urgent than ever. Human vaccine development against WNV has made limited progress (68), so development of effective antivirals post exposure is necessary.

In the study presented here, we highlight the genetic power of *D. melanogaster* to advance the study of antiviral immunity and identify components of insect and mammalian host responses that regulate WNV infection. We demonstrate that insulin induces a number of genes and signaling pathways that are both canonical and previously unidentified antiviral mediators (Fig. 1). Our study using the *D. melanogaster* model expands upon the limited knowledge pertaining to the endothelin signaling pathway in insects, specifically in regards to host survival and viral replication in the insect (Fig. 2). We also demonstrate that we can use these results to translate our findings into the more pertinent human model (Fig. 3). In addition, we demonstrate that our findings are applicable to the more virulent and clinically relevant WNV strain NY99 (Fig. 4).

In our study, we show that dysfunctional endothelin signaling results in increased host mortality and WNV replication. However, further investigation is also necessary to evaluate its role during infection. Like insulin, endothelins are linked to various physiological processes like cardiovascular health so induction of this pathway, while potentially antiviral, may impact other off-target processes. Increased production and secretion of EDN1 has been used as an indicator for oncogenic and virus-induced hepatocellular carcinoma (24, 40, 69) as a promoter of cell growth and proliferation while inhibiting pro-apoptotic signaling (70). EDN1 expression is also proposed as a biomarker for patients receiving interferon-*α* treatment as elevated levels can be used to infer progression to interferon induced pulmonary toxicity (27). In relation to insulin sensitivity and signaling, serum EDN1 is elevated in diabetic individuals who later develop diabetic microangiopathy and nephropathy that progresses to more advanced insulin resistance (71, 72). Additional concerns are apparent as endothelin signaling, while antiviral in this study, may promote or enhance infection against other pathogens. *Mycobacterium tuberculosis* secretes the protease enzyme Zmp1 that cleaves EDN1 and activates endothelin signaling that promotes bacteria survival within the lungs (23). Thus, further investigation is needed to understand how targeting the endothelin pathway influences other related viruses that are either targeted by or disrupt insulin signaling in the presence or absence of WNV infection.

Determining the overall effect that insulin-mediated protection and endothelin signaling has in a clinical context will be important if targeting the pathways are used as an intervention for WNV disease. It is unlikely that administering insulin to a patient alone is a viable approach for treating WNV since it can influence a number of off-target physiological processes and may lead to further insulin resistance or disease pathology (18, 19). Instead, we propose through our study that by targeting pathways downstream of insulin signaling, we can effectively and directly induce more potent antiviral responses with limited toxicity to the host. While our study focused on endothelin signaling, there were other gene sets and associated pathways identified in our RNAseq screen that are worth further investigation regarding their potential role in antiviral immunity.

Taken together, our study identifies a novel component of insect and human antiviral immunity and expands our current understanding regarding insulin-mediated responses to infection. Previous investigation demonstrated that a variety of viruses including influenza (49), WNV (59, 73), and ZIKV (20, 74) target and disrupt host processes associated with insulin signaling. Typically, insulin signaling disruption results in metabolic dysfunction that can cause more severe morbidity and mortality. Here we demonstrate that targeting insulin signaling protects fruit flies and humans from increased viral replication. Additionally, we show that endothelin signaling provides antiviral immunity to WNV. While endothelins have been heavily dissected as a regulator of cardiovascular health and vasoconstriction (27, 28, 38), they also possess a role in hepatic (24, 34, 40, 69) and neuronal (75–78) regulation and health. WNV disease is heavily associated with encephalitis and neurodegenerative disease (79, 80). Because EDN1 has been linked to virus-induced demyelinating disease (75) and promotes anti-inflammatory signaling in circulating immune cells (26), endothelin signaling may also function as an antiviral target and determinant in severe WNV disease progression and is worth further investigation.

Given the conservation of insulin signaling and its activation during viral infection across insect and mammalian species, downstream targets of insulin or endothelin signaling may have a broader role within an antiviral context. If possible, it may provide a means of more effectively responding to these growing pathogens of concern while also limiting potential complications associated with current intensely robust antiviral therapeutics.

## MATERIALS AND METHODS

### Fly lines and rearing

Flies used in this study are listed in Table S3. Flies were maintained on standard cornmeal food (Genesee Scientific #66-112) at 25°C and 65% relativity humidity, and a 12 h/ 12 h light/dark cycle. Flies are negative for *Wolbachia* infection. Female adult flies used for all experiments were 2-7 d post-eclosion . For insulin treatment, cornmeal food was supplemented with 10 μM bovine insulin (Sigma-Aldrich I6634) and flies were maintained on food 48 h prior and during infection as described (13).

### Cells and virus

Vero cells (ATCC, CRL-81) were provided by A. Nicola and cultured at 37 °C/5% CO_2_ in DMEM (ThermoFisher 11965) supplemented with 10% FBS (Atlas Biologicals FS-0500-A) and 1x antibiotic-antimycotic (ThermoFisher 15240062). S2 cells were cultured as described (81) and are negative for Flock House virus. HepG2 cells (ATCC, HB-8065) were provided by M. Konkel and cultured at 37 °C/5% CO_2_ in DMEM supplemented with 10% FBS. For insulin treatment, culture media with 2% FBS were supplemented with 1.7 μM bovine insulin as described (82). For interferon-β and -γ treatment, 2% FBS in DMEM media was supplemented with 10 units/mL of either IFN-β or IFN-γ for 24 h prior to infection as described (53).

West Nile virus-Kunjin strain MRM16 (WNV-Kun) was gifted by R. Tesh and propagated in Vero cells. West Nile virus strain 385-99 (WNV-NY99) was obtained by BEI Resources, NIAID, NIH (NR-158) and propagated in Vero cells. All experiments with a specific virus type utilized the same stock.

### RNA isolation, library preparation, and RNA-sequencing

*D. melanogaster* S2 cells were treated with 0 or 1.7 μM bovine insulin for 24 h. Cells were then either mock-infected or infected with WNV-Kun (MOI 1 PFU/cell) for 8 h. Total RNA was extracted from three individual wells using Direct-zol (Zymo Research, Irvine, CA) following the manufacturer’s instructions. Following total RNA extraction, the integrity of total RNA was assessed using Fragment Analyzer (Advanced Analytical Technologies, Ankeny, IA) with the High Sensitivity RNA Analysis Kit. RNA Quality Numbers (RQNs) from 1 to 10 were assigned to each sample to indicate its integrity or quality. “10” stands for a perfect RNA sample without any degradation, whereas “1” marks a completely degraded sample. RNA samples with RQNs ranging from 8 to 10 were used for RNA library preparation with the TruSeq Stranded mRNA Library Prep Kit (Illumina, San Diego, CA). Briefly, mRNA was isolated from 2.5 µg of total RNA using poly-T oligo attached to magnetic beads and then subjected to fragmentation, followed by cDNA synthesis, dA-tailing, adaptor ligation, and PCR enrichment. The sizes of RNA libraries were assessed by Fragment Analyzer with the High Sensitivity NGS Fragment Analysis Kit. The concentrations of RNA libraries were measured by StepOnePlus Real-Time PCR System (ThermoFisher Scientific, San Jose, CA) with the KAPA Library Quantification Kit (Kapabiosystems, Wilmington, MA). The libraries were diluted to 2 nM in10 mM Tris-HCl, pH 8.5 and denatured with 0.1 N NaOH. Eighteen pM libraries were clustered in a high-output flow cell using HiSeq Cluster Kit v4 on a cBot (Illumina). After cluster generation, the flow cell was loaded onto HiSeq 2500 for sequencing using HiSeq SBS kit v4 (Illumina). DNA was sequenced from both ends (paired-end) with a read length of 100 bp. The raw bcl files were converted to fastq files using software program bcl2fastq2.17.1.14. Adaptors were trimmed from the fastq files during the conversion. On average, 40 million reads were generation for each sample. RNA-sequencing was performed at the Spokane Genomics CORE at Washington State University-Spokane in Spokane, WA, USA.

### Bioinformatics Analysis

RNA-seq reads were imported and aligned using Qiagen CLC Genomics Workbench 11.0.1 to the *D. melanogaster* genomic reference sequence. Reads for each biological replicate within an experimental condition were pooled and averaged. Differential expression of transcript levels for each experimental condition (WNV-Kun infection, insulin treatment, or both infection and treatment) were normalized to reads for cells that received neither treatment nor infection. Transcripts were filtered for p-values less than or equal to 0.05 and a log_2_(fold change) > ± 1.5 for at least one experimental condition.

Filtered transcripts were imported into Tibco Spotfire for gene clustering and heatmap generation. Gene clustering was performed using hierarchical clustering using UPGMA (unweighted pair group method with arithmetic mean) with Euclidean distance with ordering weight set to average value and normalization by mean. Gene set enrichment analysis (GSEA) was performed as previously described (83) using a cutoff of p < 0.05 for at least one experimental condition for gene ontology (GO) classifications (84). Highlighted classifications are shown in Figure 1B. *Drosophila* gene ontologies were imported from FlyBase (version fb_2016_04) as previously described (85). Further GO analysis for genes clustered and presented in Figure 1E used PANTHER GO-Slim (Version 14.0) to identify endothelin signaling pathway as an overrepresented GO category.

### Fly infections

2-7 day old adult female *D. melanogaster* were anesthetized with CO_2_ and injected intrathoracically with WNV-Kun with 5,000 PFU/fly, as previously described (12, 85). Mock infected-flies received equivalent volume of PBS. For mortality studies, groups of 30-50 flies were injected and maintained on cornmeal food for 30 days. All survival studies were repeated three times and survival data were combined. Fly food vials were changed every 2-3 days. For viral titration experiments, three groups of 4-5 flies were collected, homogenized in PBS, and used as individual samples for plaque assay as described in (13). For qRT-PCR and Western blot experiments, three groups of 3-5 flies were collected, homogenized in Trizol or RIPA, respectively, and centrifuged to isolate and remove cuticle. Supernatant was collected and used for further analysis.

### *In vitro* virus replication

HepG2 cells were seeded into a 24-well plate at a confluency of 1.25 x 10^5^ cells/well with 6 independent wells for each experimental condition. The following day, cells were treated with either 1.7 μM bovine insulin or acidified water in 2% FBS in DMEM for 24 h prior to infection. For measuring viral replication following interferon treatment, 2% FBS in DMEM media was supplemented with 10 units/mL of either IFN-β or IFN-γ for 24 h prior to infection as described (53). Cells were then infected with WNV-Kun or WNV-NY99 at MOI of 0.01 PFU/cell for 1 h. Virus inoculum was removed, and fresh experimental media was added. Supernatant samples were collected at 1, 2, 3, and 5 d p.i. for later titration. WNV titer were determined by standard plaque assay on Vero cells.

### Immunoblotting

Protein extracts were prepared by lysing cells or flies with RIPA buffer (25 mM Tris-HCl pH 7.6, 150 mM NaCl, 1 mM EDTA, 1% NP-40, 1% sodium deoxycholate, 0.1% SDS, 1mM Na_3_VO_4_, 1 mM NaF, 0.1 mM PMSF, 10 μM aprotinin, 5 μg/mL leupeptin, 1 μg/mL pepstatin A). Protein samples were diluted using 2x Laemmli loading buffer, mixed, and boiled for 5 minutes at 95 °C. Samples were analyzed by SDS/PAGE using a 10% acrylamide gel, followed by transfer onto PVDF membranes (Millipore IPVH00010). Membranes were blocked with 5% BSA (ThermoFisher BP9706) in Tris-buffered saline (50 mM Tris-HCl pH 7.5, 150 mM NaCl) and 0.1% Tween-20 for 1 h at room temperature.

Primary antibody labeling was completed with anti-P-Akt (1:1,000; Cell Signaling 4060), anti-Akt (pan) (1:2,000) (Cell Signaling 4691), or anti-actin (1:10,000; Sigma A2066) overnight at 4 °C. Secondary antibody labeling was completed using anti-rabbit IgG-HRP conjugate (1:10,000; Promega W401B) by incubating membranes for 2 h at room temperature. Blots were imaged onto film using luminol enhancer (ThermoFisher 1862124). P-AKT/AKT ratio for each experimental condition was determined using densitometry analysis using BioRad Image Lab comparing band intensity of P-AKT to AKT. Reported P-AKT/AKT ratio is the mean of duplicate independent experiments.

### RNA interference *in vitro*

Double-stranded RNA (dsRNA) targeting human *EDN1* (Horizon Discovery J-016692-05-005) and non-targeting control (siScramble) dsRNA (Horizon Discovery D-001810-10-05) was transfected into HepG2 cells for 48 h prior to insulin treatment and infection as described (81). Total RNA was extracted and purified to confirm reduced expression by qRT-PCR.

### Quantitative reverse transcriptase PCR

qRT-PCR was used to measure mRNA levels in *D. melanogaster* S2 cells, adult flies, and human HepG2 cells. Cells or flies were lysed with Trizol Reagent (ThermoFisher 15596). RNA was isolated by column purification (ZymoResearch R2050), DNase treated (ThermoFisher 18068), and cDNA was prepared (BioRad 170–8891). Expression of *D. melanogaster CG43775* and *upd2* were measured using SYBR Green reagents (ThermoFisher K0222) and normalized to *Rp49* to measure endogenous gene levels for all treatment conditions. Expression of human *EDN1* was measured using the probe for *EDN1* (Hs00174961_m1 ThermoFisher 4331182) and primers and normalized to *β-actin* (Hs01060665_g1 ThermoFisher 4331182) using TaqMan Universal Master Mix (ThermoFisher 4304437). The reaction for samples included one cycle of denaturation at 95 °C for 10 minutes, followed by 50 cycles of denaturation at 95 °C for 15 seconds and extension at 60 °C for 1 minute, using an Applied Biosystems 7500 Fast Real Time PCR System. ROX was used as an internal control. qRT-PCR primer sequences are listed in Table 3 Table (13, 86, 87).

### Quantification and Statistical Analysis

Results presented as dot plots show data from individual biological replicates (n=2-6), the arithmetic mean of the data shown as a horizontal line, and error bars representing standard deviations from the mean. Biological replicates of adult *D. melanogaster* (n=6-40) consisted of triplicate pooled flies. Results shown are representative of at least duplicate independent experiments, as indicated in the figure legends. All statistical analyses of biological replicates were completed using GraphPad Prism 9 and significance was defined as p < 0.05. Ordinary one-way ANOVA with uncorrected Fisher’s LSD for multiple comparisons was used for qRT-PCR analysis. Two-way ANOVA with Šidák correction for multiple comparisons was used for multiday viral titer analysis and for siRNA viral titer analysis. One-way ANOVA with Šidák correction for multiple comparisons was used for single day viral titer in the presence of insulin and interferon-β and -γ analysis. Two-tailed unpaired t test was used for qRT-PCR validation of knocked-down expression of *EDN1*. Repeated measures one-way ANOVA with uncorrected Fisher’s LSD for multiple comparison was used for densitometry analysis. All error bars represent standard deviation (SD) of the mean. Outliers were identified using a ROUT test (Q=5%) and removed.

## DATA AVAILABILTIY

Raw and processed RNAseq data have been deposited in NCBI Gene Expression Omnibus (GEO) Accession # GSE216532.

## ACKNOWLEDGEMENTS

We thank A. Nicola, M. Konkel, and R. Tesh for cells and viruses used in this study. We also thank the Spokane Genomics CORE at Washington State University for their preparation and guidance in the RNAseq analysis. We would like to thank S. Luckhart for constructive feedback regarding experimental direction and interpretation. This research was supported by the WSU College of Veterinary Medicine Stanley L. Adler research fund, NIH / National Institute of General Medical Sciences (NIGMS)-funded pre-doctoral fellowship (T32 GM008336) and a Poncin Fellowship to C.E.T. and L.R.H.A, NIH/NIGMS pre-doctoral fellowship T32 GM008336, WSU Research Assistantships for Diverse Scholars (RADS), and ARCS Foundation Fellowship to B.J.J., The funders had no role in study design, data collection and analysis, decision to publish, or preparation of the manuscript.

## AUTHOR CONTRIBUTIONS

Conceptualization, C.E.T., L.R.H.A, and A.G.G.; Methodology, C.E.T., L.R.H.A, and A.G.G.; Validation, C.E.T., E.H.R., B.J.J., A.B.C., S.F., and A.G.G..; Investigation, C.E.T., E.H.R., B.J.J., A.B.C., S.F., L.R.H.A, and A.G.G; Resources, A.G.G.; Writing – Original Draft, C.E.T.; Writing – Review and Editing, E.H.R., B.J.J., L.R.H.A., A.B.C., S.F., and A.G.G.; Visualization, C.E.T. and A.G.G.; Funding Acquisition, C.E.T., B.J.J., L.H.R.A, and A.G.G.

## SUPPLEMENTAL INFORMATION

**Table S1:** Summary of RNAseq reads (Sheet 1), PANTHER GO analysis results (Sheet 2), and expression values of selected gene cluster (Sheet 3) (related to Figure 1).

| Experiment Condition | Sample # | Average Total Reads | Average Total Mapped | Average % Mapped |
| --- | --- | --- | --- | --- |
| 0 μM insulin + mock infection | 1, 2, 3 | 78500232.67 | 73303857.33 | 93.43 |
| 1.7 μM insulin + mock infection | 4, 5, 6 | 80376152.67 | 74634122.67 | 92.89333333 |
| 0 μM insulin + WNV-Kun | 7, 8, 9 | 77176451.33 | 73079306.67 | 94.59333333 |
| 1.7 μM insulin + WNV-Kun | 10, 11, 12 | 84974130 | 77950122.67 | 91.94666667 |
|  | Average: | 80256741.67 | 74741852.33 | 93.21583333 |

| PANTHER Pathways | Endothelin signaling pathway | Unclassified |
| --- | --- | --- |
| (REF #) | 79 | 12625 |
| # genes | 1 | 11 |
| expected | 0.7 | 10.96 |
| Fold Enrichment | 14.58 | 1 |
| (+ / -) | + | + |
| P-value | 6.73E-02 | 1.00E+00 |

| Name | 0 μM insulin + WNV-Kun | 1.7 μM insulin + mock infection | 1.7 μM insulin + WNV-Kun |
| --- | --- | --- | --- |
| CG8180 | -0.31 | 4.32 | 4.24 |
| Cyp4p1 | -0.27 | 4.67 | 4.62 |
| betaNACtes6 | -0.1 | 5.4 | 4.09 |
| CG13043 | -0.08 | 5.63 | 3.35 |
| gudu | -0.02 | 5.4 | 3.66 |
| CG5326 | -0.05 | 3.95 | 5.3 |
| Acp53Ea | -0.03 | 3.69 | 5.37 |
| CG7299 | -0.01 | 3.96 | 5.26 |
| TotF | -0.00296 | 2.55 | 5.19 |
| CG42507 | -0.03 | 5.81 | 6.68 |
| CG13670 | -0.01 | 5.9 | 5.82 |
| CG43775 | -0.03 | 5.4 | 5.08 |
| ac | 0.04 | 4.73 | 5.29 |
| Ugt36Bc | 0.06 | 5.62 | 5.49 |
| CG31769 | 0.08 | 4.48 | 4.39 |
| CG42460 | 0.11 | 5.22 | 4.35 |
| MtnB | 0.11 | 5.95 | 5.47 |
| CG2962 | 0.15 | 5.51 | 5.76 |
| CG14354 | 0.1 | 3.28 | -2.9 |
| CG46275 | 0.17 | 7.48 | -0.31 |
| Amt | 0.27 | 4 | 3.71 |
| MtnD | 0.28 | 4.41 | 3.67 |
| CG32695 | 0.27 | 3.38 | 3.32 |

**Table S2:** GO classifications (p < 0.05) in 0 μM + WNV-Kun cells (Sheet 1), 1.7 μM insulin + mock infected cells (Sheet 2), and 1.7 μM insulin + WNV-Kun cells (Sheet 3) following GSEA (related to Figure 1B).

| Table S2_Mock-KUNV |  |  |  |  |  |  |
| --- | --- | --- | --- | --- | --- | --- |
| Rank | Function | Description | Depth | p-value | es score | # genes |
| 1 | GO:0005267 | potassium channel activi | 9 | 4.4297E-05 | 0.78501124 | 6 |
| 2 | GO:0005634 | nucleus | 7 | 6.5208E-05 | 0.14593226 | 1290 |
| 3 | GO:0005575 | cellular_component | 5 | 7.4794E-05 | 0.13761325 | 1122 |
| 4 | GO:0003674 | molecular_function | 5 | 7.4859E-05 | 0.13549395 | 1121 |
| 5 | GO:0008150 | transcription factor activi | 7 | 8.6355E-05 | 0.13646187 | 970 |
| 6 | GO:0016021 | integral to membrane | 5 | 8.8062E-05 | 0.13996146 | 951 |
| 7 | GO:0008270 | zinc ion binding | 7 | 9.7288E-05 | 0.1465636 | 860 |
| 8 | GO:0005737 | cytoplasm | 8 | 0.00012038 | 0.14684099 | 694 |
| 9 | GO:0005524 | ATP binding | 8 | 0.00012751 | 0.14633411 | 655 |
| 10 | GO:0003677 | DNA binding | 8 | 0.00015258 | 0.13992077 | 547 |
| 11 | GO:0006508 | proteolysis | 6 | 0.00015398 | 0.13971907 | 542 |
| 12 | GO:0022008 | neurogenesis | 6 | 0.00015716 | 0.18100269 | 436 |
| 13 | GO:0016020 | membrane | 5 | 0.00015746 | 0.14220663 | 530 |
| 14 | GO:0005515 | protein binding | 4 | 0.00017751 | 0.14528835 | 470 |
| 15 | GO:0003676 | nucleic acid binding | 5 | 0.00018455 | 0.15069042 | 329 |
| 16 | GO:0055114 | oxidation reduction | 6 | 0.00020356 | 0.1429417 | 410 |
| 17 | GO:0005576 | extracellular region | 7 | 0.00021067 | 0.13881292 | 397 |
| 18 | GO:0003700 | transcription factor activi | 6 | 0.00022312 | 0.14161317 | 375 |
| 19 | GO:0006355 | regulation of transcriptio | 6 | 0.00022623 | 0.141856 | 370 |
| 20 | GO:0005886 | plasma membrane | 7 | 0.00023378 | 0.14408903 | 358 |
| 21 | GO:0005622 | intracellular | 6 | 0.00023958 | 0.14348851 | 350 |
| 22 | GO:0034314 | Arp2/3 complex-mediate | 5 | 0.00024866 | 0.95015442 | 3 |
| 23 | GO:0000166 | nucleotide binding | 6 | 0.00028194 | 0.14486785 | 239 |
| 24 | GO:0016651 | oxidoreductase activity, a | 8 | 0.00028723 | 0.99985637 | 1 |
| 25 | GO:0005875 | microtubule associated c | 4 | 0.00029133 | 0.14366425 | 297 |
| 26 | GO:0055085 | transmembrane transpor | 5 | 0.00030778 | 0.14196671 | 289 |
| 27 | GO:0004252 | serine-type endopeptidas | 8 | 0.00034268 | 0.13879525 | 280 |
| 28 | GO:0005811 | lipid particle | 6 | 0.00034872 | 0.22770592 | 189 |
| 29 | GO:0005739 | mitochondrion | 9 | 0.0003782 | 0.18068545 | 169 |
| 30 | GO:0005488 | binding | 6 | 0.00042379 | 0.1425647 | 248 |
| 31 | GO:0006412 | translation | 7 | 0.00044801 | 0.29830023 | 88 |
| 32 | GO:0000398 | nuclear mRNA splicing, v | 7 | 0.00045786 | 0.23125696 | 115 |
| 33 | GO:0007052 | mitotic spindle organizat | 5 | 0.00046038 | 0.24738278 | 160 |
| 34 | GO:0003735 | structural constituent of | 6 | 0.00048165 | 0.31060548 | 84 |
| 35 | GO:0003729 | mRNA binding | 7 | 0.00049357 | 0.21112823 | 123 |
| 36 | GO:0005615 | extracellular space | 8 | 0.00053064 | -0.1919234 | 105 |
| 37 | GO:0006468 | protein amino acid phosph | 4 | 0.00056053 | 0.14285714 | 227 |
| 38 | GO:0071011 | #N/A | 7 | 0.00058675 | 0.26099969 | 94 |
| 39 | GO:0043565 | sequence-specific DNA b | 4 | 0.00058967 | 0.14105279 | 229 |
| 40 | GO:0008152 | metabolic process | 5 | 0.00060547 | 0.14231499 | 224 |
| 41 | GO:0005214 | structural constituent of | 9 | 0.0006265 | -0.2516674 | 52 |
| 42 | GO:0006334 | nucleosome assembly | 6 | 0.00064584 | -0.3948263 | 52 |
| 43 | GO:0006333 | chromatin assembly or d | 6 | 0.00069425 | -0.4061024 | 47 |
| 44 | GO:0000786 | nucleosome | 8 | 0.00071205 | -0.4438224 | 49 |
| 45 | GO:0060261 | positive regulation of tra | 7 | 0.00071609 | 0.98111175 | 2 |
| 46 | GO:0001096 | #N/A | 7 | 0.00071609 | 0.98111175 | 2 |
| 47 | GO:0071013 | #N/A | 5 | 0.0007221 | 0.23626557 | 74 |
| 48 | GO:0005549 | odorant binding | 8 | 0.00073724 | -0.375216 | 55 |
| 49 | GO:0007501 | mesodermal cell fate spe | 5 | 0.0007425 | -0.5661516 | 1 |
| 50 | GO:0006457 | protein folding | 5 | 0.00081599 | 0.20864008 | 87 |
| 51 | GO:0007606 | sensory perception of che | 6 | 0.00091539 | -0.3114819 | 50 |
| 52 | GO:0042744 | hydrogen peroxide catabo | 6 | 0.00098172 | 0.72564655 | 6 |
| 53 | GO:0007608 | sensory perception of sm | 6 | 0.00103421 | -0.262884 | 49 |
| 54 | GO:0046331 | lateral inhibition | 6 | 0.00104244 | 0.1423472 | 199 |
| 55 | GO:0005840 | ribosome | 4 | 0.00106783 | 0.39194774 | 44 |
| 56 | GO:0005829 | cytosol | 6 | 0.00110693 | 0.21984129 | 85 |
| 57 | GO:0032504 | multicellular organism re | 5 | 0.00111052 | -0.359922 | 29 |
| 58 | GO:0000022 | mitotic spindle elongatio | 8 | 0.00111315 | 0.29644069 | 50 |
| 59 | GO:0005509 | calcium ion binding | 7 | 0.00112203 | 0.14203511 | 197 |
| 60 | GO:0005751 | mitochondrial respiratory | 9 | 0.00123637 | 0.43655929 | 10 |
| 61 | GO:0048477 | oogenesis | 5 | 0.00128305 | 0.14283634 | 190 |
| 62 | GO:0051082 | unfolded protein binding | 4 | 0.00137207 | 0.22702271 | 65 |
| 63 | GO:0050909 | sensory perception of tas | 5 | 0.00138794 | -0.4086687 | 24 |
| 64 | GO:0004984 | olfactory receptor activity | 7 | 0.00138795 | -0.3939276 | 28 |
| 65 | GO:0005681 | spliceosome | 5 | 0.00141251 | 0.34358488 | 44 |
| 66 | GO:0008527 | taste receptor activity | 8 | 0.00154205 | -0.4010557 | 22 |
| 67 | GO:0005975 | carbohydrate metabolic p | 5 | 0.00159235 | 0.28114836 | 48 |
| 68 | GO:0022625 | cytosolic large ribosomal | 6 | 0.00166528 | 0.49590372 | 33 |
| 69 | GO:0005762 | mitochondrial large ribos | 7 | 0.00177154 | 0.41713564 | 35 |
| 70 | GO:0006911 | phagocytosis, engulfmen | 8 | 0.00179324 | 0.14229594 | 180 |
| 71 | GO:0009593 | detection of chemical sti | 8 | 0.00194679 | -0.3258219 | 22 |
| 72 | GO:0004175 | endopeptidase activity | 8 | 0.00194951 | 0.25463929 | 46 |
| 73 | GO:0031145 | anaphase-promoting com | 9 | 0.00208312 | 0.8204281 | 4 |
| 74 | GO:0003924 | GTPase activity | 8 | 0.00251232 | 0.15976925 | 86 |
| 75 | GO:0019861 | flagellum | 7 | 0.00255884 | -0.684842 | 1 |
| 76 | GO:0043169 | cation binding | 9 | 0.00267255 | 0.27769671 | 31 |
| 77 | GO:0042981 | regulation of apoptosis | 4 | 0.00276589 | 0.42399124 | 13 |
| 78 | GO:0008010 | structural constituent of | 4 | 0.00290441 | -0.2923066 | 15 |
| 79 | GO:0015276 | ligand-gated ion channel | 4 | 0.00291154 | -0.2457158 | 33 |
| 80 | GO:0005730 | nucleolus | 4 | 0.00294518 | 0.22857143 | 46 |
| 81 | GO:0007391 | dorsal closure | 7 | 0.00313166 | 0.20098976 | 77 |
| 82 | GO:0042776 | mitochondrial ATP synthe | 5 | 0.00330317 | 0.99834829 | 1 |
| 83 | GO:0045263 | proton-transporting ATP s | 7 | 0.00330317 | 0.99834829 | 1 |
| 84 | GO:0004674 | protein serine/threonine | 6 | 0.00337868 | 0.14217218 | 161 |
| 85 | GO:0005525 | GTP binding | 7 | 0.00347973 | 0.14223449 | 160 |
| 86 | GO:0000278 | mitotic cell cycle | 7 | 0.00378567 | 0.34674727 | 27 |
| 87 | GO:0035002 | liquid clearance, open tra | 8 | 0.00385078 | -0.6219556 | 7 |
| 88 | GO:0030833 | regulation of actin filament | 8 | 0.00388987 | 0.50643191 | 9 |
| 89 | GO:0006123 | mitochondrial electron tr | 6 | 0.00392953 | 0.45232276 | 8 |
| 90 | GO:0031175 | neurite development | 6 | 0.00408652 | 0.87315952 | 3 |
| 91 | GO:0009609 | response to symbiotic ba | 7 | 0.00415214 | 0.95446711 | 2 |
| 92 | GO:0016023 | cytoplasmic membrane-b | 7 | 0.00433752 | 0.61649544 | 7 |
| 93 | GO:0009055 | electron carrier activity | 5 | 0.00440331 | 0.14145668 | 155 |
| 94 | GO:0007165 | signal transduction | 6 | 0.00456679 | 0.14236968 | 152 |
| 95 | GO:0032509 | endosome transport via r | 6 | 0.00459572 | 0.99770197 | 1 |
| 96 | GO:0051298 | centrosome duplication | 5 | 0.00463809 | 0.22423124 | 42 |
| 97 | GO:0007411 | axon guidance | 6 | 0.00465272 | -0.1513878 | 26 |
| 98 | GO:0048039 | ubiquinone binding | 4 | 0.00488295 | 0.99755835 | 1 |
| 99 | GO:0042254 | ribosome biogenesis | 7 | 0.00502559 | 0.40972929 | 16 |
| 100 | GO:0030532 | small nuclear ribonucleop | 3 | 0.00510983 | 0.2687225 | 31 |
| 101 | GO:0007594 | puparial adhesion | 6 | 0.00511152 | -0.4956522 | 4 |
| 102 | GO:0005763 | mitochondrial small ribos | 6 | 0.00541877 | 0.33053028 | 26 |
| 103 | GO:0005732 | small nucleolar ribonucle | 9 | 0.00547112 | 0.77136906 | 4 |
| 104 | GO:0006397 | mRNA processing | 6 | 0.0055465 | 0.38075486 | 16 |
| 105 | GO:0010797 | regulation of multivesicu | 8 | 0.0055601 | 0.85944121 | 3 |
| 106 | GO:0004140 | dephospho-CoA kinase ac | 9 | 0.00568593 | 0.94671072 | 2 |
| 107 | GO:0048803 | imaginal disc-derived ma | 6 | 0.00596425 | 0.45278074 | 11 |
| 108 | GO:0045944 | positive regulation of tra | 8 | 0.00645079 | 0.14163402 | 144 |
| 109 | GO:0005887 | integral to plasma memb | 7 | 0.00695345 | 0.14111587 | 143 |
| 110 | GO:0035218 | leg disc development | 9 | 0.00738506 | 0.75355552 | 4 |
| 111 | GO:0022627 | cytosolic small ribosoma | 7 | 0.00749155 | 0.27376531 | 17 |
| 112 | GO:0005783 | endoplasmic reticulum | 8 | 0.00749507 | 0.14159292 | 140 |
| 113 | GO:0003756 | protein disulfide isomera | 8 | 0.00752068 | 0.50160966 | 9 |
| 114 | GO:0045434 | negative regulation of fe | 8 | 0.00763791 | -0.5258557 | 2 |
| 115 | GO:0017051 | retinol dehydratase activ | 8 | 0.00774365 | 0.93780523 | 2 |
| 116 | GO:0007527 | adult somatic muscle dev | 8 | 0.00787378 | 0.47797341 | 11 |
| 117 | GO:0051775 | response to redox state | 8 | 0.00789889 | 0.99605027 | 1 |
| 118 | GO:0020037 | heme binding | 8 | 0.00843894 | 0.14148959 | 137 |
| 119 | GO:0032027 | myosin light chain bindin | 9 | 0.00873816 | -0.4735897 | 4 |
| 120 | GO:0008379 | thioredoxin peroxidase ac | 8 | 0.0087403 | 0.5827697 | 6 |
| 121 | GO:0007067 | mitosis | 9 | 0.0088628 | 0.16984667 | 76 |
| 122 | GO:0000922 | spindle pole | 4 | 0.00886973 | -0.3654388 | 3 |
| 123 | GO:0006364 | rRNA processing | 8 | 0.00920205 | 0.3321833 | 17 |
| 124 | GO:0050916 | sensory perception of sw | 6 | 0.00943515 | -0.4916786 | 3 |
| 125 | GO:0046961 | proton-transporting ATPa | 7 | 0.00971196 | 0.3528481 | 16 |
| 126 | GO:0006506 | GPI anchor biosynthetic p | 7 | 0.00973186 | -0.3717764 | 2 |
| 127 | GO:0031072 | heat shock protein bindin | 9 | 0.01007816 | 0.24542573 | 42 |
| 128 | GO:0046933 | hydrogen ion transporting | 7 | 0.01017061 | 0.36083684 | 16 |
| 129 | GO:0004558 | alpha-glucosidase activit | 7 | 0.01021392 | 0.43180976 | 12 |
| 130 | GO:0015992 | proton transport | 7 | 0.01044096 | 0.30803507 | 17 |
| 131 | GO:0007095 | mitotic cell cycle G2/M t | 6 | 0.01062272 | 0.20364245 | 44 |
| 132 | GO:0017128 | phospholipid scramblase | 6 | 0.01065671 | 0.92703246 | 2 |
| 133 | GO:0017121 | phospholipid scrambling | 8 | 0.01065671 | 0.92703246 | 2 |
| 134 | GO:0048813 | dendrite morphogenesis | 8 | 0.01093968 | 0.14205987 | 129 |
| 135 | GO:0055092 | sterol homeostasis | 5 | 0.01105845 | 0.99447038 | 1 |
| 136 | GO:0030032 | lamellipodium assembly | 9 | 0.01109834 | 0.61099138 | 6 |
| 137 | GO:0048102 | autophagic cell death | 7 | 0.0111669 | 0.21298162 | 47 |
| 138 | GO:0035099 | hemocyte migration | 4 | 0.01127912 | 0.39984185 | 13 |
| 139 | GO:0003723 | RNA binding | 9 | 0.01171851 | 0.14502093 | 94 |
| 140 | GO:0045454 | cell redox homeostasis | 9 | 0.01173489 | 0.22624099 | 41 |
| 141 | GO:0042595 | behavioral response to st | 9 | 0.01182528 | 0.81922 | 3 |
| 142 | GO:0004129 | cytochrome-c oxidase act | 7 | 0.01198553 | 0.32101575 | 11 |
| 143 | GO:0005685 | snRNP U1 | 9 | 0.01212354 | 0.48038229 | 10 |
| 144 | GO:0051258 | protein polymerization | 6 | 0.01229074 | 0.44056346 | 10 |
| 145 | GO:0006296 | nucleotide-excision repai | 7 | 0.012351 | 0.99382406 | 1 |
| 146 | GO:0000262 | mitochondrial chromosom | 5 | 0.01249461 | 0.99375224 | 1 |
| 147 | GO:0016491 | oxidoreductase activity | 2 | 0.01268504 | 0.17310153 | 79 |
| 148 | GO:0030170 | pyridoxal phosphate bind | 4 | 0.01272028 | 0.25086418 | 30 |
| 149 | GO:0005913 | cell-cell adherens junctio | 6 | 0.01289547 | -0.5014012 | 9 |
| 150 | GO:0045175 | basal protein localization | 6 | 0.01324957 | -0.6011494 | 6 |
| 151 | GO:0005673 | transcription factor TFIIE | 5 | 0.01341525 | 0.91812698 | 2 |
| 152 | GO:0004672 | protein kinase activity | 5 | 0.01353991 | 0.14228791 | 123 |
| 153 | GO:0006409 | tRNA export from nucleu | 7 | 0.01364354 | 0.99317774 | 1 |
| 154 | GO:0006184 | GTP catabolic process | 4 | 0.01365307 | 0.27416685 | 22 |
| 155 | GO:0046620 | regulation of organ grow | 7 | 0.01369695 | -0.3923658 | 2 |
| 156 | GO:0003839 | gamma-glutamylcyclotra | 8 | 0.01369904 | 0.91726515 | 2 |
| 157 | GO:0004386 | helicase activity | 7 | 0.01397871 | -0.2288717 | 5 |
| 158 | GO:0005671 | Ada2/Gcn5/Ada3 transcr | 6 | 0.01427207 | 0.45224578 | 9 |
| 159 | GO:0016209 | antioxidant activity | 4 | 0.01430496 | 0.41803184 | 9 |
| 160 | GO:0009041 | uridylate kinase activity | 9 | 0.01479247 | 0.99260323 | 1 |
| 161 | GO:0004127 | cytidylate kinase activity | 6 | 0.01479247 | 0.99260323 | 1 |
| 162 | GO:0042060 | wound healing | 6 | 0.01492927 | 0.43222653 | 12 |
| 163 | GO:0035009 | negative regulation of m | 9 | 0.01526274 | 0.91266877 | 2 |
| 164 | GO:0007186 | G-protein coupled recept | 8 | 0.01572785 | 0.14118073 | 121 |
| 165 | GO:0031937 | positive regulation of chr | 4 | 0.01574323 | 0.91130422 | 2 |
| 166 | GO:0005747 | mitochondrial respiratory | 4 | 0.01619612 | 0.23260337 | 14 |
| 167 | GO:0008340 | determination of adult lif | 9 | 0.01678438 | 0.1419467 | 118 |
| 168 | GO:0060213 | positive regulation of nuc | 8 | 0.01686776 | -0.7001868 | 4 |
| 169 | GO:0010171 | body morphogenesis | 6 | 0.01703454 | -0.3355936 | 11 |
| 170 | GO:0006357 | regulation of transcrip | 4 | 0.01710191 | 0.14165701 | 118 |
| 171 | GO:0005694 | chromosome | 6 | 0.01755924 | 0.2943697 | 25 |
| 172 | GO:0007155 | cell adhesion | 5 | 0.01798202 | 0.14207096 | 116 |
| 173 | GO:0006777 | Mo-molybdopterin cofact | 6 | 0.01808167 | 0.54422013 | 7 |
| 174 | GO:0015734 | taurine transport | 6 | 0.01809307 | 0.90491238 | 2 |
| 175 | GO:0003713 | transcription coactivator | 6 | 0.01813083 | 0.25741025 | 29 |
| 176 | GO:0030127 | COPII vesicle coat | 7 | 0.01821493 | 0.79120879 | 3 |
| 177 | GO:0007375 | anterior midgut invagina | 9 | 0.01824508 | -0.6950151 | 4 |
| 178 | GO:0035071 | salivary gland cell autoph | 4 | 0.01835671 | 0.18882532 | 50 |
| 179 | GO:0006520 | amino acid metabolic pro | 4 | 0.01842862 | 0.34005215 | 17 |
| 180 | GO:0005952 | cAMP-dependent protein | 5 | 0.01894281 | -0.5806753 | 6 |
| 181 | GO:0007398 | ectoderm development | 5 | 0.01913498 | -0.3306005 | 21 |
| 182 | GO:0016577 | histone demethylation | 7 | 0.01917961 | -0.691711 | 4 |
| 183 | GO:0007394 | dorsal closure, elongatio | 7 | 0.01931567 | 0.43844772 | 9 |
| 184 | GO:0007499 | ectoderm and mesoderm | 9 | 0.01962141 | -0.7859657 | 3 |
| 185 | GO:0030010 | establishment of cell pol | 8 | 0.01964213 | -0.6901307 | 4 |
| 186 | GO:0004601 | peroxidase activity | 9 | 0.01975163 | 0.35595235 | 11 |
| 187 | GO:0006308 | DNA catabolic process | 7 | 0.01991249 | 0.90024418 | 2 |
| 188 | GO:0090072 | #N/A | 6 | 0.02008982 | -0.6269665 | 5 |
| 189 | GO:0005858 | axonemal dynein comple | 7 | 0.02010609 | -0.4365792 | 3 |
| 190 | GO:0007154 | cell communication | 6 | 0.02020956 | 0.4188587 | 12 |
| 191 | GO:0006813 | potassium ion transport | 6 | 0.02043062 | 0.26536185 | 21 |
| 192 | GO:0006950 | response to stress | 9 | 0.02068778 | -0.3060171 | 7 |
| 193 | GO:0004334 | fumarylacetoacetase act | 5 | 0.02082436 | 0.98958707 | 1 |
| 194 | GO:0005742 | mitochondrial outer mem | 5 | 0.02094668 | 0.53592725 | 6 |
| 195 | GO:0003785 | actin monomer binding | 6 | 0.02111159 | 0.98944345 | 1 |
| 196 | GO:0008143 | poly(A) binding | 5 | 0.02138828 | 0.62294375 | 5 |
| 197 | GO:0045143 | homologous chromosom | 7 | 0.02236661 | 0.77641313 | 3 |
| 198 | GO:0003779 | actin binding | 6 | 0.02296813 | 0.1418645 | 110 |
| 199 | GO:0035064 | methylated histone resid | 8 | 0.02338019 | 0.56817529 | 5 |
| 200 | GO:0006408 | snRNA export from nucle | 7 | 0.02346924 | 0.89169779 | 2 |
| 201 | GO:0000062 | acyl-CoA binding | 9 | 0.02350871 | 0.47143781 | 6 |
| 202 | GO:0000122 | negative regulation of tra | 4 | 0.02368771 | 0.14135785 | 110 |
| 203 | GO:0006366 | transcription from RNA p | 7 | 0.02385198 | 0.21069954 | 19 |
| 204 | GO:0080134 | #N/A | 4 | 0.0239383 | 0.89062051 | 2 |
| 205 | GO:0008378 | galactosyltransferase act | 4 | 0.02394087 | 0.49696436 | 5 |
| 206 | GO:0010821 | regulation of mitochondr | 9 | 0.02410523 | 0.61511386 | 5 |
| 207 | GO:0005686 | snRNP U2 | 9 | 0.02467652 | 0.39456181 | 11 |
| 208 | GO:0000980 | #N/A | 8 | 0.02498117 | -0.3374557 | 4 |
| 209 | GO:0007476 | imaginal disc-derived wi | 10 | 0.02542357 | 0.14212316 | 107 |
| 210 | GO:0009107 | lipoate biosynthetic proce | 5 | 0.02543808 | 0.88724504 | 2 |
| 211 | GO:0008137 | NADH dehydrogenase (uk | 7 | 0.02575878 | 0.24322054 | 22 |
| 212 | GO:0005543 | phospholipid binding | 5 | 0.02632991 | -0.1697833 | 22 |
| 213 | GO:0009649 | entrainment of circadian | 8 | 0.02670791 | -0.4230686 | 11 |
| 214 | GO:0008033 | tRNA processing | 9 | 0.02674693 | 0.4060299 | 11 |
| 215 | GO:0070868 | #N/A | 7 | 0.02698097 | -0.5213018 | 7 |
| 216 | GO:0004766 | spermidine synthase acti | 8 | 0.02721747 | 0.88336685 | 2 |
| 217 | GO:0046329 | negative regulation of JN | 5 | 0.02728819 | 0.44134809 | 8 |
| 218 | GO:0007173 | epidermal growth factor | 8 | 0.02779408 | 0.25947602 | 24 |
| 219 | GO:0008012 | structural constituent of | 4 | 0.02786968 | -0.6664991 | 4 |
| 220 | GO:0042049 | cellular acyl-CoA homeos | 6 | 0.02804642 | 0.51901307 | 5 |
| 221 | GO:0000796 | condensin complex | 6 | 0.02839938 | -0.4874982 | 8 |
| 222 | GO:0004053 | arginase activity | 6 | 0.02860867 | 0.88042229 | 2 |
| 223 | GO:0042811 | pheromone biosynthetic | 9 | 0.02862727 | 0.75723623 | 3 |
| 224 | GO:0001191 | #N/A | 6 | 0.02872325 | 0.98563734 | 1 |
| 225 | GO:0031887 | lipid particle transport al | 8 | 0.02893317 | -0.7563743 | 3 |
| 226 | GO:0008288 | boss receptor activity | 8 | 0.02901048 | 0.98549372 | 1 |
| 227 | GO:0007009 | plasma membrane organ | 9 | 0.02941163 | -0.4596537 | 9 |
| 228 | GO:0007413 | axonal fasciculation | 10 | 0.0296921 | 0.36083675 | 14 |
| 229 | GO:0021782 | glial cell development | 6 | 0.02971849 | -0.5534483 | 6 |
| 230 | GO:0007424 | open tracheal system dev | 7 | 0.03004573 | 0.14193735 | 103 |
| 231 | GO:0000236 | mitotic prometaphase | 3 | 0.03030303 | 0.9848474 | 1 |
| 232 | GO:0035556 | #N/A | 7 | 0.03046704 | -0.1595121 | 33 |
| 233 | GO:0005086 | ARF guanyl-nucleotide ex | 4 | 0.0307539 | -0.5985202 | 5 |
| 234 | GO:0032012 | regulation of ARF protein | 6 | 0.0307539 | -0.5985202 | 5 |
| 235 | GO:0006096 | glycolysis | 7 | 0.03082621 | -0.2848687 | 1 |
| 236 | GO:0050767 | regulation of neurogenes | 7 | 0.03107088 | 0.59780188 | 5 |
| 237 | GO:0051726 | regulation of cell cycle | 5 | 0.03137036 | 0.22519804 | 34 |
| 238 | GO:0005099 | Ras GTPase activator act | 7 | 0.03139045 | -0.5970835 | 5 |
| 239 | GO:0006962 | male-specific antibacteri | 5 | 0.03159558 | 0.98420108 | 1 |
| 240 | GO:0015914 | phospholipid transport | 5 | 0.03173385 | -0.5116747 | 7 |
| 241 | GO:0015662 | ATPase activity, coupled t | 5 | 0.03173385 | -0.5116747 | 7 |
| 242 | GO:0016592 | Srb-mediator complex | 5 | 0.03191814 | 0.24710286 | 27 |
| 243 | GO:0004985 | opioid receptor activity | 8 | 0.03274451 | 0.98362657 | 1 |
| 244 | GO:0005832 | chaperonin-containing T- | 8 | 0.03281414 | 0.50966305 | 7 |
| 245 | GO:0016272 | prefoldin complex | 9 | 0.03309251 | 0.43146019 | 7 |
| 246 | GO:0045177 | apical part of cell | 8 | 0.03333181 | -0.3178369 | 3 |
| 247 | GO:0045451 | pole plasm oskar mRNA | 7 | 0.03352806 | 0.2574122 | 30 |
| 248 | GO:0006614 | SRP-dependent cotransla | 6 | 0.0336093 | 0.43065536 | 10 |
| 249 | GO:0007366 | periodic partitioning by p | 7 | 0.03408766 | -0.507364 | 7 |
| 250 | GO:0003774 | motor activity | 7 | 0.03449765 | -0.2146493 | 7 |
| 251 | GO:0007275 | multicellular organismal | 9 | 0.03468843 | 0.25626079 | 20 |
| 252 | GO:0030728 | ovulation | 5 | 0.03472591 | 0.65105588 | 4 |
| 253 | GO:0006952 | defense response | 5 | 0.03481398 | 0.14140026 | 100 |
| 254 | GO:0016319 | mushroom body develop | 8 | 0.03549324 | -0.1893879 | 15 |
| 255 | GO:0004423 | iduronate-2-sulfatase act | 7 | 0.0361913 | 0.98190305 | 1 |
| 256 | GO:0045010 | actin nucleation | 7 | 0.03643284 | 0.86505315 | 2 |
| 257 | GO:0032296 | double-stranded RNA-spe | 7 | 0.03676576 | 0.9816158 | 1 |
| 258 | GO:0004844 | uracil DNA N-glycosylase | 6 | 0.03676576 | 0.9816158 | 1 |
| 259 | GO:0016560 | protein import into perox | 8 | 0.03690938 | 0.98154399 | 1 |
| 260 | GO:0002121 | inter-male aggressive be | 7 | 0.03692272 | 0.20803496 | 38 |
| 261 | GO:0000276 | mitochondrial proton-tra | 5 | 0.0369595 | 0.37609541 | 7 |
| 262 | GO:0042800 | histone methyltransferas | 6 | 0.03712683 | -0.6462434 | 4 |
| 263 | GO:0006605 | protein targeting | 7 | 0.03738999 | 0.5387931 | 5 |
| 264 | GO:0005902 | microvillus | 9 | 0.03755851 | 0.64541733 | 3 |
| 265 | GO:0001738 | morphogenesis of a pola | 6 | 0.03771926 | -0.3398185 | 3 |
| 266 | GO:0006833 | water transport | 10 | 0.03775689 | -0.6450223 | 4 |
| 267 | GO:0005372 | water transporter activity | 4 | 0.03775689 | -0.6450223 | 4 |
| 268 | GO:0000155 | two-component sensor a | 9 | 0.03776274 | 0.73374991 | 3 |
| 269 | GO:0008586 | imaginal disc-derived wi | 8 | 0.03822779 | -0.2194765 | 9 |
| 270 | GO:0016068 | type I hypersensitivity | 7 | 0.03827775 | 0.86167768 | 2 |
| 271 | GO:0017166 | vinculin binding | 7 | 0.03892001 | 0.9805386 | 1 |
| 272 | GO:0030424 | axon | 5 | 0.03906251 | -0.2371191 | 5 |
| 273 | GO:0016231 | beta-N-acetylglucosamin | 9 | 0.03949965 | 0.72972779 | 3 |
| 274 | GO:0030677 | ribonuclease P complex | 5 | 0.03964077 | 0.85923585 | 2 |
| 275 | GO:0006718 | juvenile hormone biosynt | 5 | 0.03976218 | -0.8590204 | 2 |
| 276 | GO:0016007 | mitochondrial derivative | 6 | 0.03976218 | -0.8590204 | 2 |
| 277 | GO:0048749 | compound eye developm | 9 | 0.03982569 | 0.1417932 | 96 |
| 278 | GO:0007593 | chitin-based cuticle tanni | 6 | 0.04029194 | -0.7279322 | 3 |
| 279 | GO:0006261 | DNA-dependent DNA rep | 9 | 0.04029261 | 0.35913497 | 13 |
| 280 | GO:0004843 | ubiquitin-specific proteas | 9 | 0.04033118 | -0.4423846 | 1 |
| 281 | GO:0007254 | JNK cascade | 6 | 0.04094224 | 0.27421049 | 20 |
| 282 | GO:0046628 | positive regulation of ins | 4 | 0.04102763 | 0.85679402 | 2 |
| 283 | GO:0030259 | lipid glycosylation | 8 | 0.04107425 | 0.9794614 | 1 |
| 284 | GO:0007049 | cell cycle | 5 | 0.04110885 | 0.22586998 | 32 |
| 285 | GO:0008283 | cell proliferation | 6 | 0.04128609 | 0.17100331 | 58 |
| 286 | GO:0030029 | actin filament-based pro | 6 | 0.04156816 | 0.53182471 | 4 |
| 287 | GO:0017154 | semaphorin receptor acti | 6 | 0.04214595 | -0.8548549 | 2 |
| 288 | GO:0005215 | transporter activity | 4 | 0.04241577 | 0.14134914 | 95 |
| 289 | GO:0001104 | #N/A | 6 | 0.0426873 | 0.24139017 | 26 |
| 290 | GO:0005791 | rough endoplasmic reticu | 6 | 0.04281001 | 0.57480066 | 4 |
| 291 | GO:0007030 | Golgi organization | 7 | 0.04289409 | 0.18639466 | 39 |
| 292 | GO:0050804 | regulation of synaptic tra | 7 | 0.04295522 | -0.4931389 | 7 |
| 293 | GO:0000150 | recombinase activity | 7 | 0.04302792 | 0.63539721 | 4 |
| 294 | GO:0045861 | negative regulation of pr | 8 | 0.04330838 | -0.3988502 | 4 |
| 295 | GO:0006930 | substrate-bound cell mig | 7 | 0.04332157 | 0.85284401 | 2 |
| 296 | GO:0070725 | #N/A | 9 | 0.04377096 | -0.5283764 | 6 |
| 297 | GO:0042391 | regulation of membrane | 7 | 0.04378698 | -0.4919278 | 2 |
| 298 | GO:0050658 | RNA transport | 6 | 0.04391545 | -0.8518386 | 2 |
| 299 | GO:0045298 | tubulin complex | 6 | 0.04394157 | 0.43754641 | 7 |
| 300 | GO:0045174 | glutathione dehydrogena | 6 | 0.04453844 | 0.63281138 | 4 |
| 301 | GO:0016782 | transferase activity, trans | 7 | 0.04453844 | 0.63281138 | 4 |
| 302 | GO:0004734 | pyrimidodiazepine syntha | 9 | 0.04453844 | 0.63281138 | 4 |
| 303 | GO:0007390 | germ-band shortening | 5 | 0.04457495 | 0.32307478 | 15 |
| 304 | GO:0008757 | S-adenosylmethionine-de | 6 | 0.04516579 | 0.71737413 | 3 |
| 305 | GO:0016874 | ligase activity | 6 | 0.04533814 | 0.71701501 | 3 |
| 306 | GO:0000049 | tRNA binding | 6 | 0.0454208 | 0.43565424 | 9 |
| 307 | GO:0031473 | myosin III binding | 6 | 0.04624444 | 0.97687612 | 1 |
| 308 | GO:0006810 | transport | 5 | 0.04625047 | 0.14118412 | 93 |
| 309 | GO:0008889 | glycerophosphodiester ph | 6 | 0.04626207 | -0.5246408 | 1 |
| 310 | GO:0046034 | ATP metabolic process | 8 | 0.04642862 | 0.41343777 | 9 |
| 311 | GO:0008431 | vitamin E binding | 7 | 0.04646667 | 0.39518505 | 8 |
| 312 | GO:0008108 | UDP-glucose:hexose-1-ph | 6 | 0.04667528 | 0.97666068 | 1 |
| 313 | GO:0007399 | nervous system developm | 5 | 0.04669696 | 0.1417522 | 92 |
| 314 | GO:0008301 | DNA bending activity | 9 | 0.0467248 | 0.56834997 | 5 |
| 315 | GO:0005542 | folic acid binding | 10 | 0.04688131 | 0.62893262 | 4 |
| 316 | GO:0000221 | vacuolar proton-transport | 4 | 0.0469445 | 0.37875521 | 8 |
| 317 | GO:0004867 | serine-type endopeptidas | 7 | 0.04695711 | -0.1620319 | 37 |
| 318 | GO:0007005 | mitochondrion organizati | 6 | 0.04697853 | 0.26385169 | 22 |
| 319 | GO:0000808 | origin recognition comple | 6 | 0.04724975 | 0.97637343 | 1 |
| 320 | GO:0071805 | #N/A | 8 | 0.04731467 | 0.37835993 | 9 |
| 321 | GO:0008934 | inositol-1(or 4)-monopho | 7 | 0.04749938 | 0.52284483 | 6 |
| 322 | GO:0045179 | apical cortex | 7 | 0.04761282 | -0.2584998 | 4 |
| 323 | GO:0008511 | sodium:potassium:chlorid | 6 | 0.04777283 | -0.627496 | 4 |
| 324 | GO:0003682 | chromatin binding | 6 | 0.04794168 | 0.16271795 | 53 |
| 325 | GO:0001745 | compound eye morphoge | 5 | 0.04805089 | -0.1561274 | 11 |
| 326 | GO:0000941 | inner kinetochore of cond | 6 | 0.04811145 | 0.97594255 | 1 |
| 327 | GO:0000788 | nuclear nucleosome | 5 | 0.04811145 | 0.97594255 | 1 |
| 328 | GO:0004363 | glutathione synthase acti | 6 | 0.0481412 | 0.84487216 | 2 |
| 329 | GO:0009353 | mitochondrial oxoglutar | 3 | 0.04831398 | 0.6266341 | 4 |
| 330 | GO:0031523 | Myb complex | 4 | 0.0484711 | 0.37713813 | 11 |
| 331 | GO:0035249 | synaptic transmission, gl | 4 | 0.04858788 | -0.844154 | 2 |
| 332 | GO:0002102 | podosome | 5 | 0.04882953 | 0.97558348 | 1 |
| 333 | GO:0007202 | activation of phospholipa | 6 | 0.04890179 | 0.84365125 | 2 |
| 334 | GO:0007303 | cytoplasmic transport, nu | 5 | 0.04914411 | 0.32827103 | 14 |
| 335 | GO:0007559 | histolysis | 5 | 0.0492167 | -0.8431485 | 2 |
| 336 | GO:0030898 | actin-dependent ATPase | 6 | 0.04944226 | -0.8427894 | 2 |
| 337 | GO:0007637 | proboscis extension refle | 5 | 0.04950433 | -0.5640974 | 1 |
| 338 | GO:0005786 | signal recognition particl | 5 | 0.04956869 | 0.43062442 | 9 |
| 339 | GO:0008592 | regulation of Toll signalir | 4 | 0.04974458 | 0.56373824 | 4 |

| Table S2_Insulin-Mock |  |  |  |  |  |  |
| --- | --- | --- | --- | --- | --- | --- |
| Rank | Function | Description | Depth | p-value | es score | # genes |
| 1 | GO:0003785 | actin monomer binding | 9 | 4.27E-10 | 1 | 1 |
| 2 | GO:0070449 | elongin complex | 7 | 4.27E-10 | 1 | 1 |
| 3 | GO:0008541 | proteasome regulatory particle, lid subc | 5 | 9.87E-07 | 0.837506 | 8 |
| 4 | GO:0006360 | transcription from RNA polymerase I pr | 5 | 1.24E-06 | 0.907871 | 6 |
| 5 | GO:0005736 | transcription factor activity, RNA polym | 7 | 1.24E-06 | 0.907871 | 6 |
| 6 | GO:0006890 | retrograde vesicle-mediated transport, | 5 | 4.34E-05 | 0.78557 | 7 |
| 7 | GO:0007265 | Ras protein signal transduction | 7 | 5.35E-05 | -0.5046 | 4 |
| 8 | GO:0005634 | nucleus | 8 | 6.54E-05 | -0.09485 | 610 |
| 9 | GO:0000175 | 3'-5'-exoribonuclease activity | 8 | 6.70E-05 | 0.615518 | 12 |
| 10 | GO:0005575 | cellular_component | 8 | 7.52E-05 | 0.087778 | 1119 |
| 11 | GO:0003674 | molecular_function | 6 | 7.56E-05 | 0.086039 | 1117 |
| 12 | GO:0016021 | integral to membrane | 6 | 8.83E-05 | 0.108074 | 507 |
| 13 | GO:0008150 | biological_process | 5 | 8.90E-05 | 0.087048 | 966 |
| 14 | GO:0000502 | proteasome complex | 4 | 9.27E-05 | 0.480288 | 13 |
| 15 | GO:0008270 | zinc ion binding | 5 | 9.89E-05 | 0.094228 | 856 |
| 16 | GO:0005737 | cytoplasm | 6 | 1.22E-04 | 0.117641 | 376 |
| 17 | GO:0005524 | ATP binding | 7 | 1.28E-04 | 0.130701 | 365 |
| 18 | GO:0015030 | Cajal body | 6 | 1.52E-04 | 0.794459 | 6 |
| 19 | GO:0003677 | DNA binding | 6 | 1.53E-04 | -0.14742 | 226 |
| 20 | GO:0022008 | neurogenesis | 7 | 1.58E-04 | 0.165315 | 312 |
| 21 | GO:0016020 | membrane | 6 | 1.59E-04 | 0.119751 | 289 |
| 22 | GO:0009982 | pseudouridine synthase activity | 5 | 1.79E-04 | 0.637483 | 8 |
| 23 | GO:0055114 | oxidation reduction | 6 | 2.04E-04 | 0.20253 | 257 |
| 24 | GO:0000922 | spindle pole | 8 | 2.05E-04 | -0.47282 | 4 |
| 25 | GO:0048081 | positive regulation of cuticle pigmentat | 4 | 2.34E-04 | 0.951168 | 3 |
| 26 | GO:0008535 | respiratory chain complex IV assembly | 5 | 2.41E-04 | 0.657815 | 9 |
| 27 | GO:0006508 | proteolysis | 8 | 2.75E-04 | 0.096633 | 282 |
| 28 | GO:0005875 | microtubule associated complex | 6 | 3.03E-04 | 0.140373 | 168 |
| 29 | GO:0001522 | pseudouridine synthesis | 9 | 3.31E-04 | 0.647384 | 7 |
| 30 | GO:0055085 | transmembrane transport | 6 | 3.48E-04 | 0.13552 | 162 |
| 31 | GO:0005663 | DNA replication factor C complex | 7 | 3.52E-04 | 0.822681 | 5 |
| 32 | GO:0005739 | mitochondrion | 7 | 3.82E-04 | 0.213819 | 140 |
| 33 | GO:0005811 | lipid particle | 5 | 4.48E-04 | -0.14507 | 68 |
| 34 | GO:0000791 | euchromatin | 6 | 4.73E-04 | -0.66871 | 1 |
| 35 | GO:0003684 | damaged DNA binding | 7 | 5.68E-04 | 0.484171 | 12 |
| 36 | GO:0020037 | heme binding | 8 | 6.09E-04 | 0.235756 | 91 |
| 37 | GO:0009055 | electron carrier activity | 4 | 6.45E-04 | 0.177484 | 94 |
| 38 | GO:0006334 | nucleosome assembly | 7 | 6.46E-04 | -0.42723 | 58 |
| 39 | GO:0031427 | response to methotrexate | 4 | 6.82E-04 | 0.569022 | 10 |
| 40 | GO:0006333 | chromatin assembly or disassembly | 5 | 6.94E-04 | -0.43887 | 57 |
| 41 | GO:0000786 | nucleosome | 9 | 7.12E-04 | -0.47821 | 51 |
| 42 | GO:0045727 | positive regulation of translation | 6 | 7.18E-04 | -0.69235 | 1 |
| 43 | GO:0035076 | ecdysone receptor-mediated signaling | 6 | 7.58E-04 | -0.65156 | 8 |
| 44 | GO:0051603 | proteolysis involved in cellular protein c | 8 | 7.73E-04 | 0.542897 | 10 |
| 45 | GO:0007284 | spermatogonial cell division | 7 | 7.75E-04 | 0.859771 | 4 |
| 46 | GO:0030127 | COPII vesicle coat | 7 | 7.87E-04 | 0.926789 | 3 |
| 47 | GO:0005509 | calcium ion binding | 5 | 7.99E-04 | 0.147886 | 113 |
| 48 | GO:0005549 | odorant binding | 8 | 8.00E-04 | -0.21269 | 79 |
| 49 | GO:0005267 | potassium channel activity | 5 | 8.25E-04 | 0.687013 | 7 |
| 50 | GO:0003735 | structural constituent of ribosome | 5 | 9.22E-04 | 0.156176 | 101 |
| 51 | GO:0005215 | transporter activity | 6 | 9.24E-04 | 0.234535 | 63 |
| 52 | GO:0004497 | monooxygenase activity | 6 | 9.48E-04 | 0.499423 | 13 |
| 53 | GO:0016705 | oxidoreductase activity, acting on paire | 6 | 9.49E-04 | 0.275704 | 62 |
| 54 | GO:0005515 | protein binding | 6 | 9.51E-04 | 0.092768 | 468 |
| 55 | GO:0005576 | extracellular region | 4 | 9.54E-04 | 0.100788 | 210 |
| 56 | GO:0003723 | RNA binding | 6 | 9.57E-04 | 0.190764 | 75 |
| 57 | GO:0005832 | chaperonin-containing T-complex | 5 | 9.89E-04 | 0.67987 | 6 |
| 58 | GO:0005792 | microsome | 8 | 9.91E-04 | 0.248916 | 59 |
| 59 | GO:0005840 | ribosome | 7 | 0.001068 | -0.32879 | 8 |
| 60 | GO:0016717 | oxidoreductase activity, acting on paire | 9 | 0.001094 | 0.847147 | 4 |
| 61 | GO:0048015 | phosphoinositide-mediated signaling | 5 | 0.001098 | -0.847 | 4 |
| 62 | GO:0032504 | multicellular organism reproduction | 4 | 0.001111 | -0.30806 | 45 |
| 63 | GO:0005783 | endoplasmic reticulum | 5 | 0.001116 | 0.170919 | 84 |
| 64 | GO:0051082 | unfolded protein binding | 7 | 0.001128 | 0.241642 | 55 |
| 65 | GO:0007026 | negative regulation of microtubule dep | 5 | 0.001144 | -0.71943 | 6 |
| 66 | GO:0005730 | nucleolus | 8 | 0.001263 | 0.325898 | 35 |
| 67 | GO:0003676 | nucleic acid binding | 5 | 0.001279 | 0.09242 | 450 |
| 68 | GO:0007594 | puparial adhesion | 6 | 0.001303 | -0.54687 | 4 |
| 69 | GO:0008333 | endosome to lysosome transport | 7 | 0.001307 | -0.66861 | 7 |
| 70 | GO:0006412 | translation | 8 | 0.001316 | 0.144895 | 106 |
| 71 | GO:0008219 | cell death | 8 | 0.001318 | 0.570686 | 10 |
| 72 | GO:0004175 | endopeptidase activity | 8 | 0.001366 | 0.339285 | 31 |
| 73 | GO:0006555 | methionine metabolic process | 9 | 0.001376 | 0.911786 | 3 |
| 74 | GO:0009408 | response to heat | 8 | 0.001403 | 0.274015 | 45 |
| 75 | GO:0005615 | extracellular space | 7 | 0.001403 | -0.1544 | 121 |
| 76 | GO:0006457 | protein folding | 9 | 0.001467 | 0.184965 | 70 |
| 77 | GO:0016772 | transferase activity, transferring phosph | 4 | 0.0015 | 0.321211 | 42 |
| 78 | GO:0007517 | muscle development | 4 | 0.001511 | -0.27537 | 16 |
| 79 | GO:0008152 | metabolic process | 4 | 0.00157 | 0.128805 | 125 |
| 80 | GO:0022625 | cytosolic large ribosomal subunit | 4 | 0.001665 | -0.41099 | 1 |
| 81 | GO:0042626 | ATPase activity, coupled to transmemb | 7 | 0.001709 | 0.346649 | 38 |
| 82 | GO:0050909 | sensory perception of taste | 5 | 0.001825 | -0.26121 | 38 |
| 83 | GO:0004155 | 6,7-dihydropteridine reductase activity | 7 | 0.002019 | 0.99899 | 1 |
| 84 | GO:0043190 | ATP-binding cassette (ABC) transporter | 6 | 0.002041 | 0.375976 | 33 |
| 85 | GO:0051087 | chaperone binding | 7 | 0.002062 | 0.445645 | 14 |
| 86 | GO:0007298 | border follicle cell migration | 7 | 0.002171 | -0.24044 | 22 |
| 87 | GO:0000177 | cytoplasmic exosome (RNase complex) | 8 | 0.002199 | 0.609947 | 6 |
| 88 | GO:0022627 | cytosolic small ribosomal subunit | 8 | 0.002298 | -0.35152 | 3 |
| 89 | GO:0006810 | transport | 6 | 0.002384 | 0.195022 | 58 |
| 90 | GO:0003682 | chromatin binding | 6 | 0.002457 | -0.23021 | 22 |
| 91 | GO:0004364 | glutathione transferase activity | 7 | 0.002524 | 0.34838 | 28 |
| 92 | GO:0000022 | mitotic spindle elongation | 7 | 0.00253 | -0.21735 | 16 |
| 93 | GO:0031476 | myosin VI complex | 5 | 0.002603 | 0.890868 | 3 |
| 94 | GO:0004984 | olfactory receptor activity | 6 | 0.002637 | -0.24447 | 40 |
| 95 | GO:0008062 | eclosion rhythm | 6 | 0.002669 | 0.570789 | 9 |
| 96 | GO:0042176 | regulation of protein catabolic process | 5 | 0.002715 | 0.735651 | 4 |
| 97 | GO:0035060 | brahma complex | 6 | 0.002761 | -0.54278 | 1 |
| 98 | GO:0003899 | DNA-directed RNA polymerase activity | 4 | 0.002865 | 0.377498 | 25 |
| 99 | GO:0071805 | potassium ion transmembrane transpo | 7 | 0.002927 | 0.496993 | 10 |
| 100 | GO:0006260 | DNA replication | 3 | 0.002968 | 0.283263 | 21 |
| 101 | GO:0004655 | porphobilinogen synthase activity | 6 | 0.003029 | 0.998486 | 1 |
| 102 | GO:0035195 | gene silencing by miRNA | 6 | 0.003043 | -0.4332 | 4 |
| 103 | GO:0006911 | phagocytosis, engulfment | 9 | 0.003194 | -0.13511 | 72 |
| 104 | GO:0030126 | COPI vesicle coat | 6 | 0.003204 | 0.563317 | 7 |
| 105 | GO:0008934 | inositol-1(or 4)-monophosphatase activ | 8 | 0.003268 | 0.726663 | 5 |
| 106 | GO:0003954 | NADH dehydrogenase activity | 9 | 0.00331 | -0.36076 | 9 |
| 107 | GO:0006364 | rRNA processing | 6 | 0.003328 | 0.558029 | 18 |
| 108 | GO:0015986 | ATP synthesis coupled proton transport | 8 | 0.003371 | -0.38672 | 6 |
| 109 | GO:0005839 | proteasome core complex | 7 | 0.003556 | 0.412963 | 18 |
| 110 | GO:0005200 | structural constituent of cytoskeleton | 9 | 0.003577 | -0.28588 | 10 |
| 111 | GO:0019991 | septate junction assembly | 7 | 0.003735 | -0.44427 | 6 |
| 112 | GO:0004298 | threonine-type endopeptidase activity | 8 | 0.003761 | 0.385249 | 18 |
| 113 | GO:0005838 | proteasome regulatory particle | 8 | 0.003788 | 0.512226 | 15 |
| 114 | GO:0005700 | polytene chromosome | 8 | 0.003846 | -0.20728 | 36 |
| 115 | GO:0097033 | obsolete mitochondrial respiratory chain | 8 | 0.003894 | 0.998053 | 1 |
| 116 | GO:0042254 | ribosome biogenesis | 8 | 0.003963 | 0.540902 | 13 |
| 117 | GO:0006396 | RNA processing | 8 | 0.003989 | 0.401885 | 13 |
| 118 | GO:0016336 | establishment or maintenance of polar | 8 | 0.004167 | -0.71469 | 5 |
| 119 | GO:0007606 | sensory perception of chemical stimulus | 9 | 0.004184 | -0.18602 | 66 |
| 120 | GO:0005705 | polytene chromosome interband | 8 | 0.004192 | -0.42471 | 3 |
| 121 | GO:0007030 | Golgi organization | 9 | 0.00425 | 0.243488 | 28 |
| 122 | GO:0033842 | N-acetyl-beta-glucosaminyl-glycoprotein | 4 | 0.004327 | 0.997836 | 1 |
| 123 | GO:0042274 | ribosomal small subunit biogenesis | 8 | 0.004347 | 0.953408 | 2 |
| 124 | GO:0032008 | positive regulation of TOR signaling path | 6 | 0.004361 | 0.870384 | 3 |
| 125 | GO:0030170 | pyridoxal phosphate binding | 7 | 0.004419 | 0.288819 | 28 |
| 126 | GO:0003700 | transcription factor activity | 7 | 0.004512 | 0.091239 | 375 |
| 127 | GO:0006096 | glycolysis | 9 | 0.004726 | 0.370366 | 20 |
| 128 | GO:0006680 | glucosylceramide catabolic process | 7 | 0.00476 | 0.99762 | 1 |
| 129 | GO:0016142 | O-glycoside catabolic process | 7 | 0.00476 | 0.99762 | 1 |
| 130 | GO:0008422 | beta-glucosidase activity | 7 | 0.00476 | 0.99762 | 1 |
| 131 | GO:0008206 | bile acid metabolic process | 6 | 0.00476 | 0.99762 | 1 |
| 132 | GO:0016992 | lipoate synthase activity | 6 | 0.00476 | 0.99762 | 1 |
| 133 | GO:0003331 | positive regulation of extracellular mat | 8 | 0.004788 | 0.9511 | 2 |
| 134 | GO:0034394 | protein localization at cell surface | 8 | 0.004788 | 0.9511 | 2 |
| 135 | GO:0006120 | mitochondrial electron transport, NADH | 5 | 0.004968 | -0.31697 | 11 |
| 136 | GO:0007156 | homophilic cell adhesion | 9 | 0.005027 | 0.348876 | 21 |
| 137 | GO:0006355 | regulation of transcription, DNA-depend | 7 | 0.005129 | 0.091185 | 367 |
| 138 | GO:0005770 | late endosome | 4 | 0.005141 | -0.51774 | 1 |
| 139 | GO:0034511 | U3 snoRNA binding | 9 | 0.005336 | 0.997332 | 1 |
| 140 | GO:0004367 | glycerol-3-phosphate dehydrogenase (N | 9 | 0.005351 | -0.86122 | 3 |
| 141 | GO:0005759 | mitochondrial matrix | 9 | 0.005376 | 0.236999 | 36 |
| 142 | GO:0001661 | conditioned taste aversion | 7 | 0.005435 | -0.8605 | 3 |
| 143 | GO:0005886 | plasma membrane | 9 | 0.005449 | 0.091932 | 357 |
| 144 | GO:0004579 | dolichyl-diphosphooligosaccharide-prot | 6 | 0.00552 | 0.859781 | 3 |
| 145 | GO:0048193 | Golgi vesicle transport | 7 | 0.005625 | 0.997187 | 1 |
| 146 | GO:0005747 | mitochondrial respiratory chain comple | 5 | 0.005645 | -0.26546 | 17 |
| 147 | GO:0005884 | actin filament | 2 | 0.005691 | -0.3966 | 5 |
| 148 | GO:0000166 | nucleotide binding | 4 | 0.005704 | 0.09952 | 159 |
| 149 | GO:0006633 | fatty acid biosynthetic process | 6 | 0.005705 | 0.370665 | 16 |
| 150 | GO:0005097 | Rab GTPase activator activity | 6 | 0.005817 | -0.32695 | 10 |
| 151 | GO:0007440 | foregut morphogenesis | 5 | 0.00591 | 0.48983 | 8 |
| 152 | GO:0004379 | glycylpeptide N-tetradecanoyltransfera | 5 | 0.005913 | 0.997043 | 1 |
| 153 | GO:0006499 | N-terminal protein myristoylation | 7 | 0.005913 | 0.997043 | 1 |
| 154 | GO:0030134 | ER to Golgi transport vesicle | 4 | 0.005915 | 0.696912 | 4 |
| 155 | GO:0048060 | negative gravitaxis | 7 | 0.005945 | 0.766573 | 4 |
| 156 | GO:0005542 | folic acid binding | 8 | 0.005945 | 0.766573 | 4 |
| 157 | GO:0000176 | nuclear exosome (RNase complex) | 7 | 0.005958 | 0.452832 | 8 |
| 158 | GO:0009593 | detection of chemical stimulus | 6 | 0.006262 | -0.25606 | 11 |
| 159 | GO:0004572 | mannosyl-oligosaccharide 1,3-1,6-alpha | 4 | 0.006304 | 0.943887 | 2 |
| 160 | GO:0007405 | neuroblast proliferation | 9 | 0.006337 | -0.38629 | 6 |
| 161 | GO:0000123 | histone acetyltransferase complex | 6 | 0.006485 | -0.39614 | 5 |
| 162 | GO:0043035 | chromatin insulator sequence binding | 6 | 0.006523 | -0.59719 | 7 |
| 163 | GO:0005622 | intracellular | 9 | 0.006544 | 0.091575 | 348 |
| 164 | GO:0004689 | phosphorylase kinase activity | 8 | 0.006634 | 0.996683 | 1 |
| 165 | GO:0006007 | glucose catabolic process | 4 | 0.006634 | 0.996683 | 1 |
| 166 | GO:0045199 | maintenance of epithelial cell apical/ba | 4 | 0.006693 | -0.85048 | 3 |
| 167 | GO:0007058 | spindle assembly involved in female me | 9 | 0.00677 | -0.75885 | 4 |
| 168 | GO:0008444 | CDP-diacylglycerol-glycerol-3-phosphate | 8 | 0.006779 | 0.99661 | 1 |
| 169 | GO:0016457 | dosage compensation complex assemb | 6 | 0.00679 | -0.84975 | 3 |
| 170 | GO:0048009 | insulin-like growth factor receptor signa | 4 | 0.006867 | 0.941435 | 2 |
| 171 | GO:0030097 | hemopoiesis | 6 | 0.00688 | -0.38187 | 5 |
| 172 | GO:0030100 | regulation of endocytosis | 5 | 0.006984 | -0.59387 | 7 |
| 173 | GO:0006820 | anion transport | 6 | 0.007036 | 0.593506 | 7 |
| 174 | GO:0003839 | gamma-glutamylcyclotransferase activi | 6 | 0.007037 | 0.940714 | 2 |
| 175 | GO:0030163 | protein catabolic process | 6 | 0.007066 | 0.380086 | 17 |
| 176 | GO:0017105 | acyl-CoA delta11-desaturase activity | 7 | 0.007067 | 0.996466 | 1 |
| 177 | GO:0003689 | DNA clamp loader activity | 9 | 0.007158 | 0.847086 | 3 |
| 178 | GO:0016339 | calcium-dependent cell-cell adhesion | 4 | 0.00717 | 0.372457 | 14 |
| 179 | GO:0016303 | 1-phosphatidylinositol-3-kinase activity | 4 | 0.007178 | -0.84694 | 3 |
| 180 | GO:0046934 | phosphatidylinositol-4,5-bisphosphate 3-kinase activity | 5 | 0.007178 | -0.84694 | 3 |
| 181 | GO:0035005 | phosphatidylinositol-4-phosphate 3-kinase activity | 5 | 0.007178 | -0.84694 | 3 |
| 182 | GO:0035004 | phosphoinositide 3-kinase activity | 7 | 0.007178 | -0.84694 | 3 |
| 183 | GO:0005214 | structural constituent of chitin-based cuticle | 7 | 0.007293 | 0.145635 | 107 |
| 184 | GO:0005543 | phospholipid binding | 9 | 0.007438 | -0.19629 | 39 |
| 185 | GO:0051018 | protein kinase A binding | 8 | 0.007456 | -0.84499 | 3 |
| 186 | GO:0005852 | eukaryotic translation initiation factor 3 binding | 9 | 0.007537 | -0.40239 | 5 |
| 187 | GO:0010181 | FMN binding | 7 | 0.007539 | 0.479818 | 10 |
| 188 | GO:0051056 | regulation of small GTPase mediated signaling | 6 | 0.007772 | -0.47855 | 2 |
| 189 | GO:0031072 | heat shock protein binding | 7 | 0.007805 | 0.252606 | 30 |
| 190 | GO:0031000 | response to caffeine | 6 | 0.007977 | 0.748756 | 4 |
| 191 | GO:0008250 | oligosaccharyltransferase complex | 6 | 0.007977 | 0.748756 | 4 |
| 192 | GO:0048025 | negative regulation of nuclear mRNA splicing | 9 | 0.00806 | -0.74811 | 4 |
| 193 | GO:0006622 | protein targeting to lysosome | 5 | 0.00806 | -0.74811 | 4 |
| 194 | GO:0019908 | nuclear cyclin-dependent protein kinase activity | 5 | 0.008219 | 0.679455 | 4 |
| 195 | GO:0002781 | antifungal peptide production | 6 | 0.008285 | 0.935665 | 2 |
| 196 | GO:0008527 | taste receptor activity | 5 | 0.008379 | -0.22577 | 37 |
| 197 | GO:0070389 | chaperone cofactor-dependent protein folding | 7 | 0.00851 | 0.9348 | 2 |
| 198 | GO:0043169 | cation binding | 6 | 0.009092 | 0.237547 | 32 |
| 199 | GO:0007411 | axon guidance | 8 | 0.009144 | -0.14246 | 74 |
| 200 | GO:0008946 | oligonucleotidase activity | 7 | 0.009375 | 0.995312 | 1 |
| 201 | GO:0007010 | cytoskeleton organization | 9 | 0.009498 | -0.24427 | 17 |
| 202 | GO:0006408 | snRNA export from nucleus | 4 | 0.009516 | 0.931049 | 2 |
| 203 | GO:0008010 | structural constituent of chitin-based layer | 7 | 0.009642 | 0.249573 | 35 |
| 204 | GO:0070822 | Sin3-type complex | 4 | 0.009861 | -0.46847 | 3 |
| 205 | GO:0016233 | telomere capping | 4 | 0.010015 | -0.48902 | 1 |
| 206 | GO:0019985 | bypass DNA synthesis | 9 | 0.010074 | 0.733824 | 4 |
| 207 | GO:0043248 | proteasome assembly | 9 | 0.010143 | 0.928814 | 2 |
| 208 | GO:0005484 | SNAP receptor activity | 8 | 0.010369 | 0.36017 | 15 |
| 209 | GO:0008476 | protein-tyrosine sulfotransferase activity | 10 | 0.010529 | 0.994735 | 1 |
| 210 | GO:0048813 | dendrite morphogenesis | 5 | 0.010665 | -0.14243 | 51 |
| 211 | GO:0006779 | porphyrin biosynthetic process | 7 | 0.010673 | 0.994663 | 1 |
| 212 | GO:0004853 | uroporphyrinogen decarboxylase activity | 5 | 0.010673 | 0.994663 | 1 |
| 213 | GO:0002814 | negative regulation of biosynthetic process | 8 | 0.01079 | 0.926578 | 2 |
| 214 | GO:0008540 | proteasome regulatory particle, base subunit | 9 | 0.010997 | 0.46377 | 11 |
| 215 | GO:0007216 | metabotropic glutamate receptor signaling | 7 | 0.011182 | 0.822562 | 3 |
| 216 | GO:0042250 | maintenance of polarity of embryonic endoderm | 8 | 0.011538 | 0.99423 | 1 |
| 217 | GO:0031629 | synaptic vesicle fusion to presynaptic membrane | 5 | 0.011568 | -0.82054 | 3 |
| 218 | GO:0051297 | centrosome organization | 8 | 0.011688 | -0.24422 | 8 |
| 219 | GO:0005932 | microtubule basal body | 4 | 0.011795 | -0.65926 | 1 |
| 220 | GO:0005971 | ribonucleoside-diphosphate reductase complex | 6 | 0.011964 | 0.922683 | 2 |
| 221 | GO:0004748 | ribonucleoside-diphosphate reductase activity | 6 | 0.011964 | 0.922683 | 2 |
| 222 | GO:0000124 | SAGA complex | 6 | 0.012217 | -0.3505 | 21 |
| 223 | GO:0004657 | proline dehydrogenase activity | 9 | 0.012259 | 0.99387 | 1 |
| 224 | GO:0007099 | centriole replication | 6 | 0.012294 | -0.35434 | 6 |
| 225 | GO:0000009 | alpha-1,6-mannosyltransferase activity | 8 | 0.012404 | 0.993798 | 1 |
| 226 | GO:0006488 | dolichol-linked oligosaccharide biosynth | 8 | 0.012404 | 0.993798 | 1 |
| 227 | GO:0004849 | uridine kinase activity | 9 | 0.012498 | 0.815854 | 3 |
| 228 | GO:0000235 | astral microtubule | 10 | 0.01259 | -0.71932 | 4 |
| 229 | GO:0006123 | mitochondrial electron transport, cytoch | 6 | 0.012832 | -0.408 | 4 |
| 230 | GO:0001105 | transcription coactivator activity | 7 | 0.012898 | -0.60269 | 1 |
| 231 | GO:0007186 | G-protein coupled receptor protein sign | 3 | 0.012922 | 0.144151 | 92 |
| 232 | GO:0008137 | NADH dehydrogenase (ubiquinone) acti | 7 | 0.013118 | -0.26741 | 16 |
| 233 | GO:0008360 | regulation of cell shape | 4 | 0.013374 | -0.17213 | 42 |
| 234 | GO:0005751 | mitochondrial respiratory chain comple | 6 | 0.013441 | -0.36044 | 6 |
| 235 | GO:0003887 | DNA-directed DNA polymerase activity | 7 | 0.01361 | 0.35999 | 9 |
| 236 | GO:0008568 | microtubule-severing ATPase activity | 7 | 0.013725 | 0.810012 | 3 |
| 237 | GO:0033227 | dsRNA transport | 5 | 0.013761 | -0.32213 | 6 |
| 238 | GO:0030117 | membrane coat | 7 | 0.013769 | 0.650195 | 5 |
| 239 | GO:0005815 | microtubule organizing center | 5 | 0.014495 | -0.55635 | 7 |
| 240 | GO:0030330 | DNA damage response, signal transduc | 5 | 0.014567 | 0.992716 | 1 |
| 241 | GO:0007378 | amnioserosa formation | 5 | 0.014586 | 0.59576 | 5 |
| 242 | GO:0005665 | DNA-directed RNA polymerase II, core c | 5 | 0.014676 | 0.416964 | 11 |
| 243 | GO:0016740 | transferase activity | 8 | 0.014676 | 0.416964 | 11 |
| 244 | GO:0016080 | synaptic vesicle targeting | 8 | 0.014684 | -0.80568 | 3 |
| 245 | GO:0035050 | embryonic heart tube development | 9 | 0.014771 | -0.4167 | 3 |
| 246 | GO:0010043 | response to zinc ion | 8 | 0.014848 | 0.804962 | 3 |
| 247 | GO:0007444 | imaginal disc development | 7 | 0.014871 | 0.280146 | 22 |
| 248 | GO:0004004 | ATP-dependent RNA helicase activity | 6 | 0.015093 | 0.245984 | 27 |
| 249 | GO:0045169 | fusome | 7 | 0.015171 | -0.25565 | 17 |
| 250 | GO:0008066 | glutamate receptor activity | 7 | 0.015444 | 0.520185 | 6 |
| 251 | GO:0010468 | regulation of gene expression | 9 | 0.015649 | 0.33751 | 10 |
| 252 | GO:0030716 | oocyte fate determination | 5 | 0.015773 | -0.801 | 3 |
| 253 | GO:0015238 | drug transporter activity | 5 | 0.015819 | 0.551587 | 5 |
| 254 | GO:0004768 | stearoyl-CoA 9-desaturase activity | 8 | 0.015819 | 0.551587 | 5 |
| 255 | GO:0006431 | methionyl-tRNA aminoacylation | 7 | 0.015955 | 0.91071 | 2 |
| 256 | GO:0004825 | methionine-tRNA ligase activity | 7 | 0.015955 | 0.91071 | 2 |
| 257 | GO:0070971 | endoplasmic reticulum exit site | 7 | 0.016161 | 0.910133 | 2 |
| 258 | GO:0008092 | cytoskeletal protein binding | 6 | 0.016248 | -0.32236 | 8 |
| 259 | GO:0005483 | soluble NSF attachment protein activity | 8 | 0.016263 | 0.70259 | 4 |
| 260 | GO:0009987 | cellular process | 7 | 0.016607 | 0.20566 | 23 |
| 261 | GO:0006163 | purine nucleotide metabolic process | 5 | 0.016711 | 0.908619 | 2 |
| 262 | GO:0097062 | dendritic spine maintenance | 6 | 0.01673 | 0.991634 | 1 |
| 263 | GO:0000309 | nicotinamide-nucleotide adenyllyltransf | 7 | 0.01673 | 0.991634 | 1 |
| 264 | GO:0043625 | delta DNA polymerase complex | 9 | 0.016737 | 0.908547 | 2 |
| 265 | GO:0004725 | protein tyrosine phosphatase activity | 6 | 0.016792 | -0.24928 | 16 |
| 266 | GO:0004766 | spermidine synthase activity | 10 | 0.017135 | 0.907465 | 2 |
| 267 | GO:0007160 | cell-matrix adhesion | 4 | 0.017359 | 0.409884 | 8 |
| 268 | GO:0008440 | inositol trisphosphate 3-kinase activity | 9 | 0.017421 | 0.794287 | 3 |
| 269 | GO:0045544 | gibberellin 20-oxidase activity | 8 | 0.017452 | 0.991274 | 1 |
| 270 | GO:2000377 | regulation of reactive oxygen species m | 7 | 0.017605 | -0.58495 | 6 |
| 271 | GO:0048488 | synaptic vesicle endocytosis | 7 | 0.017637 | -0.28382 | 9 |
| 272 | GO:0006963 | positive regulation of antibacterial pept | 5 | 0.018517 | -0.40712 | 4 |
| 273 | GO:0001744 | optic lobe placode formation | 9 | 0.018669 | -0.6935 | 4 |
| 274 | GO:0004407 | histone deacetylase activity | 5 | 0.018694 | -0.58148 | 1 |
| 275 | GO:0008061 | chitin binding | 5 | 0.018943 | 0.161923 | 52 |
| 276 | GO:0046425 | regulation of JAK-STAT cascade | 6 | 0.019328 | -0.43844 | 3 |
| 277 | GO:0060857 | establishment of glial blood-brain barri | 9 | 0.019397 | -0.39114 | 5 |
| 278 | GO:0007608 | sensory perception of smell | 6 | 0.019474 | -0.16716 | 61 |
| 279 | GO:0003696 | satellite DNA binding | 9 | 0.019692 | -0.68997 | 4 |
| 280 | GO:0008449 | N-acetylglucosamine-6-sulfatase activi | 9 | 0.019713 | -0.78563 | 3 |
| 281 | GO:0018149 | peptide cross-linking | 6 | 0.019938 | 0.90018 | 2 |
| 282 | GO:0008188 | neuropeptide receptor activity | 4 | 0.01998 | 0.227005 | 37 |
| 283 | GO:0000178 | exosome (RNase complex) | 8 | 0.020256 | 0.899387 | 2 |
| 284 | GO:0006281 | DNA repair | 5 | 0.020632 | 0.216569 | 31 |
| 285 | GO:0008553 | hydrogen-exporting ATPase activity, pho | 6 | 0.020658 | -0.21882 | 22 |
| 286 | GO:0006325 | establishment or maintenance of chron | 6 | 0.020866 | -0.28295 | 9 |
| 287 | GO:0006686 | sphingomyelin biosynthetic process | 6 | 0.020913 | 0.989543 | 1 |
| 288 | GO:0000276 | mitochondrial proton-transporting ATP | 4 | 0.020923 | -0.40183 | 3 |
| 289 | GO:0031473 | myosin III binding | 6 | 0.021057 | 0.989471 | 1 |
| 290 | GO:0004590 | orotidine-5'-phosphate decarboxylase a | 6 | 0.021346 | 0.989326 | 1 |
| 291 | GO:0004588 | orotate phosphoribosyltransferase activ | 7 | 0.021346 | 0.989326 | 1 |
| 292 | GO:0044205 | 'de novo' UMP biosynthetic process | 7 | 0.021346 | 0.989326 | 1 |
| 293 | GO:0050770 | regulation of axonogenesis | 7 | 0.021581 | -0.40048 | 4 |
| 294 | GO:0008285 | negative regulation of cell proliferation | 8 | 0.021835 | -0.25921 | 10 |
| 295 | GO:0005371 | tricarboxylate secondary active transme | 7 | 0.021941 | -0.77784 | 3 |
| 296 | GO:0035641 | locomotory exploration behavior | 9 | 0.022067 | 0.988966 | 1 |
| 297 | GO:0061099 | negative regulation of protein tyrosine | 7 | 0.022067 | 0.988966 | 1 |
| 298 | GO:0004674 | protein serine/threonine kinase activity | 6 | 0.022304 | -0.118 | 97 |
| 299 | GO:0008603 | cAMP-dependent protein kinase regulat | 6 | 0.022434 | 0.570666 | 6 |
| 300 | GO:0004032 | aldehyde reductase activity | 6 | 0.022461 | 0.570594 | 6 |
| 301 | GO:0006098 | pentose-phosphate shunt | 7 | 0.022488 | 0.570522 | 6 |
| 302 | GO:0051233 | spindle midzone | 9 | 0.022568 | 0.450855 | 5 |
| 303 | GO:0017146 | N-methyl-D-aspartate selective glutam | 5 | 0.02285 | 0.774812 | 3 |
| 304 | GO:0003713 | transcription coactivator activity | 6 | 0.022992 | -0.25013 | 11 |
| 305 | GO:0035080 | heat shock-mediated polytene chromos | 6 | 0.02313 | 0.530231 | 5 |
| 306 | GO:0016758 | transferase activity, transferring hexosy | 6 | 0.023208 | -0.42985 | 11 |
| 307 | GO:0060968 | obsolete regulation of gene silencing | 6 | 0.023465 | -0.56798 | 1 |
| 308 | GO:0031936 | negative regulation of chromatin silenc | 5 | 0.023465 | -0.56798 | 1 |
| 309 | GO:0007304 | chorion-containing eggshell formation | 6 | 0.02359 | 0.370531 | 12 |
| 310 | GO:0006414 | translational elongation | 8 | 0.02359 | 0.370531 | 12 |
| 311 | GO:0032313 | regulation of Rab GTPase activity | 7 | 0.023764 | -0.28889 | 7 |
| 312 | GO:0070855 | myosin VI head/neck binding | 6 | 0.023858 | 0.890804 | 2 |
| 313 | GO:0000910 | cytokinesis | 5 | 0.024044 | -0.18884 | 30 |
| 314 | GO:0007274 | neuromuscular synaptic transmission | 9 | 0.024092 | -0.27311 | 10 |
| 315 | GO:0043204 | perikaryon | 10 | 0.02423 | 0.987884 | 1 |
| 316 | GO:0045836 | positive regulation of meiosis | 4 | 0.02423 | 0.987884 | 1 |
| 317 | GO:0046959 | habituation | 7 | 0.024317 | -0.67583 | 4 |
| 318 | GO:0006979 | response to oxidative stress | 6 | 0.024319 | 0.200324 | 34 |
| 319 | GO:0051101 | regulation of DNA binding | 6 | 0.02433 | 0.770052 | 3 |
| 320 | GO:0044431 | Golgi apparatus part | 8 | 0.024374 | 0.987812 | 1 |
| 321 | GO:0019207 | kinase regulator activity | 7 | 0.024951 | 0.987523 | 1 |
| 322 | GO:0045900 | negative regulation of translational elongation | 7 | 0.024974 | 0.88828 | 2 |
| 323 | GO:0008033 | tRNA processing | 6 | 0.025011 | 0.409154 | 10 |
| 324 | GO:0007029 | endoplasmic reticulum organization | 6 | 0.025055 | 0.494471 | 5 |
| 325 | GO:0021551 | central nervous system morphogenesis | 5 | 0.025304 | -0.76702 | 3 |
| 326 | GO:0045742 | positive regulation of epidermal growth factor receptor signaling pathway | 6 | 0.025441 | -0.56308 | 1 |
| 327 | GO:0038001 | paracrine signaling | 5 | 0.025469 | 0.766518 | 3 |
| 328 | GO:0008518 | reduced folate carrier activity | 6 | 0.025469 | 0.766518 | 3 |
| 329 | GO:0046427 | positive regulation of JAK-STAT cascade | 3 | 0.025469 | 0.766518 | 3 |
| 330 | GO:0004252 | serine-type endopeptidase activity | 4 | 0.025922 | 0.088608 | 279 |
| 331 | GO:0001578 | microtubule bundle formation | 4 | 0.026094 | 0.523232 | 4 |
| 332 | GO:0051726 | regulation of cell cycle | 5 | 0.026123 | -0.23044 | 13 |
| 333 | GO:0008138 | protein tyrosine/serine/threonine phosphorylation | 6 | 0.026157 | -0.32711 | 5 |
| 334 | GO:0030688 | preribosome, small subunit precursor | 5 | 0.026249 | 0.986874 | 1 |
| 335 | GO:0005978 | glycogen biosynthetic process | 5 | 0.026375 | 0.670291 | 3 |
| 336 | GO:0032880 | regulation of protein localization | 6 | 0.026665 | 0.608354 | 5 |
| 337 | GO:0005791 | rough endoplasmic reticulum | 5 | 0.026665 | 0.608354 | 5 |
| 338 | GO:0031941 | filamentous actin | 5 | 0.026712 | 0.560109 | 4 |
| 339 | GO:0005958 | DNA-dependent protein kinase-DNA ligase complex | 4 | 0.027146 | 0.88352 | 2 |
| 340 | GO:0035091 | phosphoinositide binding | 5 | 0.027387 | -0.30833 | 7 |
| 341 | GO:0005829 | cytosol | 5 | 0.027437 | 0.150922 | 54 |
| 342 | GO:0000070 | mitotic sister chromatid segregation | 5 | 0.027531 | 0.308113 | 12 |
| 343 | GO:0003012 | muscle system process | 5 | 0.027547 | 0.986225 | 1 |
| 344 | GO:0006855 | multidrug transport | 5 | 0.027823 | 0.882077 | 2 |
| 345 | GO:0048854 | brain morphogenesis | 5 | 0.027922 | 0.40403 | 10 |
| 346 | GO:0015276 | ligand-gated ion channel activity | 5 | 0.027949 | -0.19013 | 18 |
| 347 | GO:0016201 | synaptic target inhibition | 6 | 0.027959 | -0.88179 | 2 |
| 348 | GO:0008363 | larval chitin-based cuticle development | 5 | 0.028107 | 0.43985 | 10 |
| 349 | GO:0042654 | ecdysis-triggering hormone receptor activation | 5 | 0.028124 | 0.985937 | 1 |
| 350 | GO:0001708 | cell fate specification | 6 | 0.028268 | 0.314242 | 21 |
| 351 | GO:0045217 | cell-cell junction maintenance | 4 | 0.028269 | 0.985865 | 1 |
| 352 | GO:0070856 | myosin VI light chain binding | 4 | 0.028269 | 0.985865 | 1 |
| 353 | GO:0019749 | cytoskeleton-dependent cytoplasmic transport | 5 | 0.028269 | 0.985865 | 1 |
| 354 | GO:0019773 | proteasome core complex, alpha-subunit | 4 | 0.028563 | 0.419825 | 8 |
| 355 | GO:0006030 | chitin metabolic process | 4 | 0.028751 | 0.160658 | 70 |
| 356 | GO:0006144 | purine base metabolic process | 7 | 0.028887 | 0.879841 | 2 |
| 357 | GO:0005918 | septate junction | 8 | 0.029138 | -0.30582 | 9 |
| 358 | GO:0005774 | vacuolar membrane | 5 | 0.029422 | 0.985288 | 1 |
| 359 | GO:0046686 | response to cadmium ion | 4 | 0.029422 | 0.985288 | 1 |
| 360 | GO:0070574 | cadmium ion transmembrane transport | 4 | 0.029422 | 0.985288 | 1 |
| 361 | GO:0015086 | cadmium ion transmembrane transport | 4 | 0.029422 | 0.985288 | 1 |
| 362 | GO:0048148 | behavioral response to cocaine | 4 | 0.029442 | 0.459606 | 8 |
| 363 | GO:0042335 | cuticle development | 4 | 0.029442 | 0.459606 | 8 |
| 364 | GO:0006338 | chromatin remodeling | 5 | 0.029443 | -0.40153 | 3 |
| 365 | GO:0000049 | tRNA binding | 4 | 0.029481 | 0.459534 | 8 |
| 366 | GO:0008514 | organic anion transmembrane transport | 4 | 0.029481 | 0.459534 | 8 |
| 367 | GO:0030176 | integral to endoplasmic reticulum membrane | 5 | 0.029521 | 0.459462 | 8 |
| 368 | GO:0046622 | positive regulation of organ growth | 5 | 0.029639 | -0.60111 | 1 |
| 369 | GO:0016614 | oxidoreductase activity, acting on CH-O | 4 | 0.02986 | 0.339694 | 11 |
| 370 | GO:0035158 | regulation of tube diameter, open trachea | 4 | 0.030257 | 0.43609 | 7 |
| 371 | GO:0005779 | integral to peroxisomal membrane | 4 | 0.030292 | 0.599596 | 4 |
| 372 | GO:0005099 | Ras GTPase activator activity | 4 | 0.030386 | -0.59938 | 1 |
| 373 | GO:0015367 | oxoglutarate:malate antiporter activity | 4 | 0.030488 | -0.66026 | 1 |
| 374 | GO:0019867 | outer membrane | 7 | 0.03072 | 0.984639 | 1 |
| 375 | GO:0005762 | mitochondrial large ribosomal subunit | 6 | 0.030786 | 0.210664 | 21 |
| 376 | GO:0008306 | associative learning | 7 | 0.031075 | -0.43472 | 3 |
| 377 | GO:0008458 | carnitine O-octanoyltransferase activity | 7 | 0.031297 | 0.98435 | 1 |
| 378 | GO:0042384 | cilium assembly | 7 | 0.031315 | 0.268751 | 25 |
| 379 | GO:0035196 | gene silencing by miRNA, production of | 7 | 0.031437 | -0.87465 | 2 |
| 380 | GO:0008098 | 5S rRNA primary transcript binding | 7 | 0.031442 | 0.984278 | 1 |
| 381 | GO:0017150 | tRNA dihydrouridine synthase activity | 8 | 0.031679 | 0.657578 | 4 |
| 382 | GO:0045793 | positive regulation of cell size | 8 | 0.031752 | -0.3281 | 8 |
| 383 | GO:0009306 | protein secretion | 7 | 0.032281 | 0.267686 | 10 |
| 384 | GO:0008320 | protein transmembrane transporter activity | 8 | 0.032768 | 0.431977 | 7 |
| 385 | GO:0043021 | ribonucleoprotein binding | 7 | 0.032884 | 0.983557 | 1 |
| 386 | GO:0004854 | xanthine dehydrogenase activity | 8 | 0.033012 | 0.871547 | 2 |
| 387 | GO:0008593 | regulation of Notch signaling pathway | 6 | 0.033097 | -0.32645 | 5 |
| 388 | GO:0016585 | chromatin remodeling complex | 6 | 0.03339 | -0.50859 | 1 |
| 389 | GO:0007268 | synaptic transmission | 6 | 0.033402 | -0.20849 | 23 |
| 390 | GO:0016709 | oxidoreductase activity, acting on paired | 6 | 0.033749 | 0.983124 | 1 |
| 391 | GO:0006974 | response to DNA damage stimulus | 6 | 0.03387 | 0.149454 | 26 |
| 392 | GO:0003729 | mRNA binding | 7 | 0.03409 | -0.11051 | 80 |
| 393 | GO:0004222 | metalloendopeptidase activity | 7 | 0.034163 | 0.160198 | 46 |
| 394 | GO:0005687 | snRNP U4 | 6 | 0.034171 | 0.869311 | 2 |
| 395 | GO:0015301 | anion:anion antiporter activity | 8 | 0.034171 | 0.869311 | 2 |
| 396 | GO:0005452 | inorganic anion exchanger activity | 5 | 0.034171 | 0.869311 | 2 |
| 397 | GO:0008079 | translation termination factor activity | 4 | 0.034171 | 0.869311 | 2 |
| 398 | GO:0043044 | ATP-dependent chromatin remodeling | 4 | 0.034575 | -0.65137 | 4 |
| 399 | GO:0031670 | cellular response to nutrient | 4 | 0.034615 | 0.982691 | 1 |
| 400 | GO:0007051 | spindle organization | 4 | 0.034618 | -0.4103 | 3 |
| 401 | GO:0035007 | regulation of melanization defense response | 5 | 0.034625 | -0.86845 | 2 |
| 402 | GO:0000302 | response to reactive oxygen species | 5 | 0.034768 | -0.74098 | 3 |
| 403 | GO:0005488 | binding | 5 | 0.034788 | 0.090749 | 247 |
| 404 | GO:0031571 | G1 DNA damage checkpoint | 4 | 0.035158 | -0.86744 | 2 |
| 405 | GO:0008068 | extracellular-glutamate-gated chloride | 6 | 0.035336 | 0.982331 | 1 |
| 406 | GO:0007431 | salivary gland development | 6 | 0.035361 | 0.29802 | 16 |
| 407 | GO:0005104 | fibroblast growth factor receptor binding | 6 | 0.035529 | 0.739108 | 3 |
| 408 | GO:0035561 | regulation of chromatin binding | 7 | 0.035529 | 0.739108 | 3 |
| 409 | GO:0005096 | GTPase activator activity | 5 | 0.035582 | -0.30761 | 10 |
| 410 | GO:0016044 | membrane organization | 8 | 0.0357 | 0.377719 | 10 |
| 411 | GO:0007561 | imaginal disc eversion | 4 | 0.035867 | -0.47415 | 2 |
| 412 | GO:0004450 | isocitrate dehydrogenase (NADP+) activity | 7 | 0.035913 | 0.982042 | 1 |
| 413 | GO:0006102 | isocitrate metabolic process | 6 | 0.035913 | 0.982042 | 1 |
| 414 | GO:0006097 | glyoxylate cycle | 6 | 0.035913 | 0.982042 | 1 |
| 415 | GO:0005890 | sodium:potassium-exchanging ATPase c | 7 | 0.035953 | -0.54136 | 1 |
| 416 | GO:0016055 | Wnt receptor signaling pathway | 8 | 0.036375 | -0.19051 | 27 |
| 417 | GO:0005744 | mitochondrial inner membrane presequ | 6 | 0.036659 | 0.391026 | 8 |
| 418 | GO:0004307 | ethanolaminephosphotransferase activi | 6 | 0.036778 | 0.98161 | 1 |
| 419 | GO:0019135 | deoxyhypusine monooxygenase activity | 4 | 0.036922 | 0.981538 | 1 |
| 420 | GO:0008630 | DNA damage response, signal transduc | 5 | 0.037069 | 0.447082 | 6 |
| 421 | GO:0045039 | protein import into mitochondrial inner | 4 | 0.037273 | 0.446778 | 7 |
| 422 | GO:0003951 | NAD+ kinase activity | 5 | 0.037417 | -0.73456 | 3 |
| 423 | GO:0005763 | mitochondrial small ribosomal subunit | 3 | 0.037439 | 0.25796 | 27 |
| 424 | GO:0051764 | actin crosslink formation | 7 | 0.037788 | 0.981105 | 1 |
| 425 | GO:0008088 | axon cargo transport | 4 | 0.038021 | -0.42418 | 3 |
| 426 | GO:0008475 | procollagen-lysine 5-dioxygenase activi | 4 | 0.038404 | -0.86145 | 2 |
| 427 | GO:0042391 | regulation of membrane potential | 8 | 0.038485 | -0.49993 | 7 |
| 428 | GO:0002807 | positive regulation of antimicrobial pep | 7 | 0.038509 | 0.980744 | 1 |
| 429 | GO:0050839 | cell adhesion molecule binding | 4 | 0.038559 | 0.329195 | 9 |
| 430 | GO:0046168 | glycerol-3-phosphate catabolic process | 5 | 0.038564 | -0.86116 | 2 |
| 431 | GO:0006826 | iron ion transport | 5 | 0.038622 | -0.53669 | 6 |
| 432 | GO:0008199 | ferric iron binding | 3 | 0.038622 | -0.53669 | 6 |
| 433 | GO:0007349 | cellularization | 5 | 0.038679 | -0.22475 | 14 |
| 434 | GO:0048142 | germarium-derived cystoblast division | 4 | 0.039369 | 0.859719 | 2 |
| 435 | GO:0071632 | optomotor response | 7 | 0.039369 | 0.859719 | 2 |
| 436 | GO:0004353 | glutamate dehydrogenase [NAD(P)+] ac | 6 | 0.039369 | 0.859719 | 2 |
| 437 | GO:0003896 | DNA primase activity | 7 | 0.039369 | 0.859719 | 2 |
| 438 | GO:0008649 | rRNA methyltransferase activity | 8 | 0.039369 | 0.859719 | 2 |
| 439 | GO:0006269 | DNA replication, synthesis of RNA prim | 8 | 0.039369 | 0.859719 | 2 |
| 440 | GO:0019551 | glutamate catabolic process to 2-oxogl | 6 | 0.039369 | 0.859719 | 2 |
| 441 | GO:0043462 | regulation of ATPase activity | 7 | 0.039369 | 0.859719 | 2 |
| 442 | GO:0009328 | phenylalanine-tRNA ligase complex | 6 | 0.03941 | 0.859647 | 2 |
| 443 | GO:0006432 | phenylalanyl-tRNA aminoacylation | 4 | 0.03941 | 0.859647 | 2 |
| 444 | GO:0004826 | phenylalanine-tRNA ligase activity | 4 | 0.03941 | 0.859647 | 2 |
| 445 | GO:0016062 | adaptation of rhodopsin mediated signa | 4 | 0.039475 | 0.64178 | 4 |
| 446 | GO:0046527 | glucosyltransferase activity | 4 | 0.03953 | -0.72966 | 3 |
| 447 | GO:0004644 | phosphoribosylglycinamide formyltrans | 4 | 0.039663 | 0.980167 | 1 |
| 448 | GO:0004641 | phosphoribosylformylglycinamidine cyc | 4 | 0.039663 | 0.980167 | 1 |
| 449 | GO:0004637 | phosphoribosylamine-glycine ligase act | 4 | 0.039663 | 0.980167 | 1 |
| 450 | GO:0004591 | oxoglutarate dehydrogenase (succinyl-t | 4 | 0.039751 | 0.729155 | 3 |
| 451 | GO:0045893 | positive regulation of transcription, DN | 4 | 0.03983 | -0.17053 | 24 |
| 452 | GO:0004115 | 3',5'-cyclic-AMP phosphodiesterase act | 4 | 0.039857 | -0.85885 | 2 |
| 453 | GO:0050918 | positive chemotaxis | 5 | 0.039951 | 0.980023 | 1 |
| 454 | GO:0015992 | proton transport | 4 | 0.040009 | -0.26012 | 13 |
| 455 | GO:0017176 | phosphatidylinositol N-acetylglucosami | 4 | 0.040376 | 0.640121 | 4 |
| 456 | GO:0046530 | photoreceptor cell differentiation | 6 | 0.040511 | -0.8577 | 2 |
| 457 | GO:0034773 | histone H4-K20 trimethylation | 4 | 0.04084 | -0.85712 | 2 |
| 458 | GO:0004730 | pseudouridylate synthase activity | 4 | 0.040905 | 0.726558 | 3 |
| 459 | GO:0004677 | DNA-dependent protein kinase activity | 5 | 0.041105 | 0.979446 | 1 |
| 460 | GO:0043564 | Ku70:Ku80 complex | 4 | 0.041105 | 0.979446 | 1 |
| 461 | GO:0030532 | small nuclear ribonucleoprotein comple | 4 | 0.041168 | 0.209694 | 16 |
| 462 | GO:0048489 | synaptic vesicle transport | 5 | 0.041361 | -0.40126 | 5 |
| 463 | GO:0006072 | glycerol-3-phosphate metabolic process | 5 | 0.041535 | -0.40104 | 5 |
| 464 | GO:0016331 | morphogenesis of embryonic epitheliu | 7 | 0.042121 | -0.4647 | 2 |
| 465 | GO:0022821 | potassium ion antiporter activity | 4 | 0.042691 | 0.978653 | 1 |
| 466 | GO:0015299 | solute:hydrogen antiporter activity | 5 | 0.042691 | 0.978653 | 1 |
| 467 | GO:0003779 | actin binding | 6 | 0.04272 | -0.13139 | 54 |
| 468 | GO:0006606 | protein import into nucleus | 5 | 0.04307 | 0.307683 | 14 |
| 469 | GO:0004556 | alpha-amylase activity | 5 | 0.043179 | 0.721581 | 3 |
| 470 | GO:0009082 | branched chain family amino acid biosy | 8 | 0.043268 | 0.978364 | 1 |
| 471 | GO:0004084 | branched-chain-amino-acid transamina | 8 | 0.043268 | 0.978364 | 1 |
| 472 | GO:0035160 | maintenance of epithelial integrity, ope | 8 | 0.043332 | 0.398849 | 7 |
| 473 | GO:0006897 | endocytosis | 8 | 0.043471 | -0.23038 | 12 |
| 474 | GO:0009267 | cellular response to starvation | 8 | 0.043834 | -0.63401 | 1 |
| 475 | GO:0017133 | mitochondrial electron transfer flavopro | 5 | 0.043862 | -0.85193 | 2 |
| 476 | GO:0033119 | negative regulation of RNA splicing | 6 | 0.043956 | 0.719922 | 3 |
| 477 | GO:0005513 | detection of calcium ion | 7 | 0.043956 | 0.719922 | 3 |
| 478 | GO:0042273 | ribosomal large subunit biogenesis | 7 | 0.044278 | 0.97786 | 1 |
| 479 | GO:0019811 | cocaine binding | 7 | 0.044566 | 0.977715 | 1 |
| 480 | GO:0005330 | dopamine:sodium symporter activity | 7 | 0.044566 | 0.977715 | 1 |
| 481 | GO:0005329 | dopamine transmembrane transporter | 4 | 0.044566 | 0.977715 | 1 |
| 482 | GO:0010669 | epithelial structure maintenance | 6 | 0.044763 | -0.85041 | 2 |
| 483 | GO:0046536 | dosage compensation complex | 5 | 0.045196 | -0.84969 | 2 |
| 484 | GO:0060180 | female mating behavior | 6 | 0.045309 | 0.570625 | 5 |
| 485 | GO:0006537 | glutamate biosynthetic process | 4 | 0.045309 | 0.570625 | 5 |
| 486 | GO:0001558 | regulation of cell growth | 5 | 0.045332 | -0.33184 | 1 |
| 487 | GO:0030133 | transport vesicle | 3 | 0.045398 | 0.57048 | 5 |
| 488 | GO:0060361 | flight | 5 | 0.045398 | 0.57048 | 5 |
| 489 | GO:0008084 | imaginal disc growth factor activity | 5 | 0.045442 | 0.570408 | 5 |
| 490 | GO:0000398 | nuclear mRNA splicing, via spliceosome | 5 | 0.046112 | -0.10186 | 106 |
| 491 | GO:0043565 | sequence-specific DNA binding | 4 | 0.046436 | 0.090702 | 229 |
| 492 | GO:0005818 | aster | 5 | 0.046516 | -0.71458 | 3 |
| 493 | GO:0051298 | centrosome duplication | 7 | 0.046554 | -0.17221 | 16 |
| 494 | GO:0005992 | trehalose biosynthetic process | 5 | 0.046799 | 0.714008 | 3 |
| 495 | GO:0044432 | endoplasmic reticulum part | 6 | 0.046815 | 0.847025 | 2 |
| 496 | GO:0008310 | single-stranded DNA specific 3'-5' exonuclease activity | 4 | 0.046815 | 0.847025 | 2 |
| 497 | GO:0031398 | positive regulation of protein ubiquitination | 5 | 0.046859 | -0.84695 | 2 |
| 498 | GO:0019064 | viral envelope fusion with host membrane | 9 | 0.047018 | 0.976489 | 1 |
| 499 | GO:0019031 | viral envelope | 8 | 0.047018 | 0.976489 | 1 |
| 500 | GO:0046789 | host cell surface receptor binding | 5 | 0.047018 | 0.976489 | 1 |
| 501 | GO:0046328 | regulation of JNK cascade | 6 | 0.047053 | -0.45806 | 2 |
| 502 | GO:0008140 | cAMP response element binding protein binding | 6 | 0.047257 | -0.8463 | 2 |
| 503 | GO:0019992 | diacylglycerol binding | 5 | 0.047328 | -0.26358 | 11 |
| 504 | GO:0016250 | N-sulfoglucosamine sulfohydrolase activity | 6 | 0.047451 | 0.976273 | 1 |
| 505 | GO:0008484 | sulfuric ester hydrolase activity | 6 | 0.047451 | 0.976273 | 1 |
| 506 | GO:0006662 | glycerol ether metabolic process | 6 | 0.047556 | 0.340062 | 14 |
| 507 | GO:0006468 | protein amino acid phosphorylation | 5 | 0.047632 | 0.090976 | 226 |
| 508 | GO:0008145 | phenylalkylamine binding | 6 | 0.047861 | -0.56658 | 5 |
| 509 | GO:0005919 | pleated septate junction | 6 | 0.047925 | -0.84522 | 2 |
| 510 | GO:0021682 | nerve maturation | 6 | 0.047925 | -0.84522 | 2 |
| 511 | GO:0005249 | voltage-gated potassium channel activation | 4 | 0.048242 | -0.30308 | 7 |
| 512 | GO:0045892 | negative regulation of transcription, DNA-templated | 5 | 0.048276 | -0.16616 | 36 |
| 513 | GO:0005507 | copper ion binding | 5 | 0.048807 | 0.310564 | 18 |
| 514 | GO:0007218 | neuropeptide signaling pathway | 5 | 0.048847 | 0.217687 | 33 |
| 515 | GO:0005813 | centrosome | 3 | 0.04898 | -0.1814 | 28 |
| 516 | GO:0005720 | nuclear heterochromatin | 4 | 0.049364 | -0.84291 | 2 |
| 517 | GO:0072375 | medium-term memory | 4 | 0.049513 | -0.62476 | 4 |
| 518 | GO:0034647 | histone demethylase activity (H3-trimethyl lysine) | 4 | 0.049636 | -0.84248 | 2 |
| 519 | GO:0048027 | mRNA 5'-UTR binding | 4 | 0.049682 | -0.84241 | 2 |
| 520 | GO:0005786 | signal recognition particle, endoplasmic reticulum | 4 | 0.049872 | 0.430285 | 5 |

| Table S2_Insulin-KUNV |  |  |  |  |  |  |
| --- | --- | --- | --- | --- | --- | --- |
| Rank | Function | Description | Depth | p-value | es score | # genes |
| 1 | GO:0008946 | oligonucleotidase activity | 9 | 4.0916E-10 | 1 | 1 |
| 2 | GO:0004379 | glycylpeptide N-tetradecanoyltransferase | 7 | 4.0916E-10 | 1 | 1 |
| 3 | GO:0006499 | N-terminal protein myristoylation | 5 | 4.0916E-10 | 1 | 1 |
| 4 | GO:0008541 | proteasome regulatory particle, lid subunit | 5 | 4.541E-08 | 0.89052 | 8 |
| 5 | GO:0000502 | transcription factor activity, RNA polymerase | 7 | 1.3616E-06 | 0.56708 | 14 |
| 6 | GO:0006360 | transcription from RNA polymerase I promoter | 5 | 4.2173E-06 | 0.88692 | 6 |
| 7 | GO:0005736 | DNA-directed RNA polymerase I complex | 7 | 4.2173E-06 | 0.88692 | 6 |
| 8 | GO:0000175 | 3'-5'-exoribonuclease activity | 8 | 5.2769E-06 | 0.67903 | 10 |
| 9 | GO:0005663 | DNA replication factor C complex | 8 | 4.8091E-05 | 0.88094 | 5 |
| 10 | GO:0008535 | respiratory chain complex IV assembly | 8 | 5.0457E-05 | 0.70547 | 9 |
| 11 | GO:0005634 | nucleus | 6 | 6.5664E-05 | -0.106 | 621 |
| 12 | GO:0005575 | cellular_component | 6 | 7.558E-05 | 0.08783 | 1114 |
| 13 | GO:0003674 | molecular_function | 5 | 7.5795E-05 | 0.0861 | 1114 |
| 14 | GO:0005832 | chaperonin-containing T-complex | 4 | 8.0051E-05 | 0.76726 | 7 |
| 15 | GO:0016021 | integral to membrane | 5 | 8.8435E-05 | 0.1187 | 519 |
| 16 | GO:0008150 | biological_process | 6 | 8.9027E-05 | 0.08712 | 965 |
| 17 | GO:0008270 | zinc ion binding | 7 | 9.787E-05 | -0.1079 | 415 |
| 18 | GO:0005737 | cytoplasm | 6 | 0.00012091 | 0.13155 | 389 |
| 19 | GO:0007265 | Ras protein signal transduction | 6 | 0.00012826 | -0.4842 | 3 |
| 20 | GO:0005524 | ATP binding | 7 | 0.0001283 | 0.13158 | 366 |
| 21 | GO:0003677 | DNA binding | 6 | 0.0001537 | -0.1583 | 223 |
| 22 | GO:0022008 | neurogenesis | 5 | 0.00015835 | 0.17793 | 191 |
| 23 | GO:0016020 | membrane | 6 | 0.00015872 | 0.1299 | 295 |
| 24 | GO:0006890 | retrograde vesicle-mediated transport | 8 | 0.00018159 | 0.74119 | 7 |
| 25 | GO:0001522 | pseudouridine synthesis | 4 | 0.0002 | 0.66384 | 8 |
| 26 | GO:0055114 | oxidation reduction | 5 | 0.00020391 | 0.19575 | 256 |
| 27 | GO:0003676 | nucleic acid binding | 8 | 0.00025933 | -0.1079 | 214 |
| 28 | GO:0005875 | microtubule associated complex | 6 | 0.00028542 | 0.15178 | 172 |
| 29 | GO:0055085 | transmembrane transport | 9 | 0.00029146 | 0.18819 | 177 |
| 30 | GO:0005852 | eukaryotic translation initiation factor | 6 | 0.00037691 | -0.4956 | 4 |
| 31 | GO:0000166 | nucleotide binding | 7 | 0.00037721 | 0.12916 | 168 |
| 32 | GO:0005739 | mitochondrion | 7 | 0.00037881 | 0.22829 | 145 |
| 33 | GO:0006412 | translation | 5 | 0.00044801 | 0.23909 | 125 |
| 34 | GO:0003735 | structural constituent of ribosome | 6 | 0.00048166 | 0.26681 | 121 |
| 35 | GO:0005509 | calcium ion binding | 7 | 0.00048171 | 0.16266 | 117 |
| 36 | GO:0048081 | positive regulation of cuticle pigmentation | 8 | 0.00048343 | 0.93777 | 3 |
| 37 | GO:0009982 | pseudouridine synthase activity | 4 | 0.00048578 | 0.60545 | 9 |
| 38 | GO:0005811 | lipid particle | 7 | 0.0004899 | -0.1417 | 70 |
| 39 | GO:0000791 | euchromatin | 4 | 0.00049234 | -0.6672 | 1 |
| 40 | GO:0009055 | electron carrier activity | 5 | 0.00057002 | 0.18797 | 96 |
| 41 | GO:0007026 | negative regulation of microtubule depolymerization | 9 | 0.00059365 | -0.7454 | 6 |
| 42 | GO:0006508 | proteolysis | 6 | 0.00062002 | 0.0893 | 541 |
| 43 | GO:0020037 | heme binding | 6 | 0.00062892 | 0.20363 | 87 |
| 44 | GO:0006334 | nucleosome assembly | 8 | 0.00065088 | -0.4296 | 62 |
| 45 | GO:0030126 | COPI vesicle coat | 7 | 0.0006856 | 0.62231 | 8 |
| 46 | GO:0006333 | chromatin assembly or disassembly | 7 | 0.00069425 | -0.4388 | 57 |
| 47 | GO:0000786 | nucleosome | 5 | 0.00071205 | -0.4781 | 51 |
| 48 | GO:0051603 | proteolysis involved in cellular protein | 8 | 0.00075414 | 0.54372 | 8 |
| 49 | GO:0048015 | phosphoinositide-mediated signaling | 5 | 0.00079563 | -0.8589 | 4 |
| 50 | GO:0005549 | odorant binding | 5 | 0.00079703 | -0.2153 | 78 |
| 51 | GO:0015030 | Cajal body | 6 | 0.00088506 | 0.7298 | 6 |
| 52 | GO:0005515 | protein binding | 6 | 0.00095475 | 0.09284 | 467 |
| 53 | GO:0016705 | oxidoreductase activity, acting on paired | 6 | 0.00097245 | 0.25028 | 60 |
| 54 | GO:0005215 | transporter activity | 6 | 0.00098013 | 0.22814 | 62 |
| 55 | GO:0006810 | transport | 4 | 0.00103554 | 0.22473 | 61 |
| 56 | GO:0051082 | unfolded protein binding | 6 | 0.00106597 | 0.25136 | 56 |
| 57 | GO:0005840 | ribosome | 5 | 0.001068 | -0.3185 | 9 |
| 58 | GO:0005792 | microsome | 8 | 0.00106904 | 0.23487 | 58 |
| 59 | GO:0032504 | multicellular organism reproduction | 7 | 0.00111219 | -0.3013 | 44 |
| 60 | GO:0019773 | proteasome core complex, alpha-subunit | 9 | 0.00113721 | 0.55164 | 9 |
| 61 | GO:0003723 | RNA binding | 5 | 0.00118742 | 0.18317 | 75 |
| 62 | GO:0005730 | nucleolus | 4 | 0.001262 | 0.34652 | 37 |
| 63 | GO:0031427 | response to methotrexate | 5 | 0.00128572 | 0.54735 | 10 |
| 64 | GO:0005700 | polytene chromosome | 7 | 0.001293 | -0.2489 | 33 |
| 65 | GO:0007594 | puparial adhesion | 5 | 0.00130697 | -0.5468 | 4 |
| 66 | GO:0045727 | positive regulation of translation | 8 | 0.00133417 | -0.63 | 8 |
| 67 | GO:0006457 | protein folding | 5 | 0.00135618 | 0.1878 | 70 |
| 68 | GO:0004175 | endopeptidase activity | 6 | 0.00136521 | 0.3936 | 41 |
| 69 | GO:0007517 | muscle development | 7 | 0.00137422 | -0.2944 | 17 |
| 70 | GO:0008219 | cell death | 8 | 0.00141475 | 0.56808 | 10 |
| 71 | GO:0048542 | lymph gland development | 8 | 0.00149037 | -0.4697 | 2 |
| 72 | GO:0016772 | transferase activity, transferring phosph | 8 | 0.00150255 | 0.31859 | 42 |
| 73 | GO:0008360 | regulation of cell shape | 9 | 0.00153062 | -0.2174 | 42 |
| 74 | GO:0050909 | sensory perception of taste | 8 | 0.00160393 | -0.2718 | 38 |
| 75 | GO:0005576 | extracellular region | 7 | 0.00165335 | 0.09657 | 209 |
| 76 | GO:0022625 | cytosolic large ribosomal subunit | 9 | 0.00166546 | -0.3935 | 2 |
| 77 | GO:0042626 | ATPase activity, coupled to transmembr | 4 | 0.00170177 | 0.36451 | 39 |
| 78 | GO:0007298 | border follicle cell migration | 4 | 0.00172096 | -0.2486 | 8 |
| 79 | GO:0007052 | mitotic spindle organization | 4 | 0.00177757 | 0.14201 | 104 |
| 80 | GO:0051056 | regulation of small GTPase mediated s | 4 | 0.00183028 | -0.5348 | 3 |
| 81 | GO:0006260 | DNA replication | 7 | 0.00190797 | 0.33401 | 21 |
| 82 | GO:0030117 | membrane coat | 5 | 0.00199002 | 0.75032 | 5 |
| 83 | GO:0005762 | mitochondrial large ribosomal subunit | 7 | 0.00199191 | 0.30569 | 33 |
| 84 | GO:0008152 | metabolic process | 6 | 0.00203025 | 0.12652 | 124 |
| 85 | GO:0043190 | ATP-binding cassette (ABC) transporte | 7 | 0.00204307 | 0.37336 | 33 |
| 86 | GO:0030127 | COPII vesicle coat | 7 | 0.00215362 | 0.89756 | 3 |
| 87 | GO:0051087 | chaperone binding | 8 | 0.00223587 | 0.4431 | 14 |
| 88 | GO:0022627 | cytosolic small ribosomal subunit | 8 | 0.00228347 | -0.3541 | 3 |
| 89 | GO:0016717 | oxidoreductase activity, acting on paired | 6 | 0.0022941 | 0.81604 | 4 |
| 90 | GO:0017150 | tRNA dihydrouridine synthase activity | 6 | 0.0022941 | 0.81604 | 4 |
| 91 | GO:0000022 | mitotic spindle elongation | 7 | 0.00230051 | -0.22 | 16 |
| 92 | GO:0005622 | intracellular | 7 | 0.00233716 | -0.0997 | 160 |
| 93 | GO:0004364 | glutathione transferase activity | 5 | 0.00236422 | 0.37361 | 29 |
| 94 | GO:0004984 | olfactory receptor activity | 6 | 0.00241795 | -0.25 | 39 |
| 95 | GO:0007030 | Golgi organization | 6 | 0.00254462 | 0.26112 | 24 |
| 96 | GO:0005615 | extracellular space | 5 | 0.00255 | -0.1463 | 46 |
| 97 | GO:0008333 | endosome to lysosome transport | 6 | 0.00258937 | -0.6843 | 6 |
| 98 | GO:0006338 | chromatin remodeling | 4 | 0.00260518 | -0.5012 | 1 |
| 99 | GO:0000922 | spindle pole | 7 | 0.00264962 | -0.4035 | 4 |
| 100 | GO:0003899 | DNA-directed RNA polymerase activity | 3 | 0.00274637 | 0.39806 | 19 |
| 101 | GO:0005200 | structural constituent of cytoskeleton | 6 | 0.00276609 | -0.3044 | 17 |
| 102 | GO:0003954 | NADH dehydrogenase activity | 6 | 0.00282042 | -0.3831 | 6 |
| 103 | GO:0000049 | tRNA binding | 9 | 0.00285108 | 0.56811 | 9 |
| 104 | GO:0016992 | lipoate synthase activity | 6 | 0.00288704 | 0.99856 | 1 |
| 105 | GO:0005763 | mitochondrial small ribosomal subunit | 8 | 0.00290197 | 0.42299 | 24 |
| 106 | GO:0008250 | oligosaccharyltransferase complex | 9 | 0.00299185 | 0.80341 | 4 |
| 107 | GO:0006355 | regulation of transcription, DNA-depend | 6 | 0.00300867 | -0.0958 | 107 |
| 108 | GO:0048813 | dendrite morphogenesis | 8 | 0.00307388 | -0.1604 | 57 |
| 109 | GO:0000176 | nuclear exosome (RNase complex) | 7 | 0.003082 | 0.47702 | 9 |
| 110 | GO:0000177 | cytoplasmic exosome (RNase complex) | 9 | 0.00309586 | 0.59553 | 6 |
| 111 | GO:0006963 | positive regulation of antibacterial pep | 7 | 0.00312074 | -0.4766 | 3 |
| 112 | GO:0042176 | regulation of protein catabolic process | 8 | 0.00312645 | 0.72881 | 4 |
| 113 | GO:0005097 | Rab GTPase activator activity | 8 | 0.00317878 | -0.3785 | 8 |
| 114 | GO:0006364 | rRNA processing | 8 | 0.00332777 | 0.54725 | 17 |
| 115 | GO:0004298 | threonine-type endopeptidase activity | 8 | 0.00337557 | 0.41105 | 14 |
| 116 | GO:0003682 | chromatin binding | 8 | 0.00338955 | -0.2213 | 26 |
| 117 | GO:0003689 | DNA clamp loader activity | 8 | 0.00339105 | 0.88081 | 3 |
| 118 | GO:0006120 | mitochondrial electron transport, NADH | 8 | 0.00340603 | -0.3341 | 8 |
| 119 | GO:0005839 | proteasome core complex | 9 | 0.00341954 | 0.4326 | 14 |
| 120 | GO:0005770 | late endosome | 8 | 0.00342713 | -0.5342 | 1 |
| 121 | GO:0031476 | myosin VI complex | 9 | 0.00350311 | 0.87951 | 3 |
| 122 | GO:0016336 | establishment or maintenance of polar | 4 | 0.0035928 | -0.722 | 5 |
| 123 | GO:0004655 | porphobilinogen synthase activity | 8 | 0.00360881 | 0.9982 | 1 |
| 124 | GO:0006096 | glycolysis | 6 | 0.00361568 | 0.40782 | 21 |
| 125 | GO:0019992 | diacylglycerol binding | 7 | 0.00373121 | -0.3865 | 8 |
| 126 | GO:0004579 | dolichyl-diphosphooligosaccharide-pro | 7 | 0.00374115 | 0.87684 | 3 |
| 127 | GO:0006259 | DNA metabolic process | 9 | 0.00374345 | 0.72 | 5 |
| 128 | GO:0005761 | mitochondrial ribosome | 7 | 0.00374345 | 0.72 | 5 |
| 129 | GO:0005838 | proteasome regulatory particle | 7 | 0.00378707 | 0.51361 | 14 |
| 130 | GO:0008603 | cAMP-dependent protein kinase regula | 7 | 0.0038589 | 0.66568 | 6 |
| 131 | GO:0015986 | ATP synthesis coupled proton transport | 6 | 0.00386685 | -0.3826 | 5 |
| 132 | GO:0008113 | peptide-methionine-(S)-S-oxide reduct | 6 | 0.00389751 | 0.99805 | 1 |
| 133 | GO:0042254 | ribosome biogenesis | 8 | 0.00397004 | 0.51669 | 13 |
| 134 | GO:0009408 | response to heat | 8 | 0.00398393 | 0.2243 | 42 |
| 135 | GO:0019991 | septate junction assembly | 5 | 0.00399077 | -0.4181 | 7 |
| 136 | GO:0007606 | sensory perception of chemical stimulus | 9 | 0.00420247 | -0.1859 | 66 |
| 137 | GO:0019908 | nuclear cyclin-dependent protein kinase activity | 7 | 0.00431662 | 0.71292 | 4 |
| 138 | GO:0005705 | polytene chromosome interband | 4 | 0.00434025 | -0.4361 | 4 |
| 139 | GO:0005104 | fibroblast growth factor receptor binding | 9 | 0.00435736 | 0.87042 | 3 |
| 140 | GO:0006555 | methionine metabolic process | 9 | 0.00436464 | 0.87034 | 3 |
| 141 | GO:0048009 | insulin-like growth factor receptor signaling | 9 | 0.00442259 | 0.95301 | 2 |
| 142 | GO:0043035 | chromatin insulator sequence binding | 7 | 0.0045961 | -0.6138 | 7 |
| 143 | GO:0007284 | spermatogonial cell division | 9 | 0.0046528 | 0.78045 | 4 |
| 144 | GO:0003700 | transcription factor activity | 6 | 0.00468619 | 0.0913 | 372 |
| 145 | GO:0009593 | detection of chemical stimulus | 7 | 0.00487824 | -0.2638 | 11 |
| 146 | GO:0009987 | cellular process | 5 | 0.00489038 | 0.23505 | 18 |
| 147 | GO:0000276 | mitochondrial proton-transporting ATPase activity | 2 | 0.00489826 | -0.4602 | 3 |
| 148 | GO:0042274 | ribosomal small subunit biogenesis | 4 | 0.00493919 | 0.95034 | 2 |
| 149 | GO:0008285 | negative regulation of cell proliferation | 6 | 0.00500335 | -0.3112 | 10 |
| 150 | GO:0007411 | axon guidance | 6 | 0.00517943 | -0.1499 | 72 |
| 151 | GO:0032154 | cleavage furrow | 5 | 0.00529028 | -0.5719 | 8 |
| 152 | GO:0006826 | iron ion transport | 5 | 0.00530283 | -0.65 | 6 |
| 153 | GO:0008199 | ferric iron binding | 7 | 0.00530283 | -0.65 | 6 |
| 154 | GO:0005886 | plasma membrane | 4 | 0.0053077 | 0.09202 | 358 |
| 155 | GO:0004725 | protein tyrosine phosphatase activity | 7 | 0.00534575 | -0.2815 | 11 |
| 156 | GO:0008934 | inositol-1(or 4)-monophosphatase activity | 8 | 0.00536825 | 0.64942 | 5 |
| 157 | GO:0030097 | hemopoiesis | 7 | 0.0054839 | -0.4001 | 4 |
| 158 | GO:0045199 | maintenance of epithelial cell apical/basolateral polarity | 6 | 0.00550023 | -0.8599 | 3 |
| 159 | GO:0007010 | cytoskeleton organization | 4 | 0.00550273 | -0.26 | 21 |
| 160 | GO:0016787 | hydrolase activity | 9 | 0.00555096 | -0.2296 | 20 |
| 161 | GO:0004572 | mannosyl-oligosaccharide 1,3-1,6-alpha-glucosyltransferase activity | 6 | 0.00562124 | 0.94702 | 2 |
| 162 | GO:0016303 | 1-phosphatidylinositol-3-kinase activity | 6 | 0.00563737 | -0.8588 | 3 |
| 163 | GO:0046934 | phosphatidylinositol-4,5-bisphosphate 3-kinase activity | 9 | 0.00563737 | -0.8588 | 3 |
| 164 | GO:0035005 | phosphatidylinositol-4-phosphate 3-kinase activity | 8 | 0.00563737 | -0.8588 | 3 |
| 165 | GO:0035004 | phosphoinositide 3-kinase activity | 4 | 0.00563737 | -0.8588 | 3 |
| 166 | GO:0016063 | rhodopsin biosynthetic process | 4 | 0.00574396 | 0.56814 | 8 |
| 167 | GO:0003785 | actin monomer binding | 9 | 0.00577409 | 0.99711 | 1 |
| 168 | GO:0030134 | ER to Golgi transport vesicle | 8 | 0.0058317 | 0.69762 | 4 |
| 169 | GO:0006431 | methionyl-tRNA aminoacylation | 6 | 0.00583757 | 0.946 | 2 |
| 170 | GO:0004825 | methionine-tRNA ligase activity | 4 | 0.00583757 | 0.946 | 2 |
| 171 | GO:0005542 | folic acid binding | 6 | 0.00589192 | 0.76709 | 4 |
| 172 | GO:0032313 | regulation of Rab GTPase activity | 5 | 0.00593461 | -0.3465 | 8 |
| 173 | GO:0050770 | regulation of axonogenesis | 6 | 0.00598747 | -0.4372 | 2 |
| 174 | GO:0005267 | potassium channel activity | 6 | 0.00599851 | 0.60124 | 7 |
| 175 | GO:0043204 | perikaryon | 6 | 0.00606279 | 0.99697 | 1 |
| 176 | GO:0045836 | positive regulation of meiosis | 7 | 0.00606279 | 0.99697 | 1 |
| 177 | GO:0006633 | fatty acid biosynthetic process | 9 | 0.00620064 | 0.36805 | 16 |
| 178 | GO:0034511 | U3 snoRNA binding | 4 | 0.0063515 | 0.99682 | 1 |
| 179 | GO:0006622 | protein targeting to lysosome | 4 | 0.00636674 | -0.8529 | 3 |
| 180 | GO:0006396 | RNA processing | 5 | 0.00640186 | 0.35814 | 12 |
| 181 | GO:0005747 | mitochondrial respiratory chain complex | 5 | 0.00640686 | -0.2584 | 14 |
| 182 | GO:0000124 | SAGA complex | 7 | 0.00644438 | -0.3865 | 3 |
| 183 | GO:0006281 | DNA repair | 7 | 0.00651523 | 0.24922 | 32 |
| 184 | GO:0003684 | damaged DNA binding | 9 | 0.00674217 | 0.40633 | 13 |
| 185 | GO:0030100 | regulation of endocytosis | 8 | 0.00680123 | -0.5952 | 7 |
| 186 | GO:0043248 | proteasome assembly | 9 | 0.0070832 | 0.94052 | 2 |
| 187 | GO:0030170 | pyridoxal phosphate binding | 7 | 0.00709299 | 0.2683 | 28 |
| 188 | GO:0051018 | protein kinase A binding | 6 | 0.00709583 | -0.8475 | 3 |
| 189 | GO:0043625 | delta DNA polymerase complex | 7 | 0.00711761 | 0.94037 | 2 |
| 190 | GO:0032008 | positive regulation of TOR signaling pathway | 6 | 0.00740237 | 0.84537 | 3 |
| 191 | GO:0048025 | negative regulation of nuclear mRNA splicing | 6 | 0.00747632 | -0.7528 | 4 |
| 192 | GO:0001105 | #N/A | 9 | 0.00760009 | -0.6314 | 1 |
| 193 | GO:0031072 | heat shock protein binding | 5 | 0.00771002 | 0.25597 | 21 |
| 194 | GO:0003331 | #N/A | 5 | 0.00776914 | 0.9377 | 2 |
| 195 | GO:0034394 | protein localization at cell surface | 6 | 0.00776914 | 0.9377 | 2 |
| 196 | GO:0060968 | #N/A | 5 | 0.00794991 | -0.6291 | 1 |
| 197 | GO:0008540 | proteasome regulatory particle, base subunit | 7 | 0.00802349 | 0.47721 | 10 |
| 198 | GO:0010181 | FMN binding | 6 | 0.00802349 | 0.47721 | 10 |
| 199 | GO:0045039 | protein import into mitochondrial inner membrane | 8 | 0.00804708 | 0.52348 | 7 |
| 200 | GO:0016457 | dosage compensation complex assembly | 7 | 0.00819599 | -0.84 | 3 |
| 201 | GO:0004497 | monooxygenase activity | 9 | 0.00823917 | 0.42524 | 12 |
| 202 | GO:0004367 | glycerol-3-phosphate dehydrogenase (NAD) | 4 | 0.0083298 | -0.8392 | 3 |
| 203 | GO:0008527 | taste receptor activity | 7 | 0.00840691 | -0.2257 | 37 |
| 204 | GO:0000235 | astral microtubule | 4 | 0.00842713 | -0.7453 | 4 |
| 205 | GO:0005751 | mitochondrial respiratory chain complex | 4 | 0.00854022 | -0.3763 | 5 |
| 206 | GO:0005884 | actin filament | 9 | 0.00889479 | -0.3664 | 4 |
| 207 | GO:0008440 | inositol trisphosphate 3-kinase activity | 9 | 0.00900891 | 0.8349 | 3 |
| 208 | GO:0019731 | antibacterial humoral response | 8 | 0.00947162 | -0.3458 | 1 |
| 209 | GO:0008033 | tRNA processing | 10 | 0.00950461 | 0.45132 | 8 |
| 210 | GO:0001661 | conditioned taste aversion | 5 | 0.00964959 | -0.8311 | 3 |
| 211 | GO:0004674 | protein serine/threonine kinase activity | 7 | 0.00966002 | -0.1293 | 88 |
| 212 | GO:0060857 | establishment of glial blood-brain barrier | 5 | 0.00987136 | -0.4183 | 5 |
| 213 | GO:0008458 | carnitine O-octanoyltransferase activity | 8 | 0.01010466 | 0.99495 | 1 |
| 214 | GO:0050660 | FAD binding | 9 | 0.01017967 | 0.21325 | 23 |
| 215 | GO:0008156 | negative regulation of DNA replication | 7 | 0.01053941 | 0.57322 | 7 |
| 216 | GO:0005971 | ribonucleoside-diphosphate reductase | 8 | 0.01066056 | 0.92702 | 2 |
| 217 | GO:0004748 | ribonucleoside-diphosphate reductase | 5 | 0.01066056 | 0.92702 | 2 |
| 218 | GO:0008593 | regulation of Notch signaling pathway | 8 | 0.01074757 | -0.3683 | 6 |
| 219 | GO:0004155 | 6,7-dihydropteridine reductase activity | 4 | 0.01082642 | 0.99459 | 1 |
| 220 | GO:0008449 | N-acetylglucosamine-6-sulfatase activity | 6 | 0.01092632 | -0.8239 | 3 |
| 221 | GO:0006408 | snRNA export from nucleus | 6 | 0.01100049 | 0.92586 | 2 |
| 222 | GO:0035060 | brahma complex | 6 | 0.0112034 | -0.484 | 3 |
| 223 | GO:0003887 | DNA-directed DNA polymerase activity | 9 | 0.01125281 | 0.36673 | 9 |
| 224 | GO:0005783 | endoplasmic reticulum | 6 | 0.01149553 | 0.13622 | 79 |
| 225 | GO:0007608 | sensory perception of smell | 8 | 0.01154901 | -0.178 | 59 |
| 226 | GO:0000811 | GINS complex | 8 | 0.01160385 | 0.72464 | 4 |
| 227 | GO:0015238 | drug transporter activity | 9 | 0.01163075 | 0.5681 | 7 |
| 228 | GO:0019843 | rRNA binding | 10 | 0.01166283 | 0.56795 | 7 |
| 229 | GO:0007448 | anterior/posterior pattern formation, i | 6 | 0.01175954 | -0.5675 | 7 |
| 230 | GO:0003839 | gamma-glutamylcyclotransferase activ | 7 | 0.01196279 | 0.92269 | 2 |
| 231 | GO:0002814 | negative regulation of biosynthetic pro | 3 | 0.01200749 | 0.92254 | 2 |
| 232 | GO:0008138 | protein tyrosine/serine/threonine phos | 7 | 0.01204425 | -0.3464 | 4 |
| 233 | GO:0042742 | defense response to bacterium | 4 | 0.01209124 | -0.2661 | 6 |
| 234 | GO:0016080 | synaptic vesicle targeting | 6 | 0.01234009 | -0.8166 | 3 |
| 235 | GO:0018149 | peptide cross-linking | 7 | 0.01236809 | 0.92139 | 2 |
| 236 | GO:0005932 | microtubule basal body | 7 | 0.01240264 | -0.6563 | 1 |
| 237 | GO:0004689 | phosphorylase kinase activity | 5 | 0.01241429 | 0.99379 | 1 |
| 238 | GO:0006007 | glucose catabolic process | 7 | 0.01241429 | 0.99379 | 1 |
| 239 | GO:0006979 | response to oxidative stress | 5 | 0.01243004 | 0.2163 | 35 |
| 240 | GO:0016591 | DNA-directed RNA polymerase II, holo | 5 | 0.01247168 | 0.81598 | 3 |
| 241 | GO:0008513 | secondary active organic cation transm | 5 | 0.01250339 | 0.34108 | 17 |
| 242 | GO:0015035 | protein disulfide oxidoreductase activit | 5 | 0.01263992 | 0.3406 | 20 |
| 243 | GO:0044431 | Golgi apparatus part | 8 | 0.012703 | 0.99365 | 1 |
| 244 | GO:0003696 | satellite DNA binding | 8 | 0.01277811 | -0.7184 | 4 |
| 245 | GO:0007391 | dorsal closure | 9 | 0.01314881 | -0.1735 | 43 |
| 246 | GO:0035641 | #N/A | 8 | 0.01342476 | 0.99329 | 1 |
| 247 | GO:0061099 | #N/A | 7 | 0.01342476 | 0.99329 | 1 |
| 248 | GO:0050954 | sensory perception of mechanical stim | 6 | 0.01354982 | -0.7145 | 4 |
| 249 | GO:0006468 | protein amino acid phosphorylation | 7 | 0.01357864 | -0.1054 | 132 |
| 250 | GO:0010468 | regulation of gene expression | 7 | 0.01364383 | 0.34221 | 14 |
| 251 | GO:0004657 | proline dehydrogenase activity | 9 | 0.01385781 | 0.99307 | 1 |
| 252 | GO:0071805 | #N/A | 5 | 0.01411924 | 0.43465 | 10 |
| 253 | GO:0031122 | cytoplasmic microtubule organization | 5 | 0.01436328 | 0.43391 | 8 |
| 254 | GO:0007405 | neuroblast proliferation | 8 | 0.01452856 | -0.3576 | 6 |
| 255 | GO:0009306 | protein secretion | 7 | 0.01467725 | 0.29543 | 12 |
| 256 | GO:0005779 | integral to peroxisomal membrane | 7 | 0.01491368 | 0.64542 | 4 |
| 257 | GO:0046425 | regulation of JAK-STAT cascade | 7 | 0.01521955 | -0.4494 | 3 |
| 258 | GO:0051101 | regulation of DNA binding | 6 | 0.01521962 | 0.80335 | 3 |
| 259 | GO:0008553 | hydrogen-exporting ATPase activity, ph | 8 | 0.01537196 | -0.2264 | 21 |
| 260 | GO:0048148 | behavioral response to cocaine | 7 | 0.01537945 | 0.49285 | 9 |
| 261 | GO:0005743 | mitochondrial inner membrane | 5 | 0.01551128 | 0.197 | 39 |
| 262 | GO:0048193 | Golgi vesicle transport | 6 | 0.01559004 | 0.9922 | 1 |
| 263 | GO:0008168 | methyltransferase activity | 7 | 0.01567383 | 0.26976 | 16 |
| 264 | GO:0045217 | cell-cell junction maintenance | 9 | 0.01587874 | 0.99206 | 1 |
| 265 | GO:0070856 | #N/A | 6 | 0.01587874 | 0.99206 | 1 |
| 266 | GO:0019749 | cytoskeleton-dependent cytoplasmic tr | 10 | 0.01587874 | 0.99206 | 1 |
| 267 | GO:0003012 | muscle system process | 4 | 0.0160231 | 0.99199 | 1 |
| 268 | GO:0032956 | regulation of actin cytoskeleton organi | 9 | 0.01602838 | -0.5509 | 1 |
| 269 | GO:0004590 | orotidine-5'-phosphate decarboxylase a | 8 | 0.0163118 | 0.99184 | 1 |
| 270 | GO:0004588 | orotate phosphoribosyltransferase acti | 7 | 0.0163118 | 0.99184 | 1 |
| 271 | GO:0044205 | #N/A | 7 | 0.0163118 | 0.99184 | 1 |
| 272 | GO:0004879 | ligand-dependent nuclear receptor acti | 5 | 0.01631867 | -0.3375 | 6 |
| 273 | GO:0022821 | potassium ion antiporter activity | 9 | 0.01645615 | 0.99177 | 1 |
| 274 | GO:0015299 | solute:hydrogen antiporter activity | 5 | 0.01645615 | 0.99177 | 1 |
| 275 | GO:0019135 | deoxyhypusine monooxygenase activity | 5 | 0.01660051 | 0.9917 | 1 |
| 276 | GO:0005543 | phospholipid binding | 6 | 0.01676754 | -0.1789 | 44 |
| 277 | GO:0006914 | autophagy | 9 | 0.01690894 | -0.3618 | 6 |
| 278 | GO:0005890 | sodium:potassium-exchanging ATPase | 6 | 0.01724543 | -0.5862 | 1 |
| 279 | GO:0002781 | antifungal peptide production | 9 | 0.01740642 | 0.90674 | 2 |
| 280 | GO:0005992 | trehalose biosynthetic process | 9 | 0.01759554 | 0.7936 | 3 |
| 281 | GO:0005507 | copper ion binding | 6 | 0.01759807 | 0.35065 | 18 |
| 282 | GO:0008283 | cell proliferation | 4 | 0.01779742 | 0.18949 | 26 |
| 283 | GO:0004450 | isocitrate dehydrogenase (NADP+) acti | 8 | 0.01789968 | 0.99105 | 1 |
| 284 | GO:0006102 | isocitrate metabolic process | 5 | 0.01789968 | 0.99105 | 1 |
| 285 | GO:0006097 | glyoxylate cycle | 6 | 0.01789968 | 0.99105 | 1 |
| 286 | GO:0031941 | filamentous actin | 6 | 0.01795602 | 0.58384 | 4 |
| 287 | GO:0050832 | defense response to fungus | 6 | 0.01808912 | 0.29892 | 19 |
| 288 | GO:0031629 | synaptic vesicle fusion to presynaptic n | 4 | 0.01834418 | -0.7907 | 3 |
| 289 | GO:0017105 | acyl-CoA delta11-desaturase activity | 6 | 0.01847708 | 0.99076 | 1 |
| 290 | GO:0004972 | N-methyl-D-aspartate selective glutam | 6 | 0.01872804 | 0.63141 | 5 |
| 291 | GO:0035561 | #N/A | 7 | 0.01886134 | 0.78877 | 3 |
| 292 | GO:0045839 | negative regulation of mitosis | 7 | 0.01886134 | 0.78877 | 3 |
| 293 | GO:0045892 | negative regulation of transcription, DN | 7 | 0.01902958 | -0.1866 | 33 |
| 294 | GO:0016740 | transferase activity | 8 | 0.01940694 | 0.4051 | 7 |
| 295 | GO:0007058 | spindle assembly involved in female m | 7 | 0.01947306 | -0.6907 | 4 |
| 296 | GO:0006446 | regulation of translational initiation | 9 | 0.01947306 | -0.6907 | 4 |
| 297 | GO:0030163 | protein catabolic process | 7 | 0.01947492 | 0.32985 | 16 |
| 298 | GO:0006163 | purine nucleotide metabolic process | 6 | 0.01948511 | 0.90132 | 2 |
| 299 | GO:0008476 | protein-tyrosine sulfotransferase activi | 6 | 0.01948755 | 0.99026 | 1 |
| 300 | GO:0031936 | negative regulation of chromatin silenc | 6 | 0.01961481 | -0.5787 | 1 |
| 301 | GO:0003844 | 1,4-alpha-glucan branching enzyme ac | 7 | 0.01977625 | 0.99011 | 1 |
| 302 | GO:0045746 | negative regulation of Notch signaling | 9 | 0.01983612 | -0.3077 | 3 |
| 303 | GO:0008295 | spermidine biosynthetic process | 5 | 0.02004519 | 0.78444 | 3 |
| 304 | GO:0008649 | rRNA methyltransferase activity | 6 | 0.02008815 | 0.89981 | 2 |
| 305 | GO:0006911 | phagocytosis, engulfment | 6 | 0.02039442 | -0.1128 | 77 |
| 306 | GO:0006974 | response to DNA damage stimulus | 6 | 0.02067481 | 0.15855 | 27 |
| 307 | GO:0009328 | phenylalanine-tRNA ligase complex | 6 | 0.02072977 | 0.89822 | 2 |
| 308 | GO:0006432 | phenylalanyl-tRNA aminoacylation | 5 | 0.02072977 | 0.89822 | 2 |
| 309 | GO:0004826 | phenylalanine-tRNA ligase activity | 6 | 0.02072977 | 0.89822 | 2 |
| 310 | GO:0016614 | oxidoreductase activity, acting on CH-C | 8 | 0.02086722 | 0.35381 | 16 |
| 311 | GO:0005096 | GTPase activator activity | 7 | 0.02094127 | -0.3272 | 9 |
| 312 | GO:0007216 | metabotropic glutamate receptor signa | 6 | 0.02119387 | 0.78039 | 3 |
| 313 | GO:0005353 | fructose transmembrane transporter a | 5 | 0.02119387 | 0.78039 | 3 |
| 314 | GO:0017146 | N-methyl-D-aspartate selective glutam | 9 | 0.02119387 | 0.78039 | 3 |
| 315 | GO:0005794 | Golgi apparatus | 10 | 0.02120973 | -0.1765 | 41 |
| 316 | GO:0005099 | Ras GTPase activator activity | 4 | 0.02128558 | -0.6232 | 1 |
| 317 | GO:0000123 | histone acetyltransferase complex | 7 | 0.02146031 | -0.3527 | 3 |
| 318 | GO:0051726 | regulation of cell cycle | 6 | 0.02149009 | -0.236 | 16 |
| 319 | GO:0008414 | CDP-alcohol phosphotransferase activit | 6 | 0.02150848 | 0.98924 | 1 |
| 320 | GO:0005665 | DNA-directed RNA polymerase II, core | 8 | 0.02180646 | 0.40003 | 8 |
| 321 | GO:0045893 | positive regulation of transcription, DN | 7 | 0.0221194 | -0.1821 | 25 |
| 322 | GO:0051124 | synaptic growth at neuromuscular junc | 7 | 0.02216163 | -0.3618 | 4 |
| 323 | GO:0005249 | voltage-gated potassium channel activ | 6 | 0.02220426 | -0.3332 | 7 |
| 324 | GO:0005391 | sodium:potassium-exchanging ATPase | 6 | 0.02233681 | -0.5323 | 1 |
| 325 | GO:0030716 | oocyte fate determination | 5 | 0.02236396 | -0.7764 | 3 |
| 326 | GO:0000242 | pericentriolar material | 6 | 0.02260129 | -0.5702 | 6 |
| 327 | GO:0048488 | synaptic vesicle endocytosis | 5 | 0.02267695 | -0.2706 | 9 |
| 328 | GO:0042654 | ecdysis-triggering hormone receptor ac | 6 | 0.02280765 | 0.9886 | 1 |
| 329 | GO:0010043 | response to zinc ion | 3 | 0.02288775 | 0.77469 | 3 |
| 330 | GO:0004730 | pseudouridylate synthase activity | 4 | 0.02288775 | 0.77469 | 3 |
| 331 | GO:0008098 | 5S rRNA primary transcript binding | 4 | 0.022952 | 0.98852 | 1 |
| 332 | GO:0008444 | CDP-diacylglycerol-glycerol-3-phosphat | 5 | 0.02309636 | 0.98845 | 1 |
| 333 | GO:0007160 | cell-matrix adhesion | 6 | 0.02330877 | 0.3971 | 6 |
| 334 | GO:0005774 | vacuolar membrane | 5 | 0.02338506 | 0.98831 | 1 |
| 335 | GO:0046686 | response to cadmium ion | 5 | 0.02338506 | 0.98831 | 1 |
| 336 | GO:0070574 | #N/A | 6 | 0.02338506 | 0.98831 | 1 |
| 337 | GO:0015086 | cadmium ion transmembrane transpor | 5 | 0.02338506 | 0.98831 | 1 |
| 338 | GO:0005769 | early endosome | 5 | 0.02345899 | -0.3493 | 4 |
| 339 | GO:0043401 | steroid hormone mediated signaling | 4 | 0.0234623 | -0.3231 | 6 |
| 340 | GO:0006098 | pentose-phosphate shunt | 5 | 0.02348645 | 0.56791 | 6 |
| 341 | GO:0043021 | ribonucleoprotein binding | 5 | 0.02352941 | 0.98823 | 1 |
| 342 | GO:0019207 | kinase regulator activity | 5 | 0.02367376 | 0.98816 | 1 |
| 343 | GO:0043154 | negative regulation of caspase activity | 5 | 0.02383631 | -0.8909 | 2 |
| 344 | GO:0005978 | glycogen biosynthetic process | 5 | 0.02392162 | 0.67694 | 3 |
| 345 | GO:0005918 | septate junction | 5 | 0.02429741 | -0.3132 | 9 |
| 346 | GO:0031473 | myosin III binding | 5 | 0.02439553 | 0.9878 | 1 |
| 347 | GO:0007155 | cell adhesion | 6 | 0.0244562 | 0.1372 | 66 |
| 348 | GO:0042719 | mitochondrial intermembrane space p | 5 | 0.0245161 | 0.49565 | 6 |
| 349 | GO:0006784 | heme a biosynthetic process | 5 | 0.02453988 | 0.98773 | 1 |
| 350 | GO:0004004 | ATP-dependent RNA helicase activity | 6 | 0.02461276 | 0.23507 | 26 |
| 351 | GO:0070971 | #N/A | 4 | 0.02482341 | 0.88862 | 2 |
| 352 | GO:0006686 | sphingomyelin biosynthetic process | 4 | 0.02497293 | 0.98751 | 1 |
| 353 | GO:0016272 | prefoldin complex | 5 | 0.02503196 | 0.44568 | 7 |
| 354 | GO:0008092 | cytoskeletal protein binding | 4 | 0.02508352 | -0.305 | 10 |
| 355 | GO:0007304 | chorion-containing eggshell formation | 4 | 0.02511127 | 0.36792 | 12 |
| 356 | GO:0045742 | positive regulation of epidermal growt | 7 | 0.02529822 | -0.5634 | 1 |
| 357 | GO:0008568 | microtubule-severing ATPase activity | 8 | 0.0252991 | 0.76704 | 3 |
| 358 | GO:0008518 | reduced folate carrier activity | 5 | 0.0252991 | 0.76704 | 3 |
| 359 | GO:0007099 | centriole replication | 4 | 0.02551966 | -0.328 | 6 |
| 360 | GO:0000070 | mitotic sister chromatid segregation | 4 | 0.02554347 | 0.31114 | 13 |
| 361 | GO:0045544 | gibberellin 20-oxidase activity | 4 | 0.02555034 | 0.98722 | 1 |
| 362 | GO:0004252 | serine-type endopeptidase activity | 4 | 0.02572837 | 0.08869 | 279 |
| 363 | GO:0033227 | dsRNA transport | 4 | 0.02575885 | -0.304 | 6 |
| 364 | GO:0006855 | multidrug transport | 5 | 0.02619273 | 0.88558 | 2 |
| 365 | GO:0016201 | synaptic target inhibition | 4 | 0.02622578 | -0.8855 | 2 |
| 366 | GO:0004852 | uroporphyrinogen-III synthase activity | 4 | 0.02642452 | 0.88508 | 2 |
| 367 | GO:0008630 | DNA damage response, signal transduc | 5 | 0.02654732 | 0.46509 | 6 |
| 368 | GO:0009298 | GDP-mannose biosynthetic process | 5 | 0.02656081 | 0.98672 | 1 |
| 369 | GO:0019307 | mannose biosynthetic process | 4 | 0.02656081 | 0.98672 | 1 |
| 370 | GO:0006662 | glycerol ether metabolic process | 4 | 0.02711446 | 0.36469 | 14 |
| 371 | GO:0015276 | ligand-gated ion channel activity | 4 | 0.02720918 | -0.1924 | 18 |
| 372 | GO:0017176 | phosphatidylinositol N-acetylglucosam | 4 | 0.02734188 | 0.66782 | 4 |
| 373 | GO:0000220 | vacuolar proton-transporting V-type AT | 4 | 0.02770806 | -0.3638 | 5 |
| 374 | GO:0070822 | #N/A | 7 | 0.02784393 | -0.4211 | 3 |
| 375 | GO:0030178 | negative regulation of Wnt receptor sig | 6 | 0.02794096 | 0.31671 | 9 |
| 376 | GO:0006030 | chitin metabolic process | 7 | 0.02806445 | 0.16113 | 48 |
| 377 | GO:0010629 | negative regulation of gene expression | 7 | 0.02829312 | 0.20038 | 47 |
| 378 | GO:0019867 | outer membrane | 7 | 0.02843739 | 0.98578 | 1 |
| 379 | GO:0043652 | engulfment of apoptotic cell | 7 | 0.0284532 | 0.88075 | 2 |
| 380 | GO:0008061 | chitin binding | 7 | 0.02846935 | 0.1544 | 76 |
| 381 | GO:0000010 | trans-hexaprenyltranstransferase activ | 8 | 0.0289721 | 0.87967 | 2 |
| 382 | GO:0010507 | negative regulation of autophagy | 8 | 0.02898931 | 0.66378 | 4 |
| 383 | GO:0001578 | microtubule bundle formation | 7 | 0.02903164 | 0.51698 | 4 |
| 384 | GO:0007274 | neuromuscular synaptic transmission | 8 | 0.02903575 | -0.2667 | 11 |
| 385 | GO:0004854 | xanthine dehydrogenase activity | 7 | 0.02904164 | 0.87952 | 2 |
| 386 | GO:0070855 | #N/A | 8 | 0.02907644 | 0.87945 | 2 |
| 387 | GO:0045495 | pole plasm | 6 | 0.02941645 | 0.4184 | 6 |
| 388 | GO:0045169 | fusome | 6 | 0.02949176 | -0.2359 | 18 |
| 389 | GO:0035076 | ecdysone receptor-mediated signaling | 6 | 0.02974941 | -0.4849 | 1 |
| 390 | GO:0035242 | protein-arginine omega-N asymmetric | 6 | 0.02977684 | 0.878 | 2 |
| 391 | GO:0035241 | protein-arginine omega-N monomethy | 6 | 0.02977684 | 0.878 | 2 |
| 392 | GO:0006680 | glucosylceramide catabolic process | 7 | 0.03016961 | 0.98491 | 1 |
| 393 | GO:0016142 | O-glycoside catabolic process | 7 | 0.03016961 | 0.98491 | 1 |
| 394 | GO:0008422 | beta-glucosidase activity | 6 | 0.03016961 | 0.98491 | 1 |
| 395 | GO:0008206 | bile acid metabolic process | 8 | 0.03016961 | 0.98491 | 1 |
| 396 | GO:0003951 | NAD+ kinase activity | 5 | 0.0302459 | -0.7527 | 3 |
| 397 | GO:0045786 | negative regulation of cell cycle | 4 | 0.0302459 | -0.7527 | 3 |
| 398 | GO:0030688 | preribosome, small subunit precursor | 4 | 0.03031397 | 0.98484 | 1 |
| 399 | GO:0015367 | oxoglutarate:malate antiporter activity | 4 | 0.03052142 | -0.6602 | 1 |
| 400 | GO:0070389 | chaperone cofactor-dependent protein | 4 | 0.03055691 | 0.87642 | 2 |
| 401 | GO:0090254 | #N/A | 5 | 0.03057727 | -0.5989 | 5 |
| 402 | GO:0007029 | endoplasmic reticulum organization | 5 | 0.03081247 | 0.48288 | 5 |
| 403 | GO:0008514 | organic anion transmembrane transpo | 5 | 0.03095378 | 0.45692 | 8 |
| 404 | GO:0045202 | synapse | 4 | 0.0311148 | -0.3103 | 10 |
| 405 | GO:0006888 | ER to Golgi vesicle-mediated transport | 6 | 0.03143838 | 0.35838 | 9 |
| 406 | GO:0016709 | oxidoreductase activity, acting on paired | 6 | 0.03161314 | 0.98419 | 1 |
| 407 | GO:0000103 | sulfate assimilation | 6 | 0.03204619 | 0.98398 | 1 |
| 408 | GO:0004020 | adenylylsulfate kinase activity | 7 | 0.03204619 | 0.98398 | 1 |
| 409 | GO:0004781 | sulfate adenylyltransferase (ATP) activ | 5 | 0.03204619 | 0.98398 | 1 |
| 410 | GO:0008137 | NADH dehydrogenase (ubiquinone) act | 8 | 0.03210482 | -0.2365 | 13 |
| 411 | GO:0046328 | regulation of JNK cascade | 4 | 0.03233705 | -0.4801 | 2 |
| 412 | GO:0030337 | DNA polymerase processivity factor ac | 7 | 0.03240404 | 0.87274 | 2 |
| 413 | GO:0006272 | leading strand elongation | 6 | 0.03240404 | 0.87274 | 2 |
| 414 | GO:0043626 | PCNA complex | 6 | 0.03240404 | 0.87274 | 2 |
| 415 | GO:0007394 | dorsal closure, elongation of leading ec | 7 | 0.03254564 | -0.4134 | 2 |
| 416 | GO:0005687 | snRNP U4 | 8 | 0.03277258 | 0.87201 | 2 |
| 417 | GO:0006123 | mitochondrial electron transport, cyto | 6 | 0.03297952 | -0.3682 | 4 |
| 418 | GO:0035191 | nuclear axial expansion | 6 | 0.0330848 | -0.7452 | 3 |
| 419 | GO:0005371 | tricarboxylate secondary active transm | 4 | 0.0330848 | -0.7452 | 3 |
| 420 | GO:0051233 | spindle midzone | 5 | 0.033444 | 0.41204 | 6 |
| 421 | GO:0046959 | habituation | 4 | 0.03344402 | -0.6537 | 4 |
| 422 | GO:0008475 | procollagen-lysine 5-dioxygenase activ | 5 | 0.03362813 | -0.8704 | 2 |
| 423 | GO:0031670 | cellular response to nutrient | 3 | 0.03363407 | 0.98318 | 1 |
| 424 | GO:0005662 | DNA replication factor A complex | 7 | 0.03370305 | 0.87021 | 2 |
| 425 | GO:0003837 | beta-ureidopropionase activity | 4 | 0.03377842 | 0.98311 | 1 |
| 426 | GO:0006591 | ornithine metabolic process | 4 | 0.03406712 | 0.98297 | 1 |
| 427 | GO:0006879 | cellular iron ion homeostasis | 8 | 0.03421757 | -0.5071 | 1 |
| 428 | GO:0005919 | pleated septate junction | 7 | 0.03426758 | -0.8691 | 2 |
| 429 | GO:0021682 | nerve maturation | 4 | 0.03426758 | -0.8691 | 2 |
| 430 | GO:0031571 | G1 DNA damage checkpoint | 5 | 0.0343432 | -0.869 | 2 |
| 431 | GO:0048477 | oogenesis | 5 | 0.03449624 | 0.10355 | 186 |
| 432 | GO:0005488 | binding | 3 | 0.03454531 | 0.09083 | 247 |
| 433 | GO:0017057 | 6-phosphogluconolactonase activity | 5 | 0.03464453 | 0.98268 | 1 |
| 434 | GO:0005483 | soluble NSF attachment protein activit | 4 | 0.03465091 | 0.65122 | 4 |
| 435 | GO:0004449 | isocitrate dehydrogenase (NAD+) activ | 7 | 0.03466755 | -0.5437 | 1 |
| 436 | GO:0016250 | N-sulfoglucosamine sulfohydrolase act | 6 | 0.03493324 | 0.98253 | 1 |
| 437 | GO:0008484 | sulfuric ester hydrolase activity | 7 | 0.03493324 | 0.98253 | 1 |
| 438 | GO:0004728 | receptor signaling protein tyrosine pho | 8 | 0.03538556 | -0.7395 | 3 |
| 439 | GO:0007156 | homophilic cell adhesion | 8 | 0.03544383 | 0.2692 | 24 |
| 440 | GO:0004091 | carboxylesterase activity | 6 | 0.03570238 | 0.23996 | 29 |
| 441 | GO:0003896 | DNA primase activity | 7 | 0.0357187 | 0.86638 | 2 |
| 442 | GO:0008608 | attachment of spindle microtubules to | 6 | 0.0357187 | 0.86638 | 2 |
| 443 | GO:0006269 | DNA replication, synthesis of RNA prim | 4 | 0.0357187 | 0.86638 | 2 |
| 444 | GO:0006144 | purine base metabolic process | 4 | 0.0357187 | 0.86638 | 2 |
| 445 | GO:0017053 | transcriptional repressor complex | 4 | 0.03589693 | -0.4085 | 2 |
| 446 | GO:0015992 | proton transport | 4 | 0.03633731 | -0.2635 | 10 |
| 447 | GO:0004115 | 3',5'-cyclic-AMP phosphodiesterase act | 4 | 0.03637764 | -0.8652 | 2 |
| 448 | GO:0009058 | biosynthetic process | 4 | 0.03640556 | 0.30672 | 18 |
| 449 | GO:0005083 | small GTPase regulator activity | 4 | 0.0364999 | -0.4479 | 2 |
| 450 | GO:0004768 | stearoyl-CoA 9-desaturase activity | 4 | 0.03654167 | 0.50314 | 5 |
| 451 | GO:0008532 | N-acetyllactosaminide beta-1,3-N-acetylglucosaminidase activity | 4 | 0.03663505 | 0.73643 | 3 |
| 452 | GO:0050777 | negative regulation of immune response | 4 | 0.03680982 | 0.98159 | 1 |
| 453 | GO:0006820 | anion transport | 5 | 0.03691626 | 0.50251 | 6 |
| 454 | GO:0004644 | phosphoribosylglycinamide formyltransferase activity | 4 | 0.03695417 | 0.98152 | 1 |
| 455 | GO:0004641 | phosphoribosylformylglycinamidine cyclase activity | 4 | 0.03695417 | 0.98152 | 1 |
| 456 | GO:0004637 | phosphoribosylamine-glycine ligase activity | 6 | 0.03695417 | 0.98152 | 1 |
| 457 | GO:0007507 | heart development | 4 | 0.03713452 | -0.1918 | 8 |
| 458 | GO:0016331 | morphogenesis of embryonic epithelium | 4 | 0.03719028 | -0.472 | 2 |
| 459 | GO:0032880 | regulation of protein localization | 5 | 0.03745862 | 0.58448 | 5 |
| 460 | GO:0003007 | heart morphogenesis | 4 | 0.03754532 | -0.7343 | 3 |
| 461 | GO:0004527 | exonuclease activity | 4 | 0.03787673 | 0.53799 | 4 |
| 462 | GO:0007096 | regulation of exit from mitosis | 5 | 0.0379037 | 0.44585 | 5 |
| 463 | GO:0002807 | positive regulation of antimicrobial peptide production | 5 | 0.03796463 | 0.98102 | 1 |
| 464 | GO:0007541 | sex determination, primary response to steroid hormone | 7 | 0.03798657 | -0.5835 | 1 |
| 465 | GO:0019730 | antimicrobial humoral response | 4 | 0.03820476 | -0.2281 | 11 |
| 466 | GO:0070983 | #N/A | 5 | 0.03828997 | -0.3295 | 4 |
| 467 | GO:0016204 | determination of muscle attachment sites | 6 | 0.03877887 | 0.64308 | 3 |
| 468 | GO:0035007 | regulation of melanization defense response | 5 | 0.03883183 | -0.8607 | 2 |
| 469 | GO:0045886 | negative regulation of synaptic growth | 5 | 0.03900605 | -0.3736 | 3 |
| 470 | GO:0010669 | epithelial structure maintenance | 8 | 0.03927562 | -0.8599 | 2 |
| 471 | GO:0003713 | transcription coactivator activity | 8 | 0.0393309 | -0.2369 | 16 |
| 472 | GO:0035363 | #N/A | 8 | 0.03955025 | -0.4684 | 2 |
| 473 | GO:0030330 | DNA damage response, signal transduction | 8 | 0.03955251 | 0.98022 | 1 |
| 474 | GO:0016491 | oxidoreductase activity | 8 | 0.03957415 | 0.15156 | 49 |
| 475 | GO:0007476 | imaginal disc-derived wing morphogenesis | 5 | 0.03976603 | -0.1351 | 45 |
| 476 | GO:0019064 | viral envelope fusion with host membrane | 6 | 0.03998557 | 0.98001 | 1 |
| 477 | GO:0019031 | viral envelope | 7 | 0.03998557 | 0.98001 | 1 |
| 478 | GO:0046789 | host cell surface receptor binding | 7 | 0.03998557 | 0.98001 | 1 |
| 479 | GO:0004307 | ethanolaminephosphotransferase activity | 7 | 0.04027427 | 0.97986 | 1 |
| 480 | GO:0048142 | germarium-derived cystoblast division | 7 | 0.04049877 | 0.85772 | 2 |
| 481 | GO:0015867 | ATP transport | 4 | 0.04049877 | 0.85772 | 2 |
| 482 | GO:0015866 | ADP transport | 6 | 0.04049877 | 0.85772 | 2 |
| 483 | GO:0005958 | DNA-dependent protein kinase-DNA ligase activity | 5 | 0.04049877 | 0.85772 | 2 |
| 484 | GO:0045900 | negative regulation of translational elongation | 6 | 0.04049877 | 0.85772 | 2 |
| 485 | GO:0005471 | ATP:ADP antiporter activity | 4 | 0.04049877 | 0.85772 | 2 |
| 486 | GO:0008079 | translation termination factor activity | 5 | 0.04049877 | 0.85772 | 2 |
| 487 | GO:0006892 | post-Golgi vesicle-mediated transport | 3 | 0.04050757 | 0.63988 | 4 |
| 488 | GO:0031000 | response to caffeine | 5 | 0.04050757 | 0.63988 | 4 |
| 489 | GO:0005873 | plus-end kinesin complex | 5 | 0.04052709 | 0.7274 | 3 |
| 490 | GO:0044432 | endoplasmic reticulum part | 5 | 0.04053986 | 0.85765 | 2 |
| 491 | GO:0032147 | activation of protein kinase activity | 4 | 0.04056297 | 0.97972 | 1 |
| 492 | GO:0006886 | intracellular protein transport | 5 | 0.04105921 | 0.16616 | 26 |
| 493 | GO:0048489 | synaptic vesicle transport | 7 | 0.04106245 | -0.4016 | 5 |
| 494 | GO:0017133 | mitochondrial electron transfer flavopr | 5 | 0.04115878 | -0.8566 | 2 |
| 495 | GO:0007548 | sex differentiation | 6 | 0.04130445 | -0.3263 | 4 |
| 496 | GO:0051298 | centrosome duplication | 4 | 0.0416269 | -0.1748 | 16 |
| 497 | GO:0004591 | oxoglutarate dehydrogenase (succinyl- | 5 | 0.0417955 | 0.72459 | 3 |
| 498 | GO:0051533 | positive regulation of NFAT protein im | 9 | 0.0419291 | 0.34574 | 9 |
| 499 | GO:0046530 | photoreceptor cell differentiation | 8 | 0.04194948 | -0.8552 | 2 |
| 500 | GO:0000902 | cell morphogenesis | 5 | 0.04253996 | 0.21373 | 34 |
| 501 | GO:0046012 | positive regulation of oskar mRNA tran | 6 | 0.04302308 | -0.7219 | 3 |
| 502 | GO:0046855 | inositol phosphate dephosphorylation | 6 | 0.04302308 | -0.7219 | 3 |
| 503 | GO:0009617 | response to bacterium | 5 | 0.04310033 | -0.3002 | 7 |
| 504 | GO:0033014 | tetrapyrrole biosynthetic process | 6 | 0.04317914 | 0.63513 | 3 |
| 505 | GO:0005252 | open rectifier potassium channel activi | 6 | 0.04340274 | 0.63476 | 4 |
| 506 | GO:0015232 | heme transporter activity | 6 | 0.04363994 | 0.63434 | 3 |
| 507 | GO:0003779 | actin binding | 5 | 0.04372225 | -0.131 | 31 |
| 508 | GO:0009267 | cellular response to starvation | 6 | 0.04397748 | -0.6338 | 1 |
| 509 | GO:0097062 | #N/A | 6 | 0.04402743 | 0.97798 | 1 |
| 510 | GO:0000309 | nicotinamide-nucleotide adenylyltransf | 6 | 0.04402743 | 0.97798 | 1 |
| 511 | GO:0005786 | signal recognition particle, endoplasmic | 4 | 0.04419799 | 0.43721 | 6 |
| 512 | GO:0050918 | positive chemotaxis | 5 | 0.04446048 | 0.97777 | 1 |
| 513 | GO:0004407 | histone deacetylase activity | 5 | 0.0446733 | -0.527 | 1 |
| 514 | GO:0048027 | mRNA 5'-UTR binding | 5 | 0.04471139 | -0.8505 | 2 |
| 515 | GO:0031589 | cell-substrate adhesion | 3 | 0.04471982 | -0.7183 | 3 |
| 516 | GO:0042250 | maintenance of polarity of embryonic c | 4 | 0.04489354 | 0.97755 | 1 |
| 517 | GO:0016585 | chromatin remodeling complex | 4 | 0.04505797 | -0.4901 | 2 |
| 518 | GO:0051297 | centrosome organization | 4 | 0.04527887 | -0.207 | 9 |
| 519 | GO:0034773 | histone H4-K20 trimethylation | 4 | 0.04536125 | -0.8494 | 2 |
| 520 | GO:0005525 | GTP binding | 4 | 0.04578626 | 0.10864 | 86 |
| 521 | GO:0014706 | striated muscle development | 4 | 0.04604836 | 0.97697 | 1 |
| 522 | GO:0005484 | SNAP receptor activity | 8 | 0.04619049 | 0.30485 | 14 |
| 523 | GO:0000302 | response to reactive oxygen species | 4 | 0.04656647 | -0.7145 | 3 |
| 524 | GO:0019985 | bypass DNA synthesis | 8 | 0.04676754 | 0.62912 | 4 |
| 525 | GO:0005791 | rough endoplasmic reticulum | 4 | 0.04693292 | 0.56801 | 5 |
| 526 | GO:0060388 | vitelline envelope | 4 | 0.04693292 | 0.56801 | 5 |
| 527 | GO:0030133 | transport vesicle | 5 | 0.04702485 | 0.56787 | 5 |
| 528 | GO:0060361 | flight | 6 | 0.04702485 | 0.56787 | 5 |
| 529 | GO:0005850 | eukaryotic translation initiation factor | 6 | 0.04707087 | 0.5678 | 5 |
| 530 | GO:0019955 | cytokine binding | 5 | 0.04738341 | -0.8461 | 2 |
| 531 | GO:0005813 | centrosome | 4 | 0.04754791 | -0.1821 | 28 |
| 532 | GO:0043565 | sequence-specific DNA binding | 5 | 0.04767608 | 0.09077 | 227 |
| 533 | GO:0017064 | fatty acid amide hydrolase activity | 6 | 0.04775218 | -0.6275 | 4 |
| 534 | GO:0016884 | carbon-nitrogen ligase activity, with gl | 6 | 0.04775218 | -0.6275 | 4 |
| 535 | GO:0000178 | exosome (RNase complex) | 4 | 0.04787351 | 0.8453 | 2 |
| 536 | GO:0030722 | establishment of oocyte nucleus localiz | 7 | 0.04787351 | 0.8453 | 2 |
| 537 | GO:0042138 | meiotic DNA double-strand break form | 7 | 0.04787351 | 0.8453 | 2 |
| 538 | GO:0004766 | spermidine synthase activity | 6 | 0.04787351 | 0.8453 | 2 |
| 539 | GO:0043462 | regulation of ATPase activity | 5 | 0.04787351 | 0.8453 | 2 |
| 540 | GO:0032934 | sterol binding | 6 | 0.04788491 | 0.37777 | 11 |
| 541 | GO:0021551 | central nervous system morphogenesis | 6 | 0.04864481 | -0.7103 | 3 |
| 542 | GO:0006325 | establishment or maintenance of chrom | 6 | 0.04888273 | -0.253 | 10 |
| 543 | GO:0046536 | dosage compensation complex | 7 | 0.04899673 | -0.8435 | 2 |
| 544 | GO:0005815 | microtubule organizing center | 4 | 0.04905473 | -0.4847 | 1 |
| 545 | GO:0005214 | structural constituent of chitin-based c | 5 | 0.04959323 | 0.11774 | 113 |

**Table S3:**
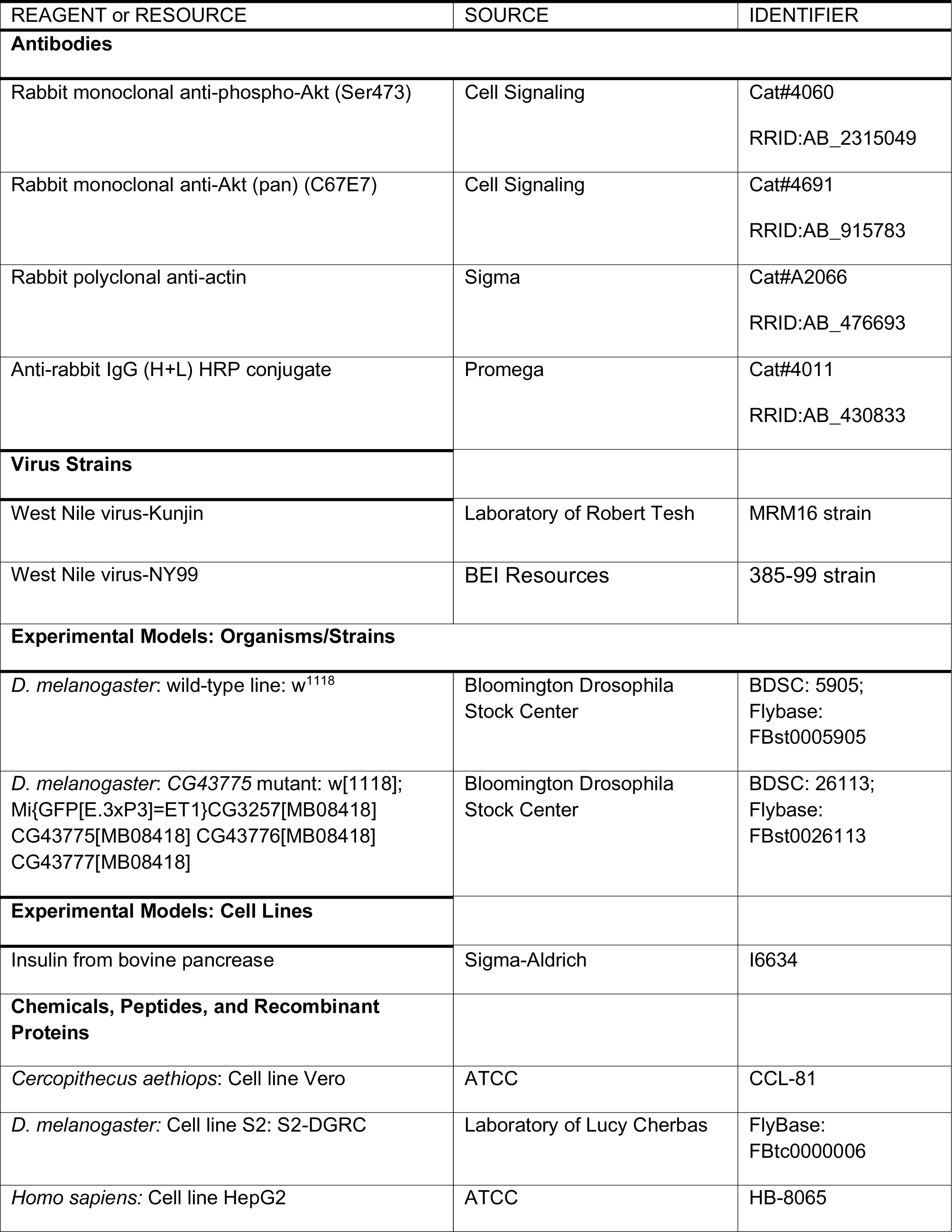
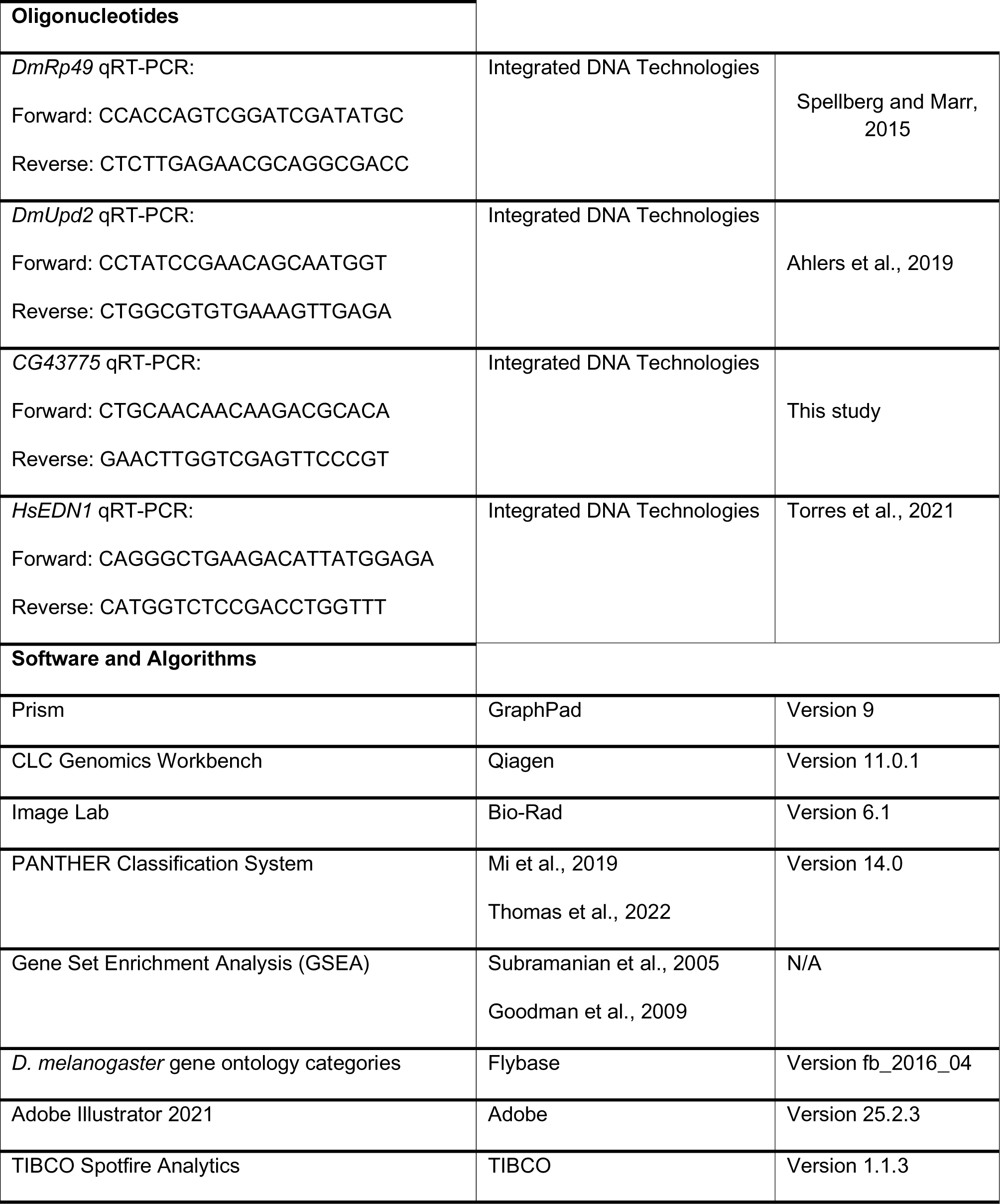
Fly lines and reagents used in this study.

**Figure S1:**
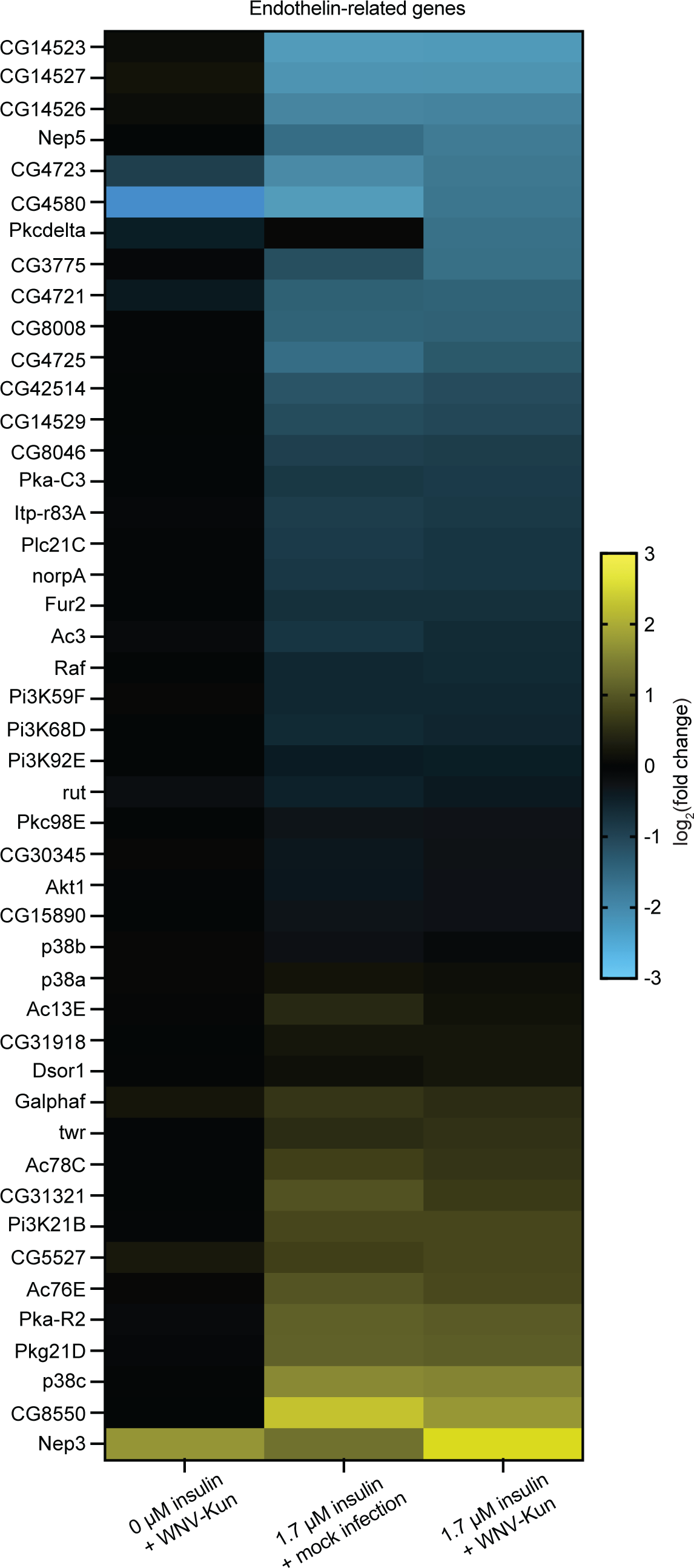
Heat map expression of genes transcriptionally enriched or suppressed as identified in Fig. 1E. PANTHER-GO analysis identifies this gene set associated with the endothelin signaling pathway.

**Figure S2:**
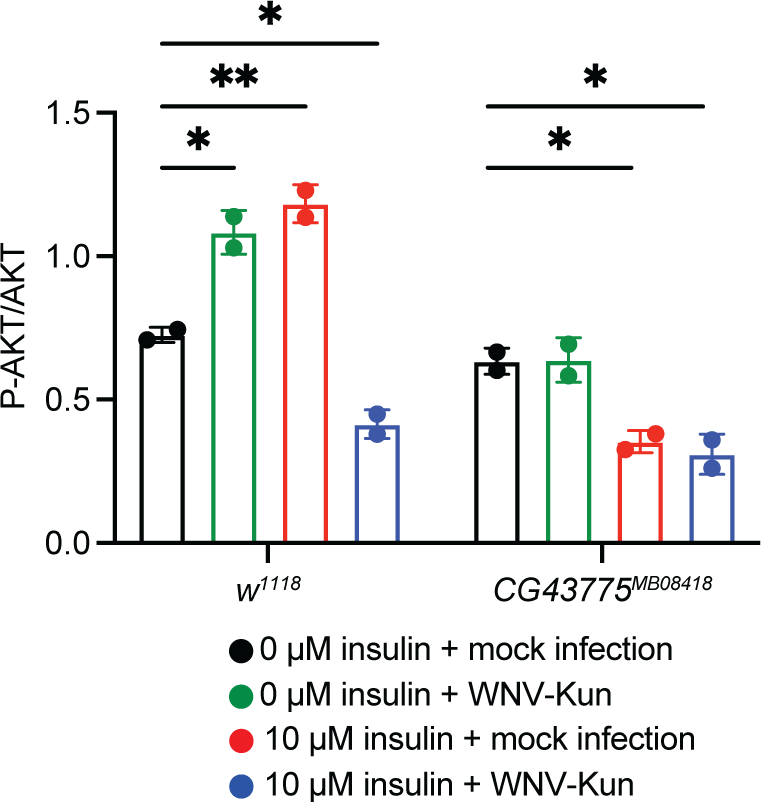
AKT phosphorylation is diminished in insulin-treated *CG43775* mutant flies but not control flies as analyzed in Fig. 2E. Densitometry analysis of western blots measuring P-AKT abundance relative to AKT shows reduced activation in *CG43775* mutants compared to control flies. (*p < 0.05, **p < 0.01, One-way ANOVA). Circles represent individual experimental replications. Horizontal bars represent the mean. Error bars represent SDs. Results are of pooled duplicate independent experiments.

**Figure S3:**
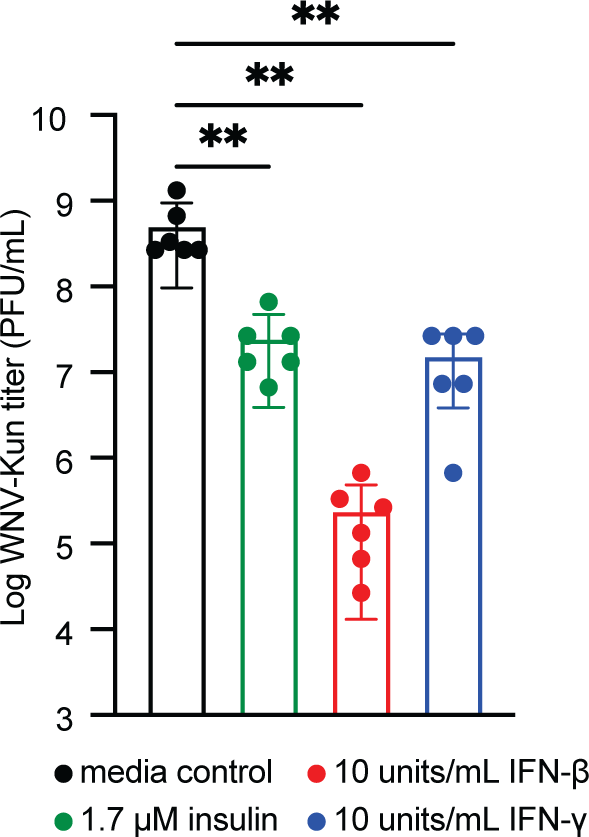
Insulin reduces WNV-Kun titer in HepG2 cells to similar levels as IFN-β or IFN-γ treatment. WNV-Kun titer at 2 d p.i is reduced in cells that received either 1.7 μM insulin, 10 units/mL IFN-β, or 10 units/mL IFN-γ treatment 24 h prior to infection (MOI=0.01 PFU/cell) (**p < 0.01, One-way ANOVA). Circles represent individual biological replications. Horizontal bars represent the mean. Error bars represent SDs. Results are representative of duplicate independent experiments.

**Figure S4:**
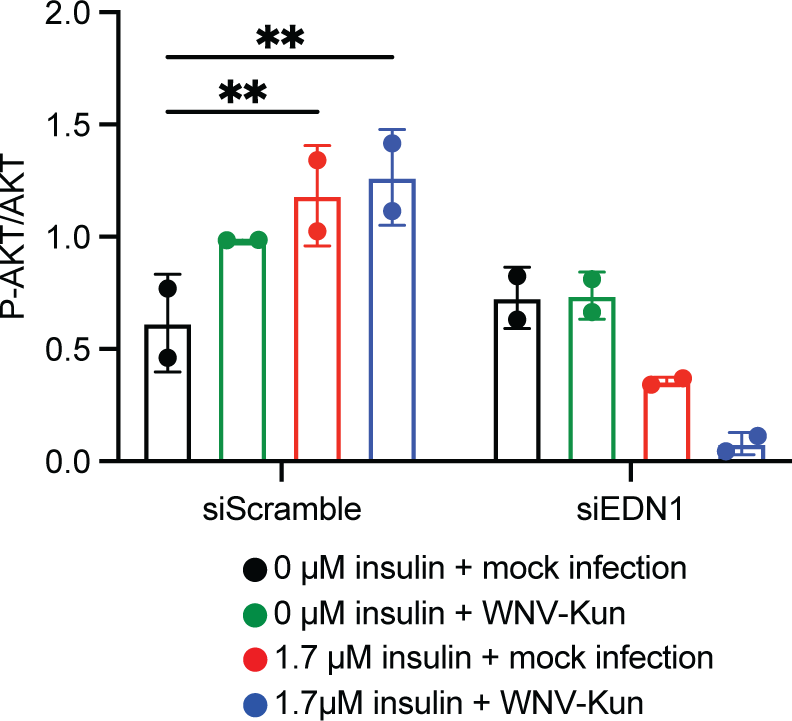
AKT phosphorylation is enhanced in HepG2 cells following insulin treatment and WNV-Kun infection but diminished following siEDN1 transfection as analyzed in Fig. 3E. Densitometry analysis of western blots measuring P-AKT abundance relative to AKT shows reduced activation in siEDN1 transfected HepG2 cells compared to siScramble transfected cells. (**p < 0.01, One-way ANOVA). Circles represent individual experimental replications. Horizontal bars represent the mean. Error bars represent SDs. Results are of pooled duplicate independent experiments.

## REFERENCES

1. Ahlers LRH, Goodman AG. 2018. The Immune Responses of the Animal Hosts of West Nile Virus: A Comparison of Insects, Birds, and Mammals. Front Cell Infect Microbiol 8:96.

2. Centers for Disease Control and Prevention (CDC). 1999. Outbreak of West Nile-like viral encephalitis--New York, 1999. MMWR Morb Mortal Wkly Rep 48:845–849.

3. Nash D, Mostashari F, Fine A, Miller J, O’Leary D, Murray K, Huang A, Rosenberg A, Greenberg A, Sherman M, Wong S, Campbell GL, Roehrig JT, Gubler DJ, Shieh W-J, Zaki S, Smith P, Layton M. 2001. The Outbreak of West Nile Virus Infection in the New York City Area in 1999. N Engl J Med 344:1807–1814.

4. Lanciotti RS, Roehrig JT, Deubel V, Smith J, Parker M, Steele K, Crise B, Volpe KE, Crabtree MB, Scherret JH, Hall RA, MacKenzie JS, Cropp CB, Panigrahy B, Ostlund E, Schmitt B, Malkinson M, Banet C, Weissman J, Komar N, Savage HM, Stone W, McNamara T, Gubler DJ. 1999. Origin of the West Nile virus responsible for an outbreak of encephalitis in the northeastern United States. Science 286:2333–2337.

5. Gorris ME, Bartlow AW, Temple SD, Romero-Alvarez D, Shutt DP, Fair JM, Kaufeld KA, Del Valle SY, Manore CA. 2021. Updated distribution maps of predominant Culex mosquitoes across the Americas. Parasit Vectors 14:547.

6. Harrigan RJ, Thomassen HA, Buermann W, Smith TB. 2014. A continental risk assessment of West Nile virus under climate change. Glob Change Biol 20:2417– 2425.

7. Ludwig A, Zheng H, Vrbova L, Drebot M, Iranpour M, Lindsay L. 2019. Increased risk of endemic mosquito-borne diseases in Canada due to climate change. Can Commun Dis Rep 45:91–97.

8. Evans BR, Kotsakiozi P, Costa-da-Silva AL, Ioshino RS, Garziera L, Pedrosa MC, Malavasi A, Virginio JF, Capurro ML, Powell JR. 2019. Transgenic Aedes aegypti Mosquitoes Transfer Genes into a Natural Population. Sci Rep 9:13047.

9. Hedges LM, Brownlie JC, O’Neill SL, Johnson KN. 2008. Wolbachia and Virus Protection in Insects. Science 322:702–702.

10. Trammell CE, Ramirez G, Sanchez-Vargas I, Clair LAS, Ratnayake OC, Luckhart S, Perera R, Goodman AG. 2022. Coupled small molecules target RNA interference and JAK/STAT signaling to reduce Zika virus infection in Aedes aegypti. PLOS Pathog 18:e1010411.

11. Alli A, Ortiz JF, Atoot A, Atoot A, Millhouse PW. 2021. Management of West Nile Encephalitis: An Uncommon Complication of West Nile Virus. Cureus 13:e13183.

12. Yasunaga A, Hanna SL, Li J, Cho H, Rose PP, Spiridigliozzi A, Gold B, Diamond MS, Cherry S. 2014. Genome-Wide RNAi Screen Identifies Broadly-Acting Host Factors That Inhibit Arbovirus Infection. PLOS Pathog 10:e1003914.

13. Ahlers LRH, Trammell CE, Carrell GF, Mackinnon S, Torrevillas BK, Chow CY, Luckhart S, Goodman AG. 2019. Insulin Potentiates JAK/STAT Signaling to Broadly Inhibit Flavivirus Replication in Insect Vectors. Cell Rep 29:1946–1960.e5.

14. Liu Y, Gordesky-Gold B, Leney-Greene M, Weinbren NL, Tudor M, Cherry S. 2018. Inflammation-Induced, STING-Dependent Autophagy Restricts Zika Virus Infection in the Drosophila Brain. Cell Host Microbe 24:57–68.e3.

15. Puig O, Marr MT, Ruhf ML, Tjian R. 2003. Control of cell number by Drosophila FOXO: downstream and feedback regulation of the insulin receptor pathway. Genes Dev 17:2006–2020.

16. Barbieri M, Bonafè M, Franceschi C, Paolisso G. 2003. Insulin/IGF-I-signaling pathway: an evolutionarily conserved mechanism of longevity from yeast to humans. Am J Physiol-Endocrinol Metab 285:E1064–E1071.

17. Šestan M, Marinović S, Kavazović I, Cekinović Đ, Wueest S, Turk Wensveen T, Brizić I, Jonjić S, Konrad D, Wensveen FM, Polić B. 2018. Virus-Induced Interferon-γ Causes Insulin Resistance in Skeletal Muscle and Derails Glycemic Control in Obesity. Immunity 49:164–177.e6.

18. Campo JA del, García-Valdecasas M, Rojas L, Rojas Á, Romero-Gómez M. 2012 The Hepatitis C Virus Modulates Insulin Signaling Pathway In Vitro Promoting Insulin Resistance. PLOS ONE 7:e47904.

19. Yu B, Li C, Sun Y, Wang DW. 2021. Insulin Treatment Is Associated with Increased Mortality in Patients with COVID-19 and Type 2 Diabetes. Cell Metab 33:65–77.e2.

20. Liang Q, Luo Z, Zeng J, Chen W, Foo S-S, Lee S-A, Ge J, Wang S, Goldman SA, Zlokovic BV, Zhao Z, Jung JU. 2016. Zika Virus NS4A and NS4B Proteins Deregulate Akt-mTOR Signaling in Human Fetal Neural Stem Cells to Inhibit Neurogenesis and Induce Autophagy. Cell Stem Cell 19:663–671.

21. Chan JF-W, Zhu Z, Chu H, Yuan S, Chik KK-H, Chan CC-S, Poon VK-M, Yip CC- Y, Zhang X, Tsang JO-L, Zou Z, Tee K-M, Shuai H, Lu G, Yuen K-Y. 2018. The celecoxib derivative kinase inhibitor AR-12 (OSU-03012) inhibits Zika virus via down-regulation of the PI3K/Akt pathway and protects Zika virus-infected A129 mice: A host-targeting treatment strategy. Antiviral Res 160:38–47.

22. Davenport AP, Hyndman KA, Dhaun N, Southan C, Kohan DE, Pollock JS, Pollock DM, Webb DJ, Maguire JJ. 2016. Endothelin. Pharmacol Rev 68:357–418.

23. Correa AF, Bailão AM, Bastos IMD, Orme IM, Soares CMA, Kipnis A, Santana JM, Junqueira-Kipnis AP. 2014. The Endothelin System Has a Significant Role in the Pathogenesis and Progression of Mycobacterium tuberculosis Infection. Infect Immun 82:5154–5165.

24. Notas G, Xidakis C, Valatas V, Kouroumalis A, Kouroumalis E. 2001. Levels of circulating endothelin-1 and nitrates/nitrites in patients with virus-related hepatocellular carcinoma. J Viral Hepat 8:63–69.

25. Freeman BD, Machado FS, Tanowitz HB, Desruisseaux MS. 2014. Endothelin-1 and its role in the pathogenesis of infectious diseases. Life Sci 118:110–119.

26. Elisa T, Antonio P, Giuseppe P, Alessandro B, Giuseppe A, Federico C, Marzia D, Ruggero B, Giacomo M, Andrea O, Daniela R, Mariaelisa R, Claudio L. 2015. Endothelin Receptors Expressed by Immune Cells Are Involved in Modulation of Inflammation and in Fibrosis: Relevance to the Pathogenesis of Systemic Sclerosis. J Immunol Res 2015:147616.

27. George PM, Cunningham ME, Galloway-Phillipps N, Badiger R, Alazawi W, Foster GR, Mitchell JA. 2012. Endothelin-1 as a mediator and potential biomarker for interferon induced pulmonary toxicity. Pulm Circ 2:501–504.

28. Jiang ZY, Zhou QL, Chatterjee A, Feener EP, Myers MG, White MF, King GL. 1999. Endothelin-1 modulates insulin signaling through phosphatidylinositol 3-kinase pathway in vascular smooth muscle cells. Diabetes 48:1120–1130.

29. Li Q, Park K, Li C, Rask-Madsen C, Mima A, Qi W, Mizutani K, Huang P, King GL. 2013. Induction of Vascular Insulin Resistance and Endothelin-1 Expression and Acceleration of Atherosclerosis by the Overexpression of Protein Kinase C-β Isoform in the Endothelium. Circ Res 113:418–427.

30. Mi H, Muruganujan A, Ebert D, Huang X, Thomas PD. 2019. PANTHER version 14: more genomes, a new PANTHER GO-slim and improvements in enrichment analysis tools. Nucleic Acids Res 47:D419–D426.

31. Thomas PD, Ebert D, Muruganujan A, Mushayahama T, Albou L-P, Mi H. 2022. PANTHER: Making genome-scale phylogenetics accessible to all. Protein Sci 31:8– 22.

32. Dagamajalu S, Rex DAB, Gopalakrishnan L, Karthikkeyan G, Gurtoo S, Modi PK, Mohanty V, Mujeeburahiman M, Soman S, Raju R, Tiwari V, Prasad TSK. 2021. A network map of endothelin mediated signaling pathway. J Cell Commun Signal 15:277–282.

33. Chahdi A, Sorokin A. 2008. Endothelin-1 Couples βPix to p66Shc: Role of βPix in Cell Proliferation through FOXO3a Phosphorylation and p27kip1 Down-Regulation Independently of Akt. Mol Biol Cell 19:2609–2619.

34. Cifarelli V, Lee S, Kim DH, Zhang T, Kamagate A, Slusher S, Bertera S, Luppi P, Trucco M, Dong HH. 2012. FOXO1 Mediates the Autocrine Effect of Endothelin-1 on Endothelial Cell Survival. Mol Endocrinol 26:1213–1224.

35. Lu J-W, Liao C-Y, Yang W-Y, Lin Y-M, Jin S-LC, Wang H-D, Yuh C-H. 2014. Overexpression of Endothelin 1 Triggers Hepatocarcinogenesis in Zebrafish and Promotes Cell Proliferation and Migration through the AKT Pathway. PLOS ONE 9:e85318.

36. Nihei S, Asaka J, Takahashi H, Kudo K. 2021. Bevacizumab Increases Endothelin -1 Production via Forkhead Box Protein O1 in Human Glomerular Microvascular Endothelial Cells In Vitro. Int J Nephrol 2021:8381115.

37. Renga B, Cipriani S, Carino A, Simonetti M, Zampella A, Fiorucci S. 2015. Reversal of Endothelial Dysfunction by GPBAR1 Agonism in Portal Hypertension Involves a AKT/FOXOA1 Dependent Regulation of H2S Generation and Endothelin-1. PLOS ONE 10:e0141082.

38. Chen Q, Edvinsson L, Xu C-B. 2009. Role of ERK/MAPK in endothelin receptor signaling in human aortic smooth muscle cells. BMC Cell Biol 10:52.

39. Foschi M, Chari S, Dunn MJ, Sorokin A. 1997. Biphasic activation of p21ras by endothelin-1 sequentially activates the ERK cascade and phosphatidylinositol 3-kinase. EMBO J 16:6439–6451.

40. Ersoy Y, Bayraktar N, Mizrak B, Ozerol I, Gunal S, Aladag M, Bayindir Y. 2006. The level of endothelin-1 and nitric oxide in patients with chronic viral hepatitis B and C and correlation with histopathological grading and staging. Hepatol Res Off J Jpn Soc Hepatol 34:111–6.

41. FlyBase Gene Report: Dmel\CG43775. http://flybase.org/reports/FBgn0264297. Retrieved 20 October 2022.

42. Regn M, Laggerbauer B, Jentzsch C, Ramanujam D, Ahles A, Sichler S, Calzada-Wack J, Koenen RR, Braun A, Nieswandt B, Engelhardt S. 2016. Peptidase inhibitor 16 is a membrane-tethered regulator of chemerin processing in the myocardium. J Mol Cell Cardiol 99:57–64.

43. Hope CM, Welch J, Mohandas A, Pederson S, Hill D, Gundsambuu B, Eastaff-Leung N, Grosse R, Bresatz S, Ang G, Papademetrios M, Zola H, Duhen T, Campbell D, Brown CY, Krumbiegel D, Sadlon T, Couper JJ, Barry SC. 2019. Peptidase inhibitor 16 identifies a human regulatory T-cell subset with reduced FOXP3 expression over the first year of recent onset type 1 diabetes. Eur J Immunol 49:1235–1250.

44. Ferland DJ, Mullick AE, Watts SW. 2020. Chemerin as a Driver of Hypertension: A Consideration. Am J Hypertens 33:975–986.

45. Deng M, Yang S, Ji Y, Lu Y, Qiu M, Sheng Y, Sun W, Kong X. 2020. Overexpression of peptidase inhibitor 16 attenuates angiotensin II–induced cardiac fibrosis via regulating HDAC1 of cardiac fibroblasts. J Cell Mol Med 24:5249–5259.

46. Sarafidis PA, Bakris GL. 2007. Insulin and Endothelin: An Interplay Contributing to Hypertension Development? J Clin Endocrinol Metab 92:379–385.

47. Bellen HJ, Levis RW, He Y, Carlson JW, Evans-Holm M, Bae E, Kim J, Metaxakis A, Savakis C, Schulze KL, Hoskins RA, Spradling AC. 2011. The Drosophila Gene Disruption Project: Progress Using Transposons With Distinctive Site Specificities. Genetics 188:731–743.

48. Metaxakis A, Oehler S, Klinakis A, Savakis C. 2005. Minos as a Genetic and Genomic Tool in Drosophila melanogaster. Genetics 171:571–581.

49. Ohno M, Sekiya T, Nomura N, Daito T ji, Shingai M, Kida H. 2020. Influenza virus infection affects insulin signaling, fatty acid-metabolizing enzyme expressions, and the tricarboxylic acid cycle in mice. Sci Rep 10:10879.

50. Xu J, Hopkins K, Sabin L, Yasunaga A, Subramanian H, Lamborn I, Gordesky-Gold B, Cherry S. 2013. ERK signaling couples nutrient status to antiviral defense in the insect gut. Proc Natl Acad Sci 110:15025–15030.

51. DiAngelo JR, Bland ML, Bambina S, Cherry S, Birnbaum MJ. 2009. The immune response attenuates growth and nutrient storage in Drosophila by reducing insulin signaling. Proc Natl Acad Sci 106:20853–20858.

52. Sansone CL, Cohen J, Yasunaga A, Xu J, Osborn G, Subramanian H, Gold B, Buchon N, Cherry S. 2015. Microbiota-dependent priming of antiviral intestinal immunity in Drosophila. Cell Host Microbe 18:571–581.

53. Diamond MS, Roberts TG, Edgil D, Lu B, Ernst J, Harris E. 2000. Modulation of Dengue Virus Infection in Human Cells by Alpha, Beta, and Gamma Interferons. J Virol 74:4957–4966.

54. Keller BC, Fredericksen BL, Samuel MA, Mock RE, Mason PW, Diamond MS, Gale M. 2006. Resistance to Alpha/Beta Interferon Is a Determinant of West Nile Virus Replication Fitness and Virulence. J Virol 80:9424–9434.

55. Laurent-Rolle M, Boer EF, Lubick KJ, Wolfinbarger JB, Carmody AB, Rockx B, Liu W, Ashour J, Shupert WL, Holbrook MR, Barrett AD, Mason PW, Bloom ME, García-Sastre A, Khromykh AA, Best SM. 2010. The NS5 Protein of the Virulent West Nile Virus NY99 Strain Is a Potent Antagonist of Type I Interferon-Mediated JAK-STAT Signaling. J Virol 84:3503–3515.

56. Lazear HM, Daniels BP, Pinto AK, Huang AC, Vick SC, Doyle SE, Gale M, Klein RS, Diamond MS. 2015. Interferon-λ restricts West Nile virus neuroinvasion by tightening the blood-brain barrier. Sci Transl Med 7:284ra59–284ra59.

57. Lazear HM, Pinto AK, Vogt MR, Gale M, Diamond MS. 2011. Beta Interferon Controls West Nile Virus Infection and Pathogenesis in Mice. J Virol 85:7186–7194.

58. Samuel MA, Diamond MS. 2005. Alpha/beta interferon protects against lethal West Nile virus infection by restricting cellular tropism and enhancing neuronal survival. J Virol 79:13350–13361.

59. Shives KD, Beatman EL, Chamanian M, O’Brien C, Hobson-Peters J, Beckham JD. 2014. West Nile Virus-Induced Activation of Mammalian Target of Rapamycin Complex 1 Supports Viral Growth and Viral Protein Expression. J Virol 88:9458– 9471.

60. Li Q, Zhang Y-Y, Chiu S, Hu Z, Lan K-H, Cha H, Sodroski C, Zhang F, Hsu C-S, Thomas E, Liang TJ. 2014. Integrative Functional Genomics of Hepatitis C Virus Infection Identifies Host Dependencies in Complete Viral Replication Cycle. PLOS Pathog 10:e1004163.

61. Wang L, Yang L, Fikrig E, Wang P. 2017. An essential role of PI3K in the control of West Nile virus infection. Sci Rep 7:3724.

62. Wang S, Xia P, Huang G, Zhu P, Liu J, Ye B, Du Y, Fan Z. 2016. FoxO1-mediated autophagy is required for NK cell development and innate immunity. Nat Commun 7:11023.

63. Daffis S, Lazear HM, Liu WJ, Audsley M, Engle M, Khromykh AA, Diamond MS. 2011. The Naturally Attenuated Kunjin Strain of West Nile Virus Shows Enhanced Sensitivity to the Host Type I Interferon Response▿. J Virol 85:5664–5668.

64. Frias MA, Montessuit C. 2013. JAK-STAT signaling and myocardial glucose metabolism. JAK-STAT 2:e26458.

65. Gual P, Baron VR, Lequoy VR, Obberghen EV. 1998. Interaction of Janus Kinases JAK-1 and JAK-2 with the Insulin Receptor and the Insulin-Like Growth Factor-1 Receptor 139:10.

66. Hahn MB, Monaghan AJ, Hayden MH, Eisen RJ, Delorey MJ, Lindsey NP, Nasci RS, Fischer M. 2015. Meteorological Conditions Associated with Increased Incidence of West Nile Virus Disease in the United States, 2004–2012. Am J Trop Med Hyg 92:1013–1022.

67. Holcomb K. 2022. Worst-ever U.S. West Nile virus outbreak potentially linked to a wetter-than-average 2021 Southwest monsoon | NOAA Climate.gov. Climate.gov.

68. Saiz J-C. 2020. Animal and Human Vaccines against West Nile Virus. 12. Pathogens 9:1073.

69. Elbadry MM, Tharwat M, Mohammad EF, Abdo EF. 2020. Diagnostic accuracy of serum endothelin-1 in patients with HCC on top of liver cirrhosis. Egypt Liver J 10:19.

70. Shi L, Zhou S-S, Chen W-B, Xu L. 2017. Functions of endothelin-1 in apoptosis and migration in hepatocellular carcinoma. Exp Ther Med 13:3116–3122.

71. Kalani M. 2008. The importance of endothelin-1 for microvascular dysfunction in diabetes. Vasc Health Risk Manag 4:1061–1068.

72. Lenoir O, Milon M, Virsolvy A, Hénique C, Schmitt A, Massé J-M, Kotelevtsev Y, Yanagisawa M, Webb DJ, Richard S, Tharaux P-L. 2014. Direct Action of Endothelin-1 on Podocytes Promotes Diabetic Glomerulosclerosis. J Am Soc Nephrol 25:1050–1062.

73. Shives KD, Massey AR, May NA, Morrison TE, Beckham JD. 2016. 4EBP-Dependent Signaling Supports West Nile Virus Growth and Protein Expression. Viruses 8:287.

74. Harsh S, Ozakman Y, Kitchen SM, Paquin-Proulx D, Nixon DF, Eleftherianos I. 2018. Dicer-2 Regulates Resistance and Maintains Homeostasis against Zika Virus Infection in Drosophila. J Immunol 201:3058–3072.

75. Jin Y-H, Kang B, Kang HS, Koh C-S, Kim BS. 2020. Endothelin-1 contributes to the development of virus-induced demyelinating disease. J Neuroinflammation 17:307.

76. Adams KL, Riparini G, Banerjee P, Breur M, Bugiani M, Gallo V. 2020. Endothelin-1 signaling maintains glial progenitor proliferation in the postnatal subventricular zone. 1. Nat Commun 11:2138.

77. Koyama Y. 2013. Endothelin systems in the brain: involvement in pathophysiological responses of damaged nerve tissues. Biomol Concepts 4:335– 347.

78. Swire M, Kotelevtsev Y, Webb DJ, Lyons DA, ffrench-Constant C. 2019. Endothelin signalling mediates experience-dependent myelination in the CNS. eLife 8:e49493.

79. Sejvar JJ. 2014. Clinical Manifestations and Outcomes of West Nile Virus Infection. Viruses 6:606–623.

80. Briese T, Jia X-Y, Huang C, Grady LJ, Lan Lipkin W. 1999. Identification of a Kunjin/West Nile-like flavivirus in brains of patients with New York encephalitis. The Lancet 354:1261–1262.

81. Ahlers LRH, Bastos RG, Hiroyasu A, Goodman AG. 2016. Invertebrate Iridescent Virus 6, a DNA Virus, Stimulates a Mammalian Innate Immune Response through RIG-I-Like Receptors. PLOS ONE 11:e0166088.

82. Zhang W, Thompson BJ, Hietakangas V, Cohen SM. 2011. MAPK/ERK Signaling Regulates Insulin Sensitivity to Control Glucose Metabolism in Drosophila. PLOS Genet 7:e1002429.

83. Goodman AG, Fornek JL, Medigeshi GR, Perrone LA, Peng X, Dyer MD, Proll SC, Knoblaugh SE, Carter VS, Korth MJ, Nelson JA, Tumpey TM, Katze MG. 2009. P58(IPK): a novel “CIHD” member of the host innate defense response against pathogenic virus infection. PLoS Pathog 5:e1000438.

84. Subramanian A, Tamayo P, Mootha VK, Mukherjee S, Ebert BL, Gillette MA, Paulovich A, Pomeroy SL, Golub TR, Lander ES, Mesirov JP. 2005. Gene set enrichment analysis: A knowledge-based approach for interpreting genome-wide expression profiles. Proc Natl Acad Sci 102:15545–15550.

85. Martin M, Hiroyasu A, Guzman RM, Roberts SA, Goodman AG. 2018. Analysis of Drosophila STING Reveals an Evolutionarily Conserved Antimicrobial Function. Cell Rep 23:3537–3550.e6.

86. Spellberg MJ, Marr MT. 2015. FOXO regulates RNA interference in Drosophila and protects from RNA virus infection. Proc Natl Acad Sci U S A 112:14587–14592.

87. Torres MJ, López-Moncada F, Herrera D, Indo S, Lefian A, Llanos P, Tapia J, Castellón EA, Contreras HR. 2021. Endothelin-1 induces changes in the expression levels of steroidogenic enzymes and increases androgen receptor and testosterone production in the PC3 prostate cancer cell line. Oncol Rep 46:171.

